# Dynamic programming algorithms for fast and accurate cell lineage tree reconstruction from CRISPR-based lineage tracing data

**DOI:** 10.1101/2024.11.15.623872

**Authors:** Junyan Dai, Erin K. Molloy

**Affiliations:** Department of Computer Science, University of Maryland, College Park, 20742, USA; University of Maryland Institute for Advanced Computer Studies, University of Maryland, College Park, 20742, USA

**Keywords:** lineage tracing, CRISPR, phylogenetics, parsimony, Star Homoplasy, Camin-Sokal

## Abstract

CRISPR-based lineage tracing, coupled with single-cell RNA sequencing, has emerged as a promising approach for studying cell transformations during development as well as disease progression. However, the high ratio of cells to CRISPR-induced mutations, combined with missing data from silencing or dropout, make cell lineage tree (CLT) reconstruction difficult. As a result, this computational problem has attracted significant attention in recent years, including the introduction of Star Homoplasy Parsimony (SHP) in 2023 to model the specific properties of CRISPR-induced mutations, along with the Startle family of methods based on integer linear programming (ILP) or heuristic search (NNI). Here, we present Star-CDP, the first dynamic programming algorithm for SHP. Star-CDP solves SHP within a constrained search space Σ defined by subsets of cells from which a solution CLT must draw its clades. When Σ is the power set, Star-CDP is an exact exponential algorithm with time complexity *O*(*nm* |Σ| ^2^), where *n* is the number of cells, *m* is the number of target sites, and |Σ |= *O*(2^*n*^). We show that it is possible to build clade constraints that are polynomially-sized and effective in practice. Motivated by the technological challenges in producing consistent phylogenetic signal across the tree during lineage tracing, we also present algorithms to efficiently count, sample, and build consensus trees from all solutions to the clade-constrained SHP problem. In simulations, Star-CDP’s strict consensus effectively reduced false positive branches while preserving many more true positives compared to the standard strict consensus implemented by PAUP*, a popular parsimony method from species phylogenetics. Likewise, Star-CDP’s strict consensus achieved the same or higher accuracy (f1-score) on all but one of the 15 model conditions tested, often outperforming leading the methods, Startle-ILP and Startle-NNI, while also scaling to larger data sets than Startle-ILP. Lastly, we analyzed lineage tracing data from the KP-Tracer mouse model of lung adenocarcinoma, finding that Star-CDP produced plausible CLTs, often lowering the number of migration and reseeding events needed to explain metastases compared to Startle. Our analysis also showed, for the first time, that strategies for preprocessing cells with missing data—specifically cell pruning and deduplicating techniques—can have a substantial impact on CLTs reconstructed with the same method, even changing relative performance across methods compared to previously published results. The same was true of postprocessing trees with LAML, a maximum likelihood method designed for mixed-type missing data. By exploring these different pipelines, we recovered the most plausible CLT for the largest KP-Tracer metastatic tumor, reducing the number of reseeding events from 42 to 10 without increasing the number of migrations. Star-CDP is available on Github: https://github.com/molloy-lab/Star-CDP.

## 1 Introduction

For humans and many multi-cellular organisms, each cell in the body traces its origin back to a single fertilized egg through a sequence of cell divisions. During this branching process, cells differentiate into various types and organize into complex tissues and organs. In some cases, cells acquire the ability to bypass regulatory mechanisms that control growth, death, and movement, leading to diseases such as cancer [21]. **Cell lineage trees (CLTs)** depict the history of cell divisions and provide important context for studying fundamental biological processes. However, CLTs cannot be directly observed under most conditions. Instead, they are typically reconstructed from mutation data and then used in downstream analyses, including the inference of cell differentiation maps [8, 22, 31] and gene expression dynamics [36]. In the context of cancer, CLTs are useful for identifying subclones within a tumor [39], testing for adaptive responses of subclones to therapeutics [11], and inferring migration histories for metastatic tumors [7]. Thus, accessible technologies for reconstructing high-resolution CLTs have the potential to transform research in cancer and developmental biology.

A major challenge is that the somatic mutation rate is low in many important contexts, such as embryo development [17]. In the case where there are ample somatic mutations across the genomes of different cells, it can be difficult to accurately and comprehensively identify these mutations, even using state-of-the-art single-cell sequencing technologies [17]. Recent advances in genome editing offer a promising path forward: the idea is to use CRISPR Cas systems to induce DNA cleavage at target sites inserted into the genome [14]. The subsequent DNA repair introduces heritable mutations at the target site, which can be effectively captured via single-cell RNA sequencing. Because target sites cannot be cleaved more than once, each CRISPR-induced mutation is inherited by all daughter cells; however, DNA repair can produce similar mutations across the tree, leading to parallel mutations or **homoplasy** at target sites. Put simply, CRISPR-induced mutations are not expected to map perfectly to the CLT, challenging CLT reconstruction.

Several lineage tracers of this form have been developed, including scGESTALT [23, 24], ScarTrace [1], LINNAEUS [32], and KPTracer [39]. Simultaneously, CLT reconstruction has attracted significant attention from the method developers. The DREAM challenge in 2021 [9] resulted in the popular method Cassiopeia [12], which is based on integer linear programming (ILP) but also implements greedy and hybrid modes for greater scalability. The leading method to date is perhaps Startle [29, 30]. Startle employs a mixed ILP to find an optimal solution for the **Star Homoplasy Parsimony (SHP)** problem. SHP is equivalent to Camin-Sokal parsimony (CKP) [3] with a restricted substitution matrix so that only mutations from the unedited state to edited states are allowed, thus modeling the *non-modifiability* property of CRISPR-based lineage tracers. Because ILP can be prohibitively expensive on large data sets [29], Startle also implements hill climbing based on nearest neighbor interchange (NNI) tree edit moves. Although Startle-NNI is more scalable than Startle-ILP, it has no optimality guarantee. Moreover, heuristic search is challenged by low information content and missing data, which lead to phylogenetic terraces of many equally optimal trees [27]. Low information content results from the small number of target sites as well as their potential for early saturation, which in turn decreases the mutation rate over time [26]. Missing data results from silencing at target sites, producing *heritable missing data*, as well as dropout from single-cell sequencing, producing *random missing data* [12, 15].

To address these challenges, we present **StarCDP**: a dynamic programming (DP) algorithm that that solves the SHP problem within a constrained search space Σ defined by subsets of cells from which a solution CLT must draw its clades. We show that the clade-constrained SHP problem can be solved in *O*(*nm* |Σ| ^1.726^ +*n* |Σ| ^2^) time, where *n* is the number of cells and *m* is the number of target sites. When Σ is the power set, Star-CDP is exact exponential algorithm. We show how that it is possible to build clade constraints that are polynomially-sized and effective in practice, leveraging techniques from species phylogenetics. Clade constraints were introduced for gene tree parsimony problems by Hallet and Lagergren in 2000 [10]. Since then, this technique has been widely employed to optimization problems from species phylogenetics with much success [34, 35, 20, 4, 5], most notably the popular species tree method ASTRAL [18, 19, 40]. Notably, the solutions to these prior clade-constrained problems are binary trees. This assumption is appropriate for CLTS, as each cell division event produces two daughter cells; however, the limited information content in lineage tracing data means that many edges in the tree will not be supported by mutations. Unfortunately, we show that requiring solutions to contain only non-terminal branches with one or more mutation escalates the problem to NP-hard, despite the use of clade constraints. This motivated us to develop algorithms for counting, sampling, and building consensus trees from all solutions to the clade-constrained SHP problem.

In simulations, we found that Star-CDP’s strict consensus effectively reduced the number of false positives, while preserving many true positives compared to the standard strict consensus, implemented within PAUP*, a popular parsimony method from species phylogenetics. Moreover, Star-CDP’s strict consensus achieved the same or higher accuracy (f1-score) than the Startle and the other methods tested. This trend held across the model conditions tested, regardless of whether mutationless branches were contracted before computing accuracy, except when the proportion of heritable missing data (versus random missing) reached 100 percent; in which case, LAML, a maximum likelihood method designed for mixed-type missing data, performed best [15, 16]. Importantly, these trends were not readily observable when method performance was evaluated using the popular Robinson-Foulds distance [25], the only tree error/accuracy metric reported in several prior studies (e.g., those presenting the Startle [29] and LAML [15] methods).

Lastly, we analyzed lineage tracing data from the KP-Tracer mouse model of lung adenocarcinoma. We found that Star-CDP was as effective as Startle-NNI at finding plausible CLTs, even lowering the number of migration and reseeding events needed to explain metastases. Our reanalysis also shows, for the first time, that strategies for preprocessing cells with missing data—specifically cell pruning and deduplicating techniques—can have a substantial impact on CLTs reconstructed with the same method, even changing relative performance across methods. The same was true of postprocessing trees with LAML. By exploring these different pipelines, we built the most plausible history of a large KP-Tracer tumor to date, reducing the number of reseeding events from 42 to 10, without increasing the number of migrations.

The remainder of this paper is organized as follows. We review the SHP problem in Section 2. Section 2 introduces the clade-constrained version of this problem and then presents the Star-CDP method and related theoretical results. Section 4 describes how methods are evaluated on synthetic data and the results of this simulation study. Our reanalysis of KP-Tracer tumors is presented in Section 5. We conclude in Section 6 with recommendations for data processing and benchmarking methods and future directions from our study.

## 2 Background

To present Star-CDP, we briefly review definitions for phylogenetic trees as well Star Homoplasy parsimony.

### Phylogenetic Trees

A cell lineage tree is rooted phylogenetic *T* : a directed acyclic graph whose leaves are bijectively labeled by elements of a set *S*, representing cells. The unique vertex with in-degree zero is called the root, vertices with out-degree zero are called leaves, and vertices with out-degree greater than two are called *polytomies*. A tree *T* is *binary* or fully resolved if all non-leaf vertices (called internal vertices) have out-degree two; otherwise *T* is *non-binary*. We use *r*(*T*), *L*(*T*), *V* (*T*), and *E*(*T*) to denote the root, leaf set, vertex set, and edge set of *T*, respectively. For simplicity of notation, we treat the leaf set *L*(*T*) and the label set *S* as being interchangeable. We now define clades and subtree bipartitions.

#### Definition 1 (Clade).

*The* clade *induced by vertex v* ∈ *V* (*T*), *denoted as Clade*(*v*), *is the set of leaves that are descendants of v. The clade encoding of T is Clade*(*T*) = *{Clade*(*v*) : *v* ∈ *V* (*T*)*}*.

#### Definition 2 (Subtree Bipartition).

*Let T be a binary CLT. Then, each internal vertex v has a set X of leaves that are descendants of its left child (denoted v*.*left) and a set Y of leaves that are descendants of its right child (denoted v*.*right) or vice versa. X and Y form the* subtree bipartition (STB) *induced by vertex v, denoted SubBip*(*v*) = *X*|*Y*. *The STB encoding of T is SubBip*(*T*) = *{SubBip*(*v*) : *v* ∈ *V* (*T*) | *L*(*T*)*}*.

### Characters and Star Homoplasy Parsimony (SHP)

A *character c* maps cells in *S* to the state set. In CRISPR-based lineage tracing, each character is a target site and each state represents the DNA sequence observed at that target site in some cell. The DNA sequences observed at each target site are coded as integers with zero (0) representing the unedited state, positive integers (1, 2, …) representing edited states, and negative one (-1) or question mark (?) representing missing or ambiguous states. We use 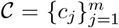 to denote a collection of characters from CRISPR-based lineage tracing, with *c*_*j*_[*s*] returning the state of cell *s* at target site *c*_*j*_. Given character *c*_*j*_ and CLT *T*, a *vertex labeling* for (*T*, ĉ_*j*_) maps the vertices of *T* to the state set for character *c*_*j*_, denoted 𝒜_*j*_, such that ĉ_*j*_[*l*] = *c*_*j*_[*l*] for all *l*∈ *L*(*T*). We say that a mutation occurs on edge *u* ⟼ *v*∈ *E*(*T*) if ĉ_*j*_[*v*] ≠ ĉ_*j*_[*u*], provided that neither ĉ_*j*_[*v*] nor ĉ_*j*_[*u*] are the missing state. Recall that only mutations from the unedited state to an edited state are allowed under **Star Homoplasy (SH) parsimony** [29].

#### Definition 3 (SH-labeling and SH-contraction).

*We say a vertex labeling* ĉ_*j*_ *for* (*T, c*_*j*_) *is a* Star Homoplasy (SH) labeling *if (1) it obeys the non-modifiability property and (2) it minimizes the number of mutations across T*. *Then, given T and a set 𝒞of characters, the* SH-contraction *of T can be created by computing the SH-labelings for all characters, denoted* 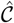, *and then contracting mutationless non-terminal branches (e*.*g*., *see Fig. 4)*.

Importantly, some mutations are more common than others due to DNA repair mechanisms. The probability of each mutation can be estimated from data and then used to determine a mutation cost function: *w*_*j*_ : *A*_*j*_ \{0, -1} ↦ ℝ_>0_. The estimated mutation cost function is then incorporated into the parsimony score.

#### Definition 4 (SH Parsimony (SHP) Score).

*Given rooted tree T and character c*_*j*_, *both on the same cell set S, as well as mutation cost function w*_*j*_, *the* SHP score *is*

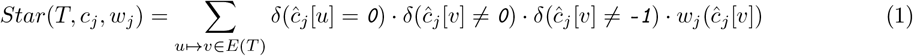

*where* Δ *is the indicator function and* ĉ_*j*_ *is an SH-labeling for* (*T, c*_*j*_).

SHP score can be computed in polynomial time via Sankoff’s algorithm adapted for the SH-labeling [28, 29].

### Problem 1 (Large SH parsimony (LSHP) Problem)

The *LSHP problem* takes as input a set 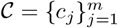 of characters, each on cell set *S*, and a set of 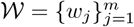 of mutation cost functions. The solution is a rooted *binary* tree *T* on cell set *S* that minimizes 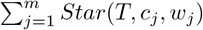.

SHP is equivalent to the classic *Camin-Sokal parsimony* problem [3] when the state set is binary (i.e., {-1, 0, 1}) and the mutation cost function is 1 for all characters *j*∈ {1, 2, …, *m*}. The large Camin-Sokal parsimony problem is NP-hard [6]; thus, it is easy to see that the large SHP problem is NP-hard.

## 3 Methods

### 3.1 Clade-constrained large SHP problem and the Star-CDP method

We now introduce the clade-constrained large SHP problem and the Star-CDP method.

#### Problem 2 (Clade-constrained (CC) LSHP problem)

The *CC-LSHP problem* takes as input: a set 𝒞 of characters, each on cell set *S*, the mutation cost functions 𝒲, and a set Σ of clades (i.e., subsets of cell set *S*). The input size is described in terms *n* = |*S*|, *m* = |*𝒞*|, and *q* = |Σ|. The solution is a rooted *binary* tree *T* on cell set *S* such that 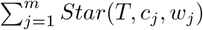 is minimized and *Clade*(*T*) ⊆ Σ if such a tree exists. A solution to CC-LSHP exists provided there is at least one rooted binary tree *T* on *S* satisfying *Clade*(*T*) ⊆ Σ.

Our dynamic programming algorithms for CC-LSHP, called Star-CDP, follow from our observation that the clade set Σ not only constrains the search space in terms of the allowed tree topologies but also the allowed state assignments (i.e., SH-labelings) at the internal vertices.

##### Theorem 1.

*Let T be a rooted tree and let c be a character, both on cell set S. Assume that there is no missing data so c does not map to the ambiguous state. Then, for any vertex v* ∈*V* (*T*), *the SH-labeling* ĉ[*v*] *is unique and only depends on Clade*(*v*); *no other information about T is needed*.

The proof of Theorem 1 appears in the Supplement. In the presence of missing data, the SH-labeling exists but it may not be unique. However, it is possible to compute a unique SH-labeling ĉ_*_ for (*T, c*) such that ĉ_*_[*v*] depends only the clade induced by *v* (Theorem 2 in the Supplement). In brief, state 0 must be assigned anytime there are two or more non-missing states in an induced clade; otherwise the only non-missing state in the clade must be assigned (Fig. 1). This is summarized as the function `GetStates`(*A*, 𝒞), which returns the SH-labelings of the vertex inducing clade *A* given character set 𝒞 in *O* (*nm*) time (Algorithm 1 in the Supplement). If the SH-labelings for some vertex *v* and its two children are known, the cost of mutations on the two outgoing edges from *v* can be computed in just *O* (*m*) time. Putting this all together, we can compute the SHP cost of mutations on the two edges associated with subtree bipartition *X*| *Y* (with *A* = *X*∪ *Y*), denoted `SHPCost`(*A, X, Y*, 𝒞, 𝒲), in *O* (*nm*) time (Algorithm 2 in the Supplement). Before presenting the dynamic programming (DP) algorithm for CC-LSHP, we need some final notation. We let 𝒮 𝒯 ℬ= {*X*| *Y* : *X, Y, X* ∪*Y* ∈ Σ and *X*∩ *Y* =∅} denote the subtree bipartitions that are allowed to appear in solutions based on the constraint set Σ. Likewise, we let 𝒮 𝒯 ℬ (*A*) = *X Y* : *X Y* = *A* be the subtree bipartitions in 𝒮 𝒯 ℬassociated with clade *A*, which can be computed for *A* on the fly or as part of preprocessing (Algorithm 3 in Supplement).

**Fig. 1.**
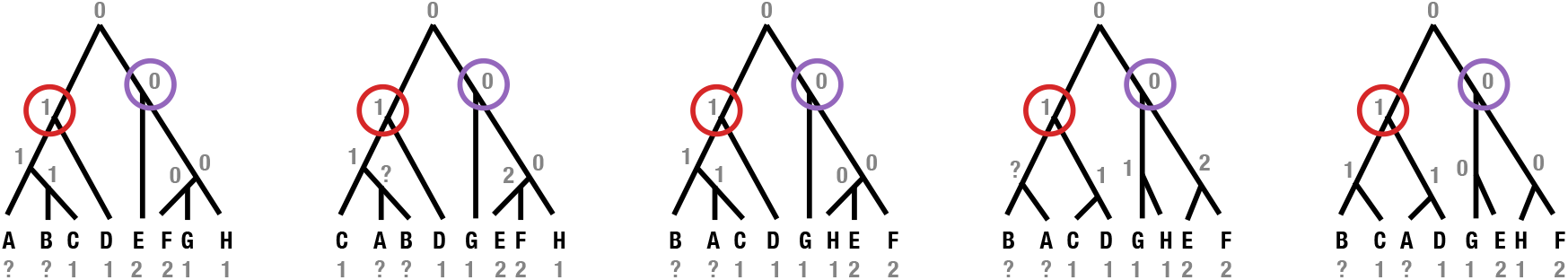
Constrained SH-labeling for clades. We show five trees on leaf set *A*-*H*, all of which contain clades *A*− *D* and *E*− *H*. For the character shown at the leaves, the SH-labeling at the verteices inducing clades *A* − *D* and *E* − *H* does NOT change even when the tree topologies below these clades changes. Note that ? denotes the missing or ambiguous state in this example instead of -1.

#### DP algorithm for CC-LSHP (Star-CDP)

Let *Star*[*A*] be the SHP cost of a solution to the CC-LSHP for clade *A*. Then, the following DP algorithm solves CC-LSHP.

**Base Case:** Clade *A* contains a single cell. *Star*[*A*] := 0

**Recurrence:** Clade *A* contains multiple cells.

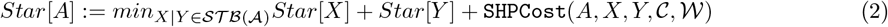

The SHP score for any solution to CC-LSHP equals *Star*[*S*]. An arbitrary solution tree can be recovered by backtracking. Star-CDP has time complexity *O*(*nm* |Σ| ^1.726^ + *n* |Σ| ^2^) and space complexity *O*(|Σ|) excluding the inputs (Theorem 5 in the Supplement; also see Theorem 4 in the Supplement for proof of correctness).

#### Clade constraints

When Σ is the power set, Star-CDP is an exact exponential algorithm for LSHP. In practice, our goal is to construct Σ from the data so that (1) good solutions to LSHP lie with the constrained search space and (2) the constrained search space is small enough for Star-CDP to be fast. Effective techniques for building Σ have been proposed as part of the popular method ASTRAL [18], which reconstructs species trees from a set of *k* input gene trees. ASTRAL builds Σ by adding any missing taxa into each gene tree, arbitrarily rooting each gene tree at the same leaf, and finally extracting all clades from each gene tree [19]. If the input gene trees are non-binary, ASTRAL-III extends the constrained search space refining polytomies in different ways but limiting the sampling to ensure the size of Σ is polynomial in the input [40]. Overall, this process results in |Σ| = *O*((*nk*)^1.726^*D*) where *D* is the total number of unique nodes in the input gene trees. If all nodes are unique, |Σ| = *O*((*nk*)^2.726^). Unlike ASTRAL, Star-CDP takes characters, rather than gene trees, as input. Thus, to apply ASTRAL’s techniques for building Σ, we first perform heuristic search under SHP, saving the *k* best trees found. These trees are then given to ASTRAL-III to build Σ, typically after SH-contraction.

### 3.2 Dealing with low phylogenetic informativeness and significance of Star-CDP compared to prior results

Star-CDP is that returns a single binary tree, like Startle-NNI, Cassiopeia-Greedy, and many other methods. Although the true CLT is binary, lineage tracing systems are unlikely to produce a mutation on each internal branch [12]. Moreover, if there are many mutationless non-terminal branches are adjacent in the CLT *T*, there will be many equally good solutions (i.e., binary trees with the same SHP score)—just consider all ways to refine polytomies in the SH-contraction of *T*. This motivates us to introduce the mutation-and-clade constrained large star homoplasy parsimony problem (MCC-LSHP), which is the same as CC-LSHP but additionally requires the output tree *T* to have at least one mutation on each of its non-terminal edges. Unfortunately, unlike the problem CC-LSHP, which can be solved in polynomial time, MCC-LSHP is NP-hard (Theorem 5 in the Supplement). Our proof is based on the proof that (unconstrained) large Camin-Sokal parsimony (LCKP) is NP-complete by Day, Johnson, and Sankoff [6]. Their proof gives a Karp reduction from maximum independent set (MIS) for 3-regular graphs to LCKP. It is easy to see that the solution CLTs will be non-binary. Our proof explicitly encodes the restrictions on the input graph for MIS as mutation and clade constraints for LCKP (note that the number of clade constraints for LCKP is linear in the number of vertices in the input graph for MIS). Intuitively, MIS is NP-complete even for highly restricted (3-regular) graphs and LCKP is NP-hard even with mutation and clade constraints. To our knowledge, the hardness of LCKP when solutions must be binary was unknown, making our exponential algorithm more interesting. Additionally, our exponential algorithm for LSHP becomes polynomial time when there are polynomially many clade constraints, unlike the mutation and clade constrained version of the problem.

#### Counting and building consensus trees from solutions to CC-LSHP

A standard approach to addressing low phylogenetic signal in cell lineage tracing data is to take the SH-contraction of trees before computing tree error [29]. However, this process will not remove false positive edges on which mutations occur, motivating us to develop algorithms to build consensus trees from all solutions to CC-LSHP. The key computational problem is counting the number of solutions to CC-LSHP that contain each clade *A* ∈Σ. Dividing by total number of solutions gives the clade frequency *Freq*[*A*]. This quantity can be used to select clades when building strict, majority, and greedy consensus trees.

To compute *Freq*[*A*], we introduce two related problems and DP algorithms, with solutions denoted by *Below*[*A*] and *Above*[*A*]. *Below*[*A*] is the number of solutions “below” clade *A* (i.e., it is the number of binary trees on cell set *A* that draw their clades from Σ and achieve score *Star*[*A*]). *Above*[*A*] is the number of solutions “above” clade *A*, which we define procedurally:

1. Initialize 𝒢_*A*_ to the solution set for CC-LSHP.
2. Remove every tree from 𝒢_*A*_ that does not contain clade *A*.
3. For each tree *t*∈𝒢 _*A*_: 3a. Find vertex *v* that induces clade *A*. 3b. Delete the two outgoing edges from *v* as well as the subtrees below them.

The number of (unique) elements remaining in 𝒢_*A*_ is *Above*[*A*].

Now we can formulate any solution to CC-LSHP that contains clade *A* by taking any tree counted in *Below*[*A*] and attaching its root to the only unlabeled vertex in any tree counted in *Above*[*A*]. Putting this together, the fraction of solutions to CC-LSHP that contain clade *A* is *Freq*[*A*] = (*Below*[*A*] ·*Above*[*A*])*/Below*[*S*], where *Below*[*S*] is the total number of solutions for CC-LSHP, which equals *Above*[*s*] for any leaf *s*∈ *S*.

Our DP algorithms for computing *Above* and *Below* make use of two new data structures. The first structure *I* is indexed by a clade *A* and returns a list of pairs of pointers to the subproblems that solve *Star*[*A*] (i.e., we store all pairs of pointers that yield a solution to *Star*[*A*] instead of just storing one pair of pointers for backtracking). The second structure *J* reverses the mapping stored in *I* so that we can find parent subproblems. Specifically, for each (*X, Y*) ∈ *I*[*X*∪ *Y*], we create *J* [*X*] = (*X*∪ *Y, Y*) and *J* [*Y*] = (*X*∪ *Y, X*). It is easy to incorporate building *I* and *J* into the DP for CC-LSHP (Star-CDP; Algorithm 3).

#### DP algorithm for Below

Let *Below*[*A*] be the number of solutions to CC-LSHP containing clade *A* (see above). Then, the following DP algorithm computes *Below*[*A*].

**Base Case:** Clade *A* contains a single cell. *Below*[*A*] := 1 since there is just the one trivial solution.

**Recurrence:** Clade *A* contains multiple cells.

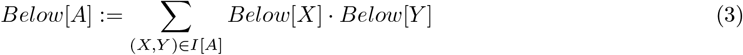

where *I*[*A*] stores pointers to the pairs of child subproblems when combined yield solutions to CC-LSHP for clade *A*. We have already argued the correctness of recurrence (see Theorem 6 in the Supplement for the remainder of the proof). The computation of *Below* can be integrated into the DP for CC-LSHP (StarCDP; Algorithm 3) without increasing time or storage complexity because *I* does not need to be maintained across subproblems. This enables us to efficiently count solutions to CC-LSHP. Now we turn to the computation for *Above*.

#### DP algorithm for Above

Let *Above*[*X*] be the number of solutions to CC-LSHP “above” clade *X*. Then, the following DP algorithm computes *Above*[*X*].

**Base Case:** Clade *X* is the full set *S* of cells. *Above*[*X*] := 1 since there is only the one solution containing clade *S* (nothing is above).

**Recurrence:** If clade *X* is a proper subset of *S*, the calculation is more complicated. Consider a solution to CC-LSHP that contains both clade *X* and clade *A* so (*X, Y*) ∈ *I*[*A*]. Such a solution can be formed by taking a tree counted in *Below*[*X*], joining it at the root to a tree counted in *Below*[*Y*], and then attaching the new root to the unlabeled vertex of a tree counted in *Above*[*A*]. The number of ways to build such a tree is *Below*[*X*] ·*Below*[*Y*] ·*Above*[*A*]. However, we are only interested in counting the solutions above *X* that also contain clade *A*, giving the recurrence:

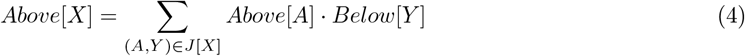

where *J* [*X*] stores pointers to pairs of parent *A* and child *Y* subproblems s.t. combining *Y* with *X* yields solutions to CC-LSHP for clade *A*.

We have already argued the correctness of recurrence for *Above* (see Theorem 6 in the Supplement for the remainder of the correctness proof). The computation of *Above* requires *J* to be stored during the DP for CC-LSHP (Star-CDP; Algorithm 3), increasing the storage complexity from *O*(|*Sigma*|) *O*(|Σ| ^1.726^) (see Theorem 7 in the Supplement; also [13]). Given *J*, the time complexity of *Above* is the same as *Below*. After computing *Below* and *Above, Freq*[*A*] can be computed for any clade *A* ∈Σ in constant time.

Thus, the computation of clade frequencies *Freq* (as well as the related structures *I, J, Above*, and *Below*) does not increase the time complexity of Star-CDP but it does increase the storage complexity from *O*(|Σ|) to *O*(|Σ|^1.726^). After computing *Freq*, it is trivial to build consensus trees [37]. The **strict consensus (SC)** tree contains only clades *A* ∈ Σ such that *Freq*[*A*] = 1.

### Sampling solutions to CC-LSHP

The storage of *I* enables us to sample solutions to CC-LSHP (note that storing *I* in addition to *J* does not impact space complexity). Starting at the root (clade *S*), we select an element (subtree bipartition) from *I*[*S*] arbitrarily, yielding two child subproblems. Continuing in this fashion for child subproblems produces a **random** solution to CC-LSHP. We can also generate **biased** solutions to CC-LSHP, for example by selecting the most imbalanced subtree bipartition in *I*, breaking ties arbitrarily.

### Significance of our DP algorithms for building consensus trees compared to prior results

Algorithms for counting solutions in a constrained search space and building consensus trees from them have been described Vachaspati and Warnow [35]; however, they introduce algorithms *unrooted* trees, which simplifies the problem (only the *Below* matrix is needed). We introduce the first algorithms for *rooted* trees (which is why the *Above* matrix is needed). It is easy to see that these two counting problems are different because there are only 3 unrooted topologies on *S* = {1, 2, 3, 4}, exactly one of which contains bipartition 1, 2 |3, 4; however, there are 15 rooted topologies on *S*, 3 of which contain clade {1, 2} based on the three ways to join 3, 4 in the tree above clade {1, 2}. Our algorithms could be applied to other clade-constrained dynamic programming algorithms for rooted phylogenies, like those based on gene tree parsimony [2].

## 4 Simulation Study

### 4.1 Methods

We now describe an experimental study benchmarking Star-CDP against the leading methods on synthetic data sets where the true CLT is known. Details and software commands are in Section 2 of the Supplement. Scripts, along with links to data files, are available on Github: https://github.com/molloy-lab/star-study.

#### Startle simulated data sets

We analyzed synthetic data sets (with true CLTs and character matrices) from the Startle study [29]. These data sets were simulated using Cassiopeia [12] in three steps: (1) CLTs were generated with the Birth-Death Fitness Simulator, (2) data (mutations) were generated down the CLT using the Cas9 Lineage Tracing Data Simulator, and (3) dropout (missing values) were introduced (rate: 0.15). Twelve model conditions, each with 21 replicate data sets, were created by varying the number of cells (50, 100, 150, 200) and the number of characters/sites (10, 20, 30), yielding 251 data sets in total. The true CLTs available online already had mutationless branches contracted; thus, we only compare true and estimated trees after SH-contraction.

#### LAML simulated data sets

We also analyzed synthetic data sets from the LAML study [15]. These data sets (with 30 characters/sites) were simulated with a similar protocol to Startle, except that the in step (1) CLTs with 1024 cells were simulated under a birth only process with 10 generations, and in step (2) silencing was also modeled, resulting in heritable missing data. Finally, 250 cells (from the 1024) were sampled. Five model conditions were produced by varying the proportion of missing data from silencing versus dropout (0%, 25%, 50%, 75%, and 100%). Each of the 50 replicate data sets for each model condition encompassed 10 different mutations rates, yielding 250 data sets in total. The true CLTs available online were binary.

#### CLT reconstruction methods for CRISPR-based lineage tracing data

We primarily compared Star-CDP to *parsimony* methods for lineage tracing data, specifically the leading methods (Startle-ILP and Startle-NNI) as well as Cassiopeia-Greedy and PAUP* [33], a popular species phylogenetics package that, to our knowledge, has yet to be evaluated as a heuristic for SHP. ILP-based methods **Startle-ILP** and **Cassiopeia-Greedy** did not require special options. **Startle-NNI** was used to execute heuristic search under SHP using Nearest Neighbor Interchange (NNI) moves initiated from a user-provided starting tree; we built the starting tree with neighbor-joining (NJ) following the recommended commands (thus, the runtimes reported for Startle-NNI include time to compute the NJ tree). Importantly, there are two versions of Startle: the earlier Python version and the latest C++ version. We initially used the latest C++ version, but after we were unable to replicate results from the Startle study [29]; to address this issue, we also ran the Python version on the Startle simulated data sets. Like Startle-NNI, **PAUP\*** can be used to perform heuristic search under SHP. We implemented SHP by transforming multi-state characters into binary characters in the natural way and then executing the search under weighted Camin-Sokal parsimony (these two optimization problems are equivalent as described by [29]; see the Supplement for details). The heuristic search with PAUP* differed from Startle-NNI in that randomized taxon addition was used to produce a starting tree instead of NJ and Tree Bisection and Reconnection (TBR) edit moves were used instead of NNI edit moves. We ran PAUP* so that it saved 500 of the best-scoring trees found during its search, sorted from lowest to highest scores (notably, these trees were non-binary, likely related to SH-contraction). The first best-scoring tree saved is the PAUP* tree. The **PAUP*-SC** tree is the (standard) strict consensus of the 500 best trees. Our method Star-CDP requires users to provide trees so that clade constraints can be constructed with ASTRAL-III [40]; we gave Star-CDP the 500 best-scoring trees from PAUP* (thus the runtimes reported for StarCDP also include runtimes of ASTRAL and PAUP* unless otherwise noted). Star-CDP returns multiple trees: **StarCDP-Rand, StarCDP-Bias**, and **StarCDP-SC**, described in Section 2. Lastly, we ran **LAML**, a *maximization likelihood for mixed-type (i*.*e*., *heritable and random) missing data*. LAML uses heuristic search with NNI moves from a user-provided starting tree; we gave LAML the tree computed with Startle-NNI (C++) (thus, the runtimes reported for LAML include time to estimate the starting tree unless otherwise noted).

#### Tree error and accuracy

On simulated data sets, methods were evaluated in terms tree error and accuracy. We first computed the number of false positive (**FP**) branches (i.e., branches in the estimated tree not in the true tree), the number of false negative (**FN**) branches (i.e., branches in the true tree not in estimated tree), and the number of true positive (**TP**) branches (i.e., branches in both true and estimated trees). We then summed FN and FP to get the Robinson-Foulds (RF) error [25]. We also used these quantities to compute precision, recall, and f1-score. In the Startle study [29], tree error was computed *after* the SH-contraction was taken of both the true and estimated CLTs; we followed the same approach. In the LAML study [15], tree error was computed for binary trees *without* SH-contraction. To evaluate the impact of SH-contraction, we computed tree error and accuracy metrics both with and without SH-conraction of both the true and estimated CLTs.

#### Normalized SHP score

Methods were also evaluated in terms of normalization SHP score, defined as the SHP score of the estimated CLT divided by the SHP score of the true CLT.

#### Runtime

All CLT reconstruction methods were limited to 48 GB of RAM, 1 or 16 CPU (from AMD EPYC 7313 16-Core processor), and a maximum wallclock time of 24 hours. All methods were run with a single thread; except Startle-NNI (Python), which was run with 16 threads. We tested running Startle-ILP on the LAML data sets with 16 threads, but it still did not complete within the maximum wall clock time of 24 hours. When collecting running time data, methods were given exclusive access to a compute node, with resources managed by the SLURM submission system.

### 4.2 Results on Startle Simulated Data Sets

We now present the results of benchmarking methods for on the Startle simulated data sets, looking at tree error/accuracy, SHP score, and runtime.

#### Tree error/accuracy after SH-contraction

Across all model conditions, StarCDP-SC and PAUP*-SC achieved similar *RF error* on average, improving upon taking a single high-scoring tree and even outperforming the leading methods: Startle-ILP and Startle-NNI (Fig. 4.2; Figs. S2–S5 and Tables S2–S3 in the Supplement). Although StarCDP-SC and PAUP*-SC appeared to perform similarly in terms of RF score, they performed very differently in terms of FPs and TPs. PAUP*-SC lowered the number of FP branches to nearly zero (0–3.4 branches) on average, at the cost of TP branches. Star-CDP maintained more TPs than PAUP*-SC while also lowering FPs compared to than other methods tested (excluding PAUP*-SC). As a result, Star-CDP achieved the highest mean f1-score on 11/12 model conditions, closely followed by Startle-ILP who achieved the highest mean f1-score on the remaining 1/12 model conditions. In contrast, Cassiopeia-Greedy, Startle-NNI (C++), and LAML had the lowest f1-scores and the highest RF error. LAML’s poor performance is likely due to the poor performance of its starting tree, estimated with Startle-NNI (C++). Overall, the findings for these three methods do not agree with the results presented in the Startle study, in which Startle-NNI substantially lower RF error compared to Cassiopeia-Greedy (Fig. 2a in [29]). A key difference is that our study ran the latest version of Startle-NNI (implemented in C++), whereas the Startle study ran the earliest version of Startle-NNI (implemented in Python). When we ran the original version implemented in Python, Startle-NNI had much lower RF error than Cassiopeia-Greedy and similar RF error to PAUP*. Lastly, it is worth noting that StarCDP-Bias often had dramatically lower FPs compared to StarCDP-Rand but similar TPs (we conjecture this is due to the effectiveness of SH-contraction; however, we could not test this hypotheses because the true binary CLTs for the Startle simulated data sets were not available online).

**Fig. 2.**
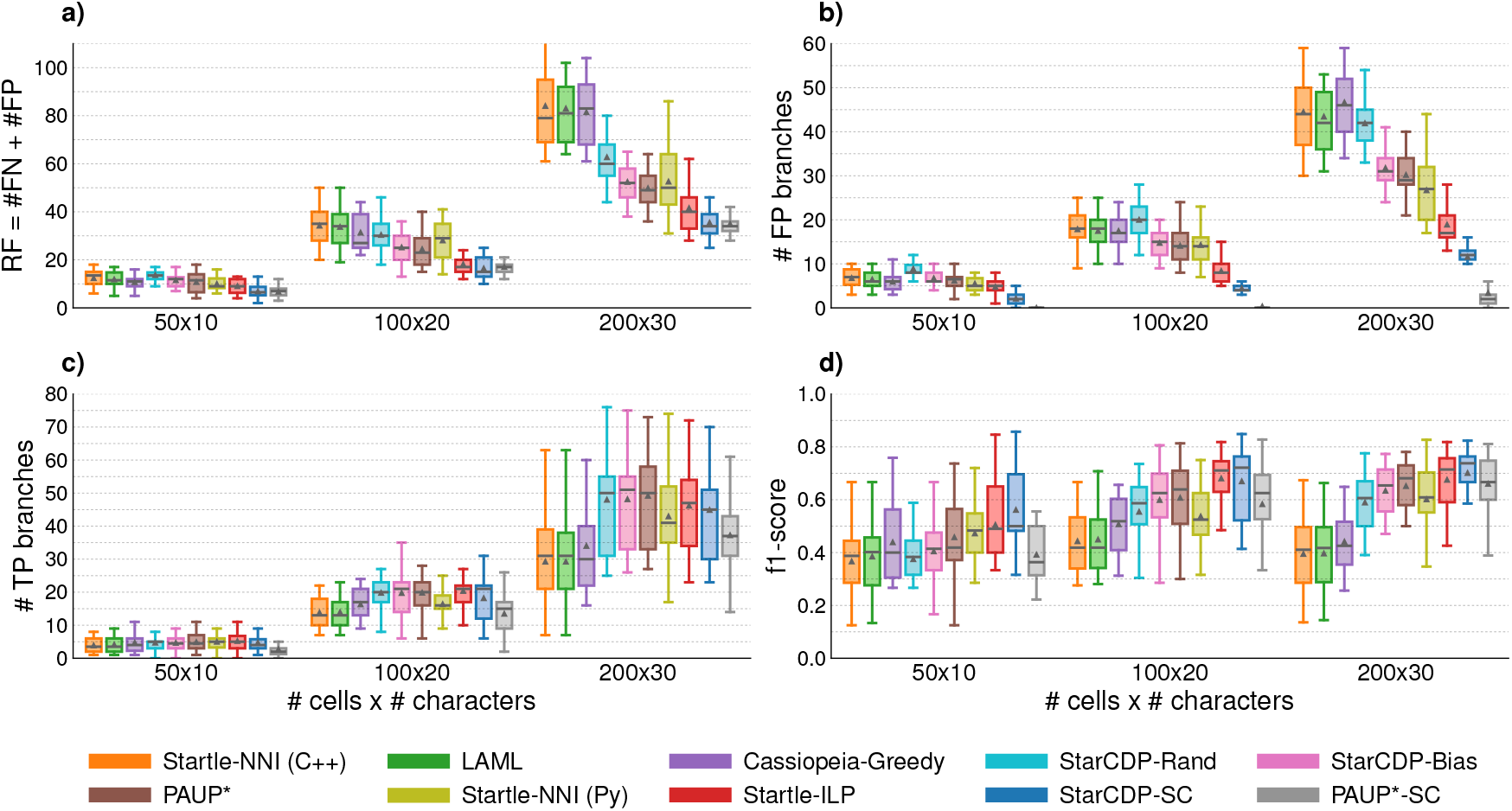
Tree error/accuracy for Startle simulated data sets after contracting mutationless branches in both the true and estimated trees. **(a)** and **(b)** show error metrics (lower is better). **(c)** and **(b)** show accuracy metrics (higher is better). Triangles and bars are means and medians, respectively, across replicates. Note that Startle-ILP was not run on the 200 × 30 matrix in Startle study due to their 8-hour time limit [29].

#### Normalized SHP Score

Across all model conditions, StarCDP-Bias, StarCDP-Ran, PAUP*, and Startle-ILP achieved normalized SHP scores of ∼1 (Figs. S6–S7 in the Supplement). Scores were slightly higher for Startle-NNI (Python) but still close to one. Startle-NNI (C++) and LAML had scores even farther from 1, typically between 1.125–1.875. The worst scores were from StarCDP-SC and PAUP*-SC, as expected because building a strict consensus effectively contracts many branches in a single high scoring tree, increasing its SHP score.

#### Runtime

Most methods were fast taking less than a minute to run on average for all model conditions (Figs. S8–S9 and Table S3 in the Supplement). The exceptions were Startle-NNI (Python; 16 threads) and LAML, which took on average 78.4 minutes and 312.4 minutes, respectively, on the largest data matrix.

### 4.3 Results on LAML Simulated Data Sets

We now present the results of benchmarking methods on the LAML simulated data sets.

#### Tree error/accuracy <u>after</u> SH-contraction

Unlike for the Startle simulated data sets, relative tree error and accuracy across methods depended greatly on the model condition (Fig. 3; Fig. S10 and Table S4–S7 in the Supplement). At 0–50% heritable missing data, PAUP* and StarCDP outperformed Startle-NNI (C++) and LAML, with StarCDP-SC and PAUP*-SC achieving the highest mean f1-score (between 0.68–0.72); in contrast, the mean f1-score of Startle-NNI and LAML was between 0.37–0.64. At 75% heritable missing data, StarCDP-SC, PAUP*-SC, and LAML all tied for mean f1-score (0.76). Finally, at 100% heritable missing data, Startle-NNI (C++) and LAML outperformed Star-CDP and PAUP*, with mean f1-scores of 0.83 and 0.85, respectively; under this condition, the mean f1-score of StarCDP-SC and PAUP*-SC was 0.80 and 0.79, respectively. Although StarCDP-SC and PAUP*-SC performed similarly in terms of f1-score, they differed in terms of TPs and FPs, as observed for the Startle simulated data sets. It is worth noting that the number of FPs for PAUP*-SC was ∼10 times higher on the LAML versus the Startle simulated data sets (30.3-34.2 for 250 cells compared to 3.4 for 200 cells).

**Fig. 3.**
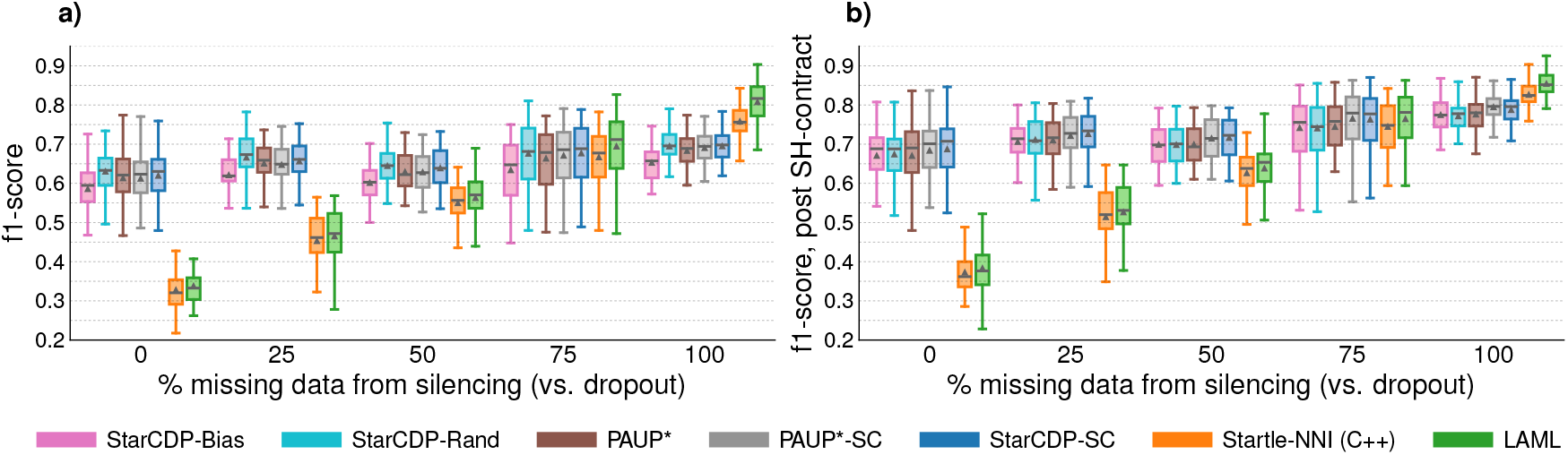
Tree accuracy for LAML simulated data sets with 250 cells. **(a)** shows f1-score (note that the true tree is binary and the estimated trees are binary except PAUP*, PAUP*-SC, and StarCDP-SC). **(b)** shows f1-score computed after taking the SH-contraction of both the true and estimated trees.

#### Tree error/accuracy <u>before</u> SH-contraction

When mutationless branches were not contracted in both true and estimated trees, the f1-scores of all methods decreased. Additionally, LAML’s advantage on model conditions with 75% and 100% heritable data increased; otherwise, trends in tree error and accuracy were similar. It is also interesting that StarCDP-Bias performed much worse than StarCDP-Rand without SH-contraction. This seems reasonable because the StarCDP-Bias tree favors imbalanced tree shapes, but the true CLTs should be fairly balanced as they were simulated under a birth only model.

#### Normalized SHP score and Runtime

Star-CDP and PAUP* methods took just a few minutes and achieved normalized SHP scores close to ∼1 (Fig. S11 in the Supplement). Interestingly, the normalized SHP score of Startle-NNI and LAML dropped from 1.175 to ∼1 as the proportion of heritable missing data increased from 0 to 100 percent; simultaneously, the mean runtime of LAML dropped from 144 to 51 minutes.

## 5 KP-Tracer Analysis

### 5.1 Methods

We also evaluated methods on biological data sets, reanalyzing lineage tracing data from the KP-Tracer mouse model of lung adenocarcinoma [39]. We focused on three metastatic data sets: two KP tumors, `3724_NT_All` (primary tumor and four metastasis locations: 3 lung, 1 soft tissue) and `3513_NT_T1_Fam` (primary tumor and two metastasis: kidney and lymph), as well as one KPL tumor, `3515_Lkb1_T1_Fam` (primary tumor and six metastasis locations: 3 kidney, 2 lung, 1 lymph). The plausibility of CLTs reconstructed on `3724_NT_All` in terms of the number of inferred migration and reseeding events was previously evaluated in the Startle and LAML studies [29, 15]; the Startle study additionally reported SHP score and runtime. We evaluated methods under these same metrics. Additionally, we evaluated the impact of data processing pipelines used in prior studies on relative method performance (see Section 2 of the Supplement for details).

In the Startle study, cells with missing data were clustered into *equivalence classes (ECs)* (i.e., cells that were equivalent excluding missing data) and then **pruned** from the matrix before reconstructing the CLT with Startle-NNI; later, these cells were then placed back into the reconstructed tree as a polytomy, which does not change the SHP score (Fig. 4). The resulting Startle-NNI tree was compared to the published Cassiopeia tree [39], which had not been estimated on the complete (i.e., unpruned) data set. Later, in the LAML study, the resulting Startle-NNI tree was given to LAML as a starting tree, after **duplicate** cells (i.e., cells that are equivalent including missing data) were removed. We refer to this end-to-end process as **preprocessing pipeline 2: pruning, depruning, and deduplicating (PDD)**. Alternatively, the duplicate cells could be removed from the data matrix before CLT reconstruction, referred to as **preprocessing pipeline 1: de-duplicating**. Both preprocessing pipelines 1 and 2 yield a CLT on the same deduplicated cell set; thus, trees are comparable in terms of RF distance as well as inferred migration and reseeding events.

**Fig. 4.**
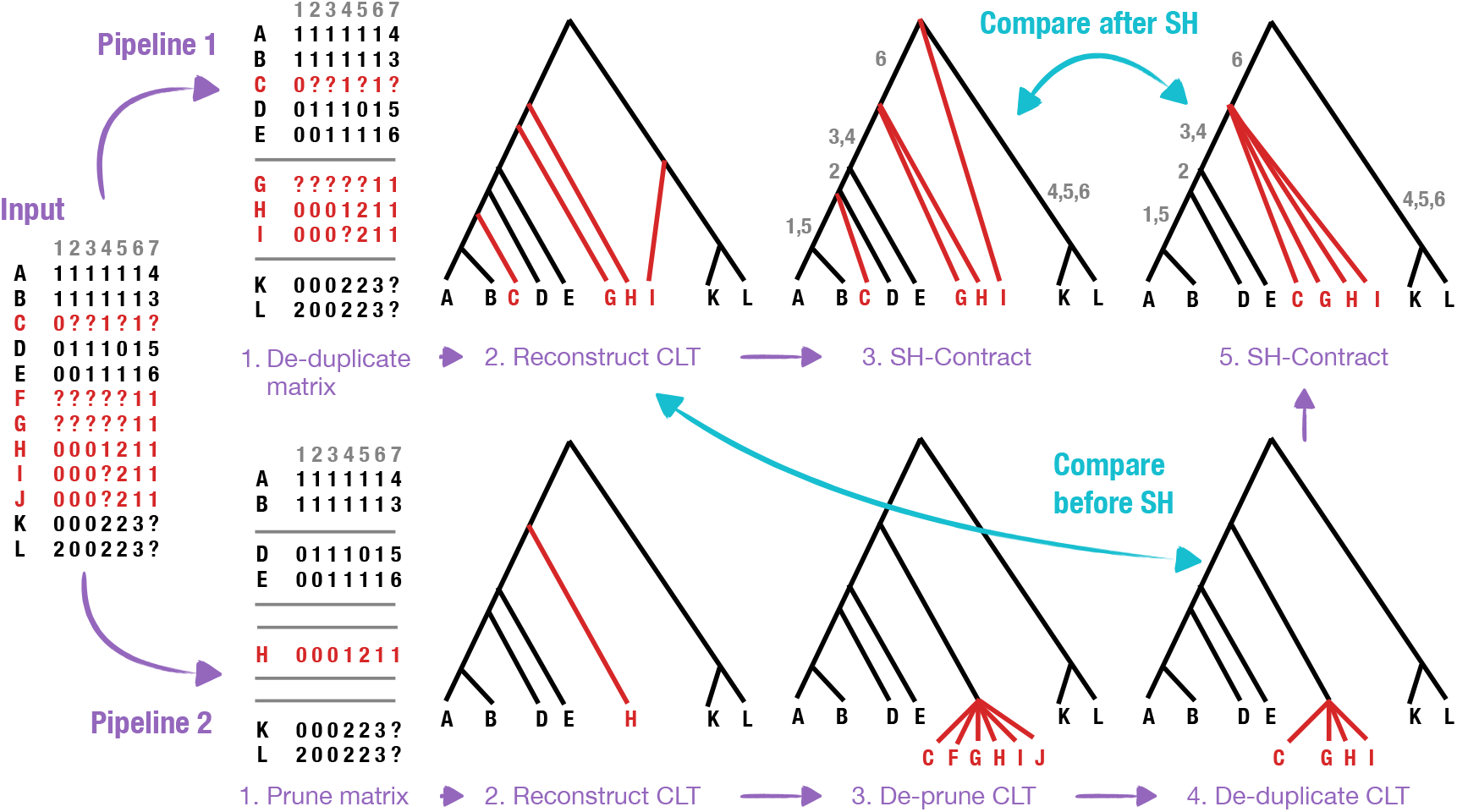
Data processing pipelines and SH-contraction. This figure shows the two data processing pipelines: #1) the de-duplicate only pipeline (resulting CLT is produced in step 2) and #2) the prune, deprune, deduplicate (PDD) pipeline (resulting CLT is produced in step 4). It also shows the SH-contraction of the resulting CLTs (grey numbers on edges indicate mutations; edges without mutations are contracted). CLTs can be compared either before or after SH-contraction.

After a CLT *T* is produced on the deduplicated data set, either with pipeline 1 or 2, LAML can be used to perform post-processing. We consider three different post-processing strategies:

a. **LAML-branch:** estimate branch lengths on *T*, which does not change topology of *T*,
b. **LAML-resolve:** resolve polytomies in *T* after SH-contraction, and
c. **LAML-search:** perform a full tree search with NNI moves, starting from *T*.

Combining the two preprocessing options and the three postprocessing options gives six different pipelines. For each of the six pipelines, we performed CLT reconstruction with six methods: Startle-NNI (C++), StarCDP-Bias, StarCDP-Rand, StarCDP-SC, PAUP*, and PAUP*-SC. This yielded a total of 36 estimated CLTs for each of the three KP-Tracer tumors. We also computed evaluation metrics for the published Cassiopeia trees, after de-duplicating leaves and PDD leaves (see Fig. S2 in the Supplement).

The impact of these preprocessing steps on CLT reconstruction, but they can have a dramatic affect on number of cells (Table 1; also see Table S1 in the supplement for more information about the number and size of ECs). Most notably, the KP-Tracer `3724_NT_All` tumor dropping from 21108 cells down to 1461 and 1207 cells after deduplicating and pruning, respectively. Even after reducing the number of cells, the three KP-Tracer metastatic tumors also have very different information content in terms of the ratio of cells to parsimony-informative sites, with `3724_NT_All` having the lowest ratio and `3515_Lkb1_T1_Fam` having the highest ratio.

**Table 1.**
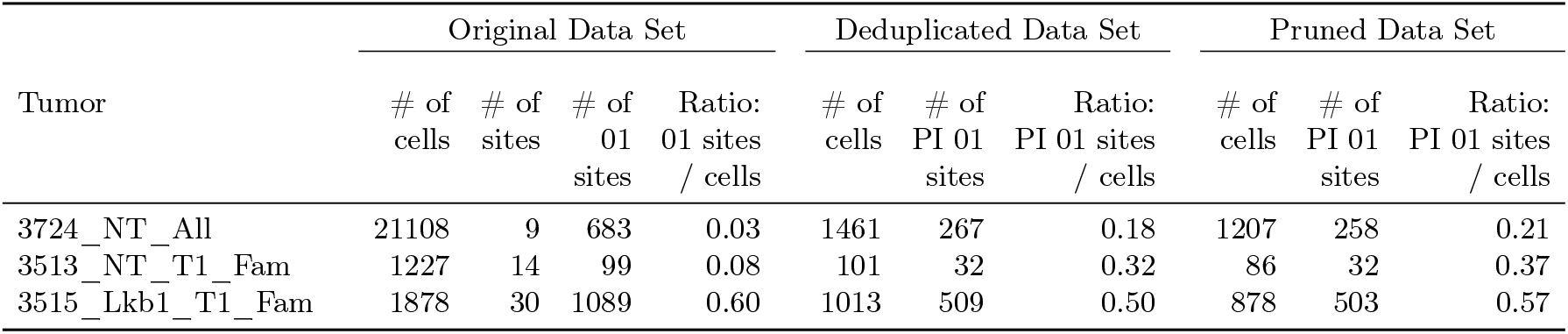
Properties of KP-Tracer data sets. Columns 2, 5, 8 are the number of cells in the original, deduplicated, and pruned data sets, respectively. Column 3 is the number of multi-state sites in the original data set, and column 4 is the number of sites after converting each multi-state site into a collection of 2-state (01) sites. Columns 6 and 9 are the number of *parsimony-informative (PI)* 2-state (01) sites for the deduplicated and pruned data sets, respectively.

|  | Original Data Set |  |  |  | Deduplicated Data Set |  |  | Pruned Data Set |  |  |
| --- | --- | --- | --- | --- | --- | --- | --- | --- | --- | --- |
|  | # of cells | # of sites | # of 01 sites | Ratio: 01 sites / cells | # of cells | # of PI 01 sites | Ratio: PI 01 sites / cells | # of cells | # of PI 01 sites | Ratio: PI 01 sites / cells |
| Tumor |  |  |  |  |  |  |  |  |  |  |
| 3724_NT_All | 21108 | 9 | 683 | 0.03 | 1461 | 267 | 0.18 | 1207 | 258 | 0.21 |
| 3513_NT_T1_Fam | 1227 | 14 | 99 | 0.08 | 101 | 32 | 0.32 | 86 | 32 | 0.37 |
| 3515_Lkb1_T1_Fam | 1878 | 30 | 1089 | 0.60 | 1013 | 509 | 0.50 | 878 | 503 | 0.57 |

**Table 2.**
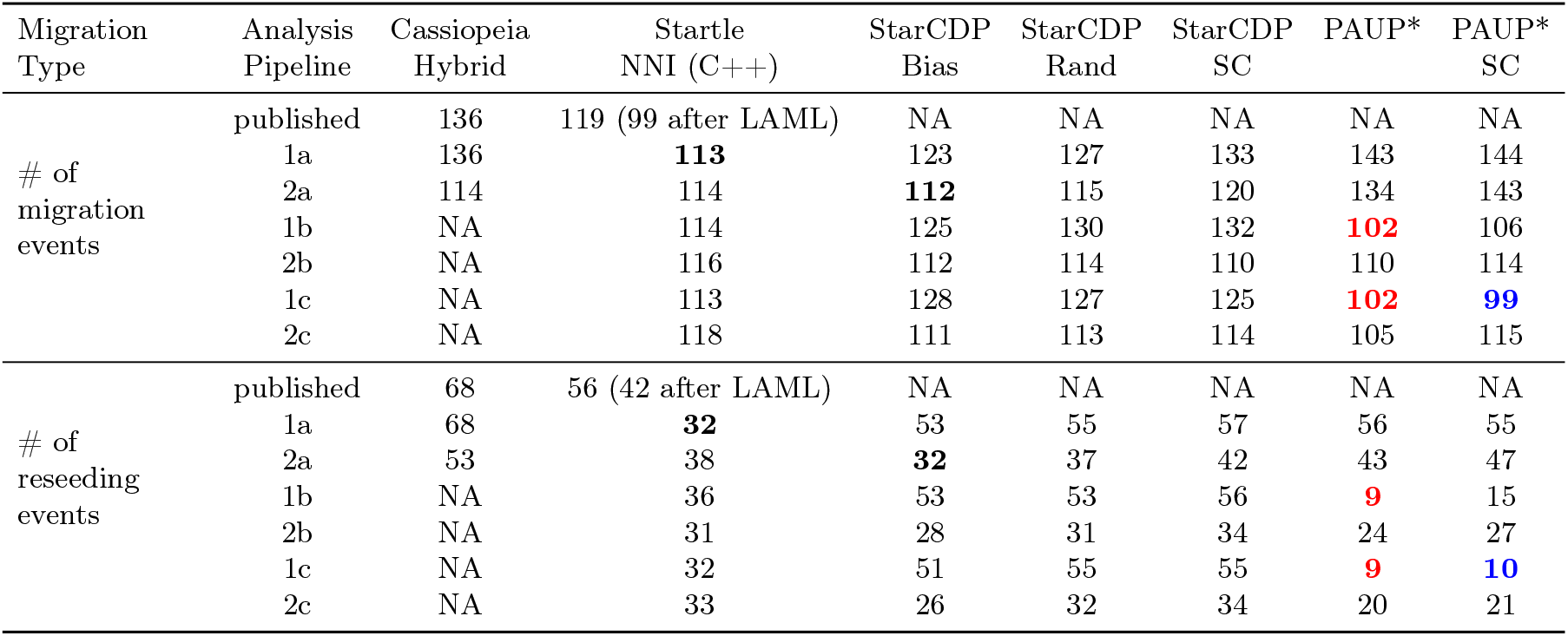
Number of migrations and reseedings inferred for KP-Tracer tumor *3724_NT_All*.

### 5.2 Results on KP-Tracer Tumor 3724_NT_All

We now present results for KP-Tracer tumor `3724_NT_All`, which was previously analyzed in the Startle and LAML studies. Our main focus was on inferring migrations and reseeding events using the reconstructed CLTs CLTs, implementing the methodology proposed in the Startle study [29] and also employed in the LAML study [15]. To test our code, we first replicated the results reported in the LAML study for their published Startle-NNI tree and LAML tree, finding 119 migrations (56 reseedings) and 99 (42 reseedings), respectively (Table S4 in [15]). We also replicated their results for the Cassiopeia-Hybrid tree published by Yan *et al*. [38] after removing duplicate cells, finding 136 migrations and 68 reseedings. Because Yan *et al*. [38] ran Cassiopeia-Hybrid on the complete cell set, we pruned-depruned-deduplicated (PDD) on the Cassiopeia-Hybrid tree to enable a fair comparison to Startle-NNI and LAML (Fig. S2 in the Supplement). The PDD pipeline dropped the number of migrations and reseeding for the published Cassiopeia-Hybrid tree below that of the published Startle-NNI tree (114 migrations and 53 reseedings compared to 119 migrations and 56 reseedings). Likewise, the SHP score of the Cassiopeia tree dropped from 4824.7 to 4759.1, putting it much closer to the published Startle-NNI tree (SHP score: 4715.5). To summarize, the trends in relative method performance changed when the data processing pipeline (PDD), with Cassiopeia-Hybrid outperforming Startle-NNI.

Next we looked at our own CLTs reconstructed with the six pipelines and six methods (Tables S8–S10 and S14 in the Supplement). We foudn that PAUP*-SC run within pipeline 1c (i.e., deduplicate, followed by LAML-search) achieved the same number of migrations as in the published LAML tree (99) but reduced the number of reseedings to 10 (from 42), making it the most plausible tree reconstructed to date. However, to reconstruct this tree, we had to run LAML for 24 hours, making it the most time consuming method tested, and even after 24 hours LAML had not converged. The only other approaches to achieve a similar number of migrations and reseedings was PAUP* within the dedup pipeline followed by either LAML-resolve or LAML-search, which yielded the same CLT with 102 migrations and 9 reseedings. Further inspection revealed that LAML-search may not have executed properly and was effectively the same as LAML-resolve, potentially due to running LAML with 16 threads (instead of 1 thread like in our simulation study). In any case, replacing LAML-search (pipeline 1c with PAUP*-SC) with LAML-resolve (pipeline 1b with PAUP*) dropped the runtime of LAML from 24 hours to just 23 minutes, with only a small change in the number of migrations and reseedings. Additionally the (binary) trees produced by this pipeline were fairly similar: they shared 844 branches (56%) before SH-contraction and 362 branches (87%) after SH-contraction.

When considering the preprocessing pipelines without LAML, StarCDP-Rand was the best method for the PDD pipeline (112 migrations and 32 reseedings), and Startle-NNI (C++) was the best method for the dedup pipeline (113 migrations and 32 reseedings). The StarCDP-Rand tree shared 241/1459 (17%) branches with the best tree (pipeline 1c with PAUP*-SC) branches before SH-contraction and 192/415 branches (46%) after SH-contraction. In contrast, the Startle-NNI tree we estimated shared 70/1459 (5%) with the best tree (pipeline 1c with PAUP*-SC) before SH-contraction and 41/415 (10%) after SH-contraction. The take-away is that the StarCDP-Rand tree and the PAUP*-SC tree had similar numbers of migrations and reseedings but very different topologies. Lastly, the best SHP scores were produced by PAUP*, StarCDP-Rand, and StarCDP-Bias within the PDD pipeline (no LAML).

### 5.3 Results on KP-Tracer Tumor 3513_NT_T1_Fam

The overall trends observed for KP-Tracer tumor 3513_NT_T1_Fam were similar to those observed for the `3724_NT_All`, except the data sets had fewer cells and thus migrations (Tables S7–8 and S11 in the Supplement). The best migration results were obtained on the deduplicate data sets by StarCDP-SC/LAML-search (23 migrations and 0 reseedings), PAUP*/LAML-resolve (20 migrations and 2 reseedings), and PAUP*+LAML-search (19 migrations and 3 reseedings). When considering the preprocessing pipelines without LAML, StarCDP-Rand chieved the fewest migrations for the deduplicate pipeline 2a (25 migrations and 2 reseedings) and Startle-NNI achieved the fewest migrations for the PDD pipeline 1a (28 migrations and 7 reseedings). Both of these trees shared 7/28 (25%) branches with the StarCDP*-SC tree described above, with zero reseedings.

### 5.4 Results on KP-Tracer Tumor 3515_Lkb1_T1_Fam

The trends were somewhat different on KP-Tracer tumor `3515_Lkb1_T1_Fam` (Tables S7–8 and S12 in the Supplement). StarCDP-Bias achieved the lowest number of migrations for both the dedup (1a) and PDD (1b) pipelines without LAML: 145 migrations with 3 reseedings and 154 migrations with 4 reseedings, respectively. In contrast, Startle-NNI (C++) had 170 migrations and 12 reseedings and 161 migrations and 5 reseedings on the dedup (1a) and PDD (1b) pipelines. Thus, the StarCDP trees were more plausible tree than Startle-NNI trees. The only other approach that was close in performance to Startle-Bias ran StarCDP-Rand on the deduplicate data set followed by LAML-resolve (1b); this resulted in 146 migrations and 2 reseedings. Overall, LAML did not have much of an impact on these data, which makes sense given that the level of missing data was much lower and the informativeness was much higher.

## 6 Discussion

CLT reconstruction methods designed for lineage tracing data have garnered much attention in recent years. Despite significant advances, CLT reconstruction continues to be challenged by low phylogenetic signal due to small number of target sites, early saturation of sites, and missing data. To address these issues, we introduced the clade-constrained SHP problem and then presented dynamic programming algorithms, collectively called StarCDP. A key feature of StarCDP is its ability to construct rooted consensus trees from all solutions in the constrained search space. Our study did not explore techniques for building clade constraints for StarCDP but found that effective clade constraints could be generated using techinques from the popular species tree estimation method ASTRAL-III.

In our simulation study, StarCDP’s strict consensus was useful for reducing false positive branches without removing too many true positives, enabling it to achieve greater accuracy (f1-score) than a highest-scoring tree found by PAUP*, PAUP*’s strict consensus, as well as the leading methods: Startle-NNI and Startle-ILP. However, there was little in performance of StarCDP-SC and PAUP* under RF error. Based on these findings, we <u>recommend</u> CLT benchmarking studies incorporate other tree error and accuracy metrics beyond RF error.

Simulations also revealed the utility of PAUP*, a phylogenetics method, which despite its popularity for species phylogenetics, has never been tested as a heuristic for SHP until our study (although see related work by [26]). In simulations, PAUP*’s strict consensus tree was highly unresolved, with a very low number of false positives branches. The utility of consensus trees is in line with the results of the DREAM challenge [9], which reported that taking the consensus of CLTS produced by different *methods* improved accuracy. Based on these results, we <u>recommend</u> that methods based on heuristic search enable users to save trees found during search and build consensus trees from them to enable applications where it is critical to minimize FP branches.

Finally, we evaluated methods on KP-Tracer tumors within two data preprocessing pipelines: one that only deduplicates cells with missing data, and one that prunes cells with missing data (based on equivalence classes) and then deprunes and deduplicates (PDD). This analysis showed that StarCDP (either Rand, Bias, or SC) inferred the lowest number of inferred migrations for all three tumors under the PDD pipeline and one tumor under the deduplicate pipeline (although the best methods changed after applying LAML postprocessing). More importantly, our analysis revealed that data processing pipelines can change relative method performance, swapping the effectiveness of the published Startle-NNI and Cassiopeia-Hybrid trees on the `3724_NT_All` tumor in terms lowering migrations and reseedings from what was reported in the Startle and LAML studies. Thus, we <u>recommend</u> that all methods be evaluated with similar data processing techniques to ensure fair comparisons and to identify whether it is changes in the CLT reconstruction method or the data processing pipeline that is driving relative performance.

Lastly, by exploring different pipelines, our study reconstructed produced the most plausible history of KP-Tracer tumor `3724_NT_All` to date, keeping the number of migrations the same as the LAML study but dropping the number of migrations from 42 to just 10. Given the nature of heuristic search (no guarantees), we emphasize that an advantage of running many permutations of analyses gave us more opportunities to get lucky during the tree search.

## Supporting information

starcdp-supplement

## 8 Acknowledgements

We thank Henri Schmidt, Gillian Chu, and Dr. Palash Sashittal, Dr. Uyen Mai for helpful information about installing software and data sets used in the Startle and LAML studies.

