## Supplementary material for "Dynamic programming algorithms for fast and accurate cell lineage tree reconstruction from CRISPR-based lineage tracing data": starcdp-supplement

November 15, 2024

**References** **45**

### 1 Supplemental Proofs

**Theorem 1.** *Let $T$ be a rooted tree and let $c$ be a character, both on cell set $S$. Assume that there is no missing data so $c$ does not map to the ambiguous state. Then, for any vertex $v \in V(T)$, the Star Homoplasy labeling $\hat{c}[v]$ is unique and only depends on $Clade(v)$; no other information about $T$ is needed.*

*Proof of Theorem 1.* If $v \in L(T)$, then $\hat{c}[v]$ must be set to the only element in $Clade(v)$ by definition. Now consider an arbitrary internal vertex $v \in V(T) \setminus L(T)$. Let $st(v)$ denote the set of state assignments for all leaves that are descendants of $v$, i.e., $st(v) = \{c[l] : l \in Clade(v)\}$. There are two cases.

- **Case A:** $|st(v)| = 1$. Let $\mathtt{s}$ be only state in $st(v)$. We claim that any SH labeling $\hat{c}$ for $(T, c)$ must satisfy $\hat{c}[v] = \mathtt{s}$. Assume for the sake of contradiction, there exists an SH labeling $\hat{c}'$ for $(T, c)$ with $\hat{c}'[v] = \mathtt{s}' \neq \mathtt{s}$. There are three cases to consider. (A.1) $\mathtt{s} \neq \mathtt{0}$ and $\mathtt{s}' \neq \mathtt{0}$, so non-modifiability is violated. (A.2) $\mathtt{s} = \mathtt{0}$ and $\mathtt{s}' \neq \mathtt{0}$, so non-modifiability is violated. (A.3) $\mathtt{s} \neq \mathtt{0}$ and $\mathtt{s}' = \mathtt{0}$, so there must be at least two edges in the subtree below $v$ with mutation $\mathtt{0} \mapsto \mathtt{s}$. Thus, assigning state $\mathtt{s}$ to vertex $v$ would reduce the SHP score and still satisfy non-modifiability. Thus, $\hat{c}'$ is not an SH labeling, which is a contradiction.

- **Case B:** $|st(v)| > 1$. We claim that any SH labeling $\hat{c}$ for $(T, c)$ must satisfy $\hat{c}[v] = \mathtt{0}$. Assume for the sake of contradiction, there exists an SH labeling $\hat{c}'$ for $(T, c)$ with $\hat{c}'[v] = \mathtt{s} \neq \mathtt{0}$. There are two cases to consider. (B.1) There is a leaf below $v$ is in the unedited state $\mathtt{0}$ and there is a leaf below $v$ in some edited state $\mathtt{s}'$, which may or may not equal $\mathtt{s}$. Regardless, non-modifiability is violated. (B.2) There are two leaves below $v$ with different edited labels $\mathtt{s}'$ and $\mathtt{s}''$, which may or may not equal $\mathtt{s}$. Again, non-modifiability is violated. Thus, $\hat{c}'$ is not an SH labeling, which is a contradiction.

To summarize, for any vertex $v \in V(T)$, $\hat{c}[v]$ is unique and only depends on $Clade(v)$. □

**Theorem 2.** *Let $T$ be a rooted tree and let $c$ be a character, both on the same cell set $S$. Assume that $c$ contains missing values (i.e., it maps at least one cell to the ambiguous state denoted by $\mathtt{-1}$). Then, we can define a unique SH labeling $\hat{c}_*$ for $(T, c)$ such that for any vertex $v \in V(T)$, $\hat{c}_*[v]$ only depends on $Clade(v)$; no other information about $T$ is needed.*

*Proof.* Let $R \subseteq S$ be the subset of leaves not assigned the ambiguous state $\mathtt{-1}$ and let $T|_R$ and $c|_R$ be the restriction of $T$ and $c$ to $R$ respectively. By Theorem 1, we can find a unique SH labeling for $(T|_R, c|_R)$ with SHP score $Star(T|_R, c|_R)$. If we add the leaves in $S \setminus R$ back to $T|_R$ the best case is that we do not increase the SHP score. Thus, our goal is to define a unique SH labeling $\hat{c}_*$ for $(T, c)$ with SHP score $Star(T|_R, c|_R)$.

We now propose a procedure for producing $\hat{c}_*$; later we will show $\hat{c}_*$ is an SH labeling. Let $st_*(v)$ be the non-ambiguous states of leaves that are descendants of $v$ (i.e., $st_*(v) = \{c[l] : l \in Clade(v)\} \setminus \{\mathtt{-1}\}$), Then there are three conditions for assigning a state to $v \in V(T)$.

- **Condition 0:** $|st_*(v)| = 0$

  If condition 0 holds, we set $\hat{c}_*[v] = \mathtt{-1}$.

- **Condition 1:** $|st_*(v)| = 1$

  If condition 1 holds, we set $\hat{c}_*[v] = \mathtt{s}$ to the only non-ambiguous state in $st_*(v)$.

- **Condition 2:** $|st_*(v)| > 1$

  If condition 2 holds, we set $\hat{c}_*[v] = \mathtt{0}$.

To show that $Star(T, \hat{c}_*) = Star(T|_R, c|_R)$, we partition verties in $V(T) \setminus L(T)$ into three sets based on the formation of $T|_R$.

- First, we identify every edge incident to the root of a maximally-size subtree of $T$ whose leaves are labeled by the ambiguous state. To be more specific, we identify every edge $a \mapsto b \in E(T)$ such that all leaves in $Clade(b)$ are assigned state $\mathtt{-1}$, and there is at least one leaf in $Clade(a)$ is assigned a non-ambiguous state. Let $\mathcal{E}$ denote the set of such edges.

- Second, we delete each edge in $\mathcal{E}$ but not its endpoints from $T$. This yield $T|_R$, which is $T$ restricted to leaves that are not assigned state `-1`, as well as a collection $\mathcal{P}$ of subtrees of $T$ whose leaves are all assigned state `-1`.

Now consider an arbitrary vertex $u \in V(T)$.

- If condition 0 holds for $u \in \cup_{t\in\mathcal{P}}\{V(t) \setminus L(t)\}$, then $u \in \cup_{t\in\mathcal{P}}\{V(t) \setminus L(t)\}$. Thus, setting state `-1` to $u$ will not increase the SHP score for $T$.
- If condition 1 holds for $u$, then $u \in V(T|_R) \setminus L(T|_R)$ and by applying Theorem 1, we must assign $\hat{c|_R}[u] = \texttt{s}$. In this case, $\hat{c}_*[u] = \hat{c}|_R[u]$, which does not increase the SHP score for $T$.
- If condition 2 holds for $u$, then $u \in V(T|_R) \setminus L(T|_R)$ and by applying Theorem 1 we must assign $\hat{c}|_R[u] = \texttt{0}$. In this case, $\hat{c}_*[u] = \hat{c}|_R[u]$, which does not increase the SHP score for $T$.

Since we hae shown that our labeling does not increase the SHP score for any vertex $u \in V(T)$, $Star(T, \hat{c}_*) = Star(T|_R, c|_R)$ and our defined $\hat{c}_*$ is an SH labeling for $(T, c)$. ☐

Theorem 2 naturally yields Algorithm 1. It is worth noting that in Startle [9, 10], the ILP algorithm may induce a state assignment to some internal vertex $u$ such that condition 0 holds for $u$ with the state as the parent of $u$ assigned, which may lead to a different state assignment for $u$. But both will not increase the SHP score. Thus, they are both valid ways of assigning states to achieve the SHP score.

**Theorem 3.** *Star-CDP is correct.*

*Proof.* **Base Case:** The base case for $Star[A]$ is trivial because when $|A| = 1$, the solution is a leaf given the same states as the input characters in $\mathcal{C}$. There are no mutations, so $Star[A] = 0$.

**Recurrence:** Now let's consider the case where $|A| > 1$. Suppose we have already correctly computed $Star[X]$ and $Star[Y]$ for any $X|Y \in \mathcal{STB}(A)$ by induction hypothesis. Let the pair $(t_X, \hat{\mathcal{C}}_X)$ be an arbitrary solution (i.e., rooted binary tree and SH-labeling) to subproblem $Star[X]$, and similarly let $(t_Y, \hat{\mathcal{C}}_Y)$ be an arbitrary solution to subproblem $Star[Y]$. Joining $t_X$ and $t_Y$ at their roots forms a new rooted binary tree $t_A$ with leaf set $A = X \cup Y$ and $Clade(t_A) \subseteq \Sigma$. By Theorem 2, the state assignments $\hat{\mathcal{C}}_X$ and $\hat{\mathcal{C}}_Y$ for all vertices in $t_X$ and $t_Y$, respectively, will not change after these two trees are joined; thus, the substitutions counted on the internal edges of $t_X$ and $t_Y$ (i.e., their SHP scores) do not change. Lastly, the SH labeling for the root of $t$ can be determined by $\texttt{GetStates}(A, \mathcal{C})$, after which it is easy to compute the cost of mutations on the outgoing edges. To summarize, $t_A$ is a candidate solution for $Star[A]$ with SHP score equal to $Star[X] + Star[Y] + \texttt{SHPCost}(A, X, Y, \mathcal{C}, \mathcal{W})$. However, a better SHP score could be obtained from joining another pair of trees on different leaf sets than $X$ and $Y$ but whose combined leaf sets still equals $A$. All candidate solutions can be formed by looking at all possible subtree bipartitions on $A$, i.e., $X|Y \in \mathcal{STB}(A)$. Thus, the $Star[A] = min_{X|Y \in \mathcal{STB}(A)} Star[X] + Star[Y] + Cost(X|Y, S, \mathcal{C}, \mathcal{W})$. This is our recurrence. We also store pointers to the two subproblems yielding the solution for $Star[A]$ so that these subproblems can be accessed during backtracking.

**Recurrence is solvable by DP:** In our recurrence, $Star[A]$ only depends on $Star[X]$ and $Star[Y]$ for all $X|Y \in \mathcal{STB}(A)$. Since $|X|$ and $|Y|$ must be less than $|A|$, we can solve subproblems in the order of clade cardinality. This guarantees the trivial subproblems (base cases) are solved first and ensures all subproblems dependencies are satisfied moving forward. Thus, our recurrence is solvable by DP.

**Putting it all together:** After computing $Star[S]$, we can backtrack through the subproblems to find a tree. The resulting tree $T$ is a solution to the CC-SHP problem because it minimizes SHP score and draws its clades from $\Sigma$. ☐

**Theorem 4.** *Star-CDP has time complexity $O(nm|\Sigma|^{1.726} + n|\Sigma|^2)$ and space complexity $O(|\Sigma|)$ excluding the inputs.*

*Proof.* The time cost of solving subproblem $Star[A]$ for clade $A$ is $O(nm|\mathcal{STB}(A)|)$ because we need to run `SHPCost` (Algorithm 2, with time complexity $O(nm)$, for each of the subtree bipartitions in $\mathcal{STB}(A)$. We then need to evaluate all subproblems $A \in \Sigma$, yielding time complexity

$$O\Bigg(\sum_{A\in\Sigma} nm|\mathcal{STB}(A)|\Bigg) = O(nm|\mathcal{STB}|) = O(nm|\Sigma|^{1.726}) \tag{1}$$

where we apply the bound on the number of allowed subtree bipartitions given $\Sigma$ from Kane and Tao [5]. Lastly, we must consider the work done during the preprocessing to build $\mathcal{STB}(A)$ for all $A \in \Sigma$ (Algorithm 3). The trivial upper bound on the preprocessing is $O(n|\Sigma|^2)$, bringing the time complexity of the entire Star-CDP algorithm to $O(nm|\Sigma|^{1.726} + n|\Sigma|^2)$.

In terms of storage, the Star-CDP algorithm saves the solution to each subproblem $Star[A]$ as well as two pointers for backtracking. There are only $|\Sigma|$ subproblems, so the storage is linear in the number of clade constraints. The space complexity increases to $|\Sigma|^{1.726}$ if the allowed subtree bipartitions $\mathcal{STB}$ are computed as part of preprocessing; however, this computation can be integrated into the preprocessing phase to avoid explicitly storing $\mathcal{STB}$. □

**Theorem 5.** *The Mutation- and Clade-Constrained Large Star Homoplasy Parsimony (MCC-LSHP) problem is in P if and only if $P = NP$.*

*Proof.* We prove the hardness of MCC-LSHP by constructing a Karp reduction to MCC-LSHP from Maximum Independent Set for an arbitrary 3-regular graph, which is NP-complete (see proof by Fleischner, Sabidussi, and Sarvanov [4]). To begin, we define optimization problems as decision problems.

**Maximum Independent Set (IS) for 3-Regular Graphs (MIS3).**

- **Input:** A 3-Regular graph $G \in \mathcal{G}_3$ and a positive Integer $K$
- **Output:** YES, if there is an independent set with a size of at least $K$; NO otherwise.

Recall that $G$ is a *3-regular graph* if every vertex in $G$ has exactly 3 neighbors and that an *independent set* is a set of vertices in graph $G$ such that no two of are adjacent. As in [4], we assume $G$ is *finite* and *simple* (i.e., no parallel edges or self loops).

**Mutation- and Clade-Constrained Large SH Parsimony (MCC-LSHP).**

- **Input:** $(S, \mathcal{C}, \Sigma, K')$ where $S$ denotes cell set, $\mathcal{C}$ denotes character matrix, $\Sigma$ is a set of clades that denotes our constrained search space, and $K'$ is a non-negative integer $K'$.
- **Output:** YES if there is a rooted tree $T$ on cell set $S$ satisfying the following conditions:
  (1) $Clade(T) \subseteq \Sigma$
  (2) Given the SH-labeling for $(T, \mathcal{C})$, there is at least one mutation on each non-terminal edge in $T$.
  (3) $\sum_{c\in\mathcal{C}} Star(T, c) \leq K'$

  NO, otherwise.

For simplicity, we assume all mutations have equal weight 1. It is not hard to see that if the unweighted MCC-LSHP problem is NP-Hard, the weighted version of the problem is also NP-Hard. We also assume that all characters in $\mathcal{C}$ have 2-states: `0` = unedited/ancestral and `1` = edited/derived. This is reasonable given that any instance of LSHP can be transformed into an instance of the large Camin-Sokal parsimony (LCKP) problem with 2-state characters. This does not impact our proof because the "SH-labeling" derived in Theorem 1 still applies in the case of LCKP.

**Reduction $\mathcal{F}(G, K) = (S, \mathcal{C}, \Sigma, K')$.** Consider the following reduction $\mathcal{F}$ that takes an instance of `MIS3` $G$ with vertex set $V(G) = [n]$ and positive integer $K$ and returns an instance of `MCC-LSHP` $(S, \mathcal{C}, \Sigma, K')$ with positive integer $K' = |V| - K + |E|$ by the following procedure:

- **Step 1: Build $S$.** We build a set $S$ of cells from each edge in $G$, i.e., $S = \{e : \forall e \in E(G)\}$.
- **Step 2: Build $\mathcal{C}$.** We build a character matrix $\mathcal{C}$ from $G$, where each row in $\mathcal{C}$ represents an edge in $E(G)$ (also a cell in $S$) each column represents a vertex in $V(G)$. We set any entry of $\mathcal{C}$, denoted $\mathcal{C}_{(e,v)}$, to 1 if edge $e$ is incident to vertex $v$ in $G$; otherwise we set entry $\mathcal{C}_{(e,v)}$ to 0.
- **Step 3: Build $\Sigma$.** Lastly, we form $\Sigma$ from $G$ as follows. We add the full edge set $E(G)$ to $\Sigma$. Then, for every vertex $u \in V(G)$, $u$ is incident to three edges $e_1, e_2, e_3$, so we add the following seven clades to $\Sigma$: $\{e_1\}$, $\{e_2\}$, $\{e_3\}$, $\{e_1, e_2\}$, $\{e_1, e_3\}$, $\{e_2, e_3\}$, and $\{e_1, e_2, e_3\}$. We say that these clades are *contributed by* $u$ to $\Sigma$ or that $u$ *is the contributor* of these clades to $\Sigma$. It is easy to see that each of the three singleton clades have two contributor vertices and that the non-singleton clades have exactly one contributor vertex because $G$ is simple.

Note that the first two steps of the reduction $\mathcal{F}$ come from the hardness proof for the *unconstrained* large Camin-Sokal parsimony problem in Day, Johnson and Sankoff [2]; the last step in the reduction is added by us. It is easy to see that the time complexity of $\mathcal{F}$ is time complexity of $O(|E(G)| \cdot |V(G)|)$ because of the construction of the character matrix $\mathcal{C}$ from $G$. It is also easy to see that size of the output $\mathcal{F}(G)$ is bounded by $O(|E(G)| \cdot |V(G)|)$ for any input $G$ of MIS3.

We now claim that $(G, K)$ is a YES instance of `MIS3` if and only if $\mathcal{F}(G) = (S, \mathcal{C}, \Sigma, K')$ is a YES instance to `MCC-LSHP` for $K' = 2(|E(G)| - K)$. Before we give the proof, we make the following observations:

- **Observation 0:** By construction of the character matrix $\mathcal{C}$, the SH-labelings for each cell (i.e., edge in $G$) is a vector in $\{0, 1\}^{|V(G)|}$ with exactly 2 ones and $|V(G)| - 2$ zeros.
- **Observation 1:** Let $\sigma \in \Sigma$ be a clade contributed by an arbitrary vertex $v$ in $G$. By combining *Observation 0* with *Theorem 1*, we can determine the SH-labelings at clade $\sigma$, denoted $\hat{c}[\sigma]$, as follows. First, $v$ is incident to three edges: $e_1 = (v, x)$, $e_2 = (v, y)$, and $e_3 = (v, z)$, where $x \neq y \neq z$ because $G$ is 3-regular and simple. Second, by construction of $\Sigma$, $\sigma$ must take on one of the three forms:
  - *Form 1:* $\sigma = \{e_i\}$ for $i \in \{1, 2, 3\}$.
  - *Form 2:* $\sigma = \{e_i, e_j\}$ for $i \neq j \in \{1, 2, 3\}$.
  - *Form 3:* $\sigma = \{e_1, e_2, e_3\}$

  If $\sigma$ has form 1, then $\hat{c}[\sigma]$ is just the entry of the character matrix $\mathcal{C}$ corresponding to the only edge in $\sigma$, which is a cell (row) in $\mathcal{C}$. If $\sigma$ has forms 2 or 3, then $\hat{c}[\sigma]$ has state 1 for the column corresponding to vertex $v$ in $\mathcal{C}$ and all other entries are zero.
- **Observation 2:** Any rooted tree $T$ on $S$ that is a witness to `MCC-LSHP` cannot contain two different non-trivial clades contributed by the same vertex $v \in V(G)$. By rules for clade compatibility [11], we know that any rooted tree $T$ on $S$ can contain at most three clades induced by vertex $v$ and that these three clades, denoted $\sigma_1$, $\sigma_3$, and $\sigma_3$, must have cardinality 1, 2, and 3, respectively. By *Observation 1*, $\sigma_2$ and $\sigma_3$ must have the same SH-labeling, which means there are no mutations on the edge between the vertices that induce these clades in $T$, violating condition 2.
- **Observation 3:** Any rooted tree $T$ on $S$ that is a witness to `MCC-LSHP` has a root $r$ assigned SH-labeling $\hat{c}[r] = \mathbf{0}$ where $\mathbf{0}$ denotes zero vector. The root $r$ is associated with the full edge set $E(G)$, which we added as a clade to $\Sigma$. By the way of constructing the character $\mathcal{C}$, each column in $\mathcal{C}$ (i.e., a vertex in $V(G)$) has exactly three cells in $\mathcal{C}$ (i.e., edges in $E(G)$) in state 1 and the remaining cells are all 0. By *Theorem 1*, the SH-labeling, returned by *GetState*$(E, \mathcal{C})$, is $\mathbf{0}$.
- **Observation 4:** If we add any other clades to $\Sigma$, its will also be $\mathbf{0}$. This means there are no mutations on the branch between the vertex inducing this clade and the root. Thus, it will be contracted. We therefore do not need to consider adding such clades to $\Sigma$.

From these observations, it is easy to see that any solution to MCC-LSHP given $\mathcal{F}(G)$ must be a non-binary tree. Now we continue with the proof that $(G, K)$ is a YES instance to `MIS3` if and only if $\mathcal{F}(G) = (S, \mathcal{C}, \Sigma, K')$ is a YES-instance to `MCC-LSHP` setting $K' = 2(|E(G) - K|)$.

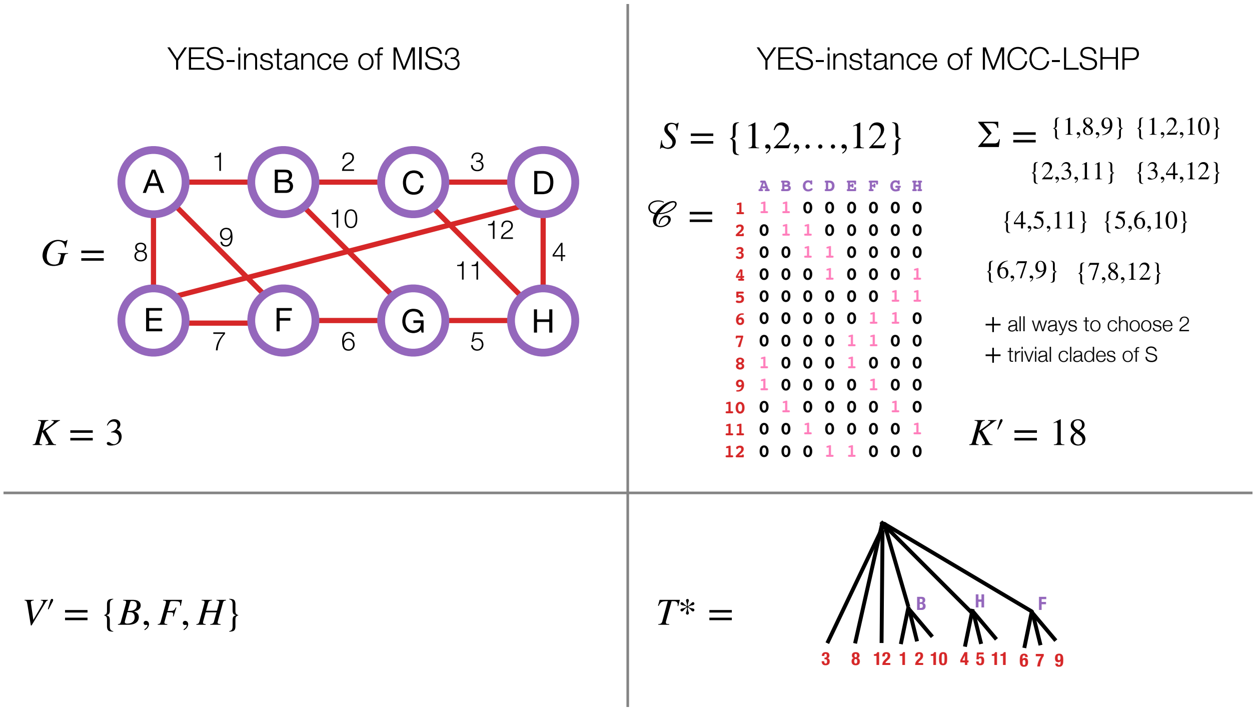

Figure S1: **Karp Reduction Example**

($\Longrightarrow$) Suppose $(G, K)$ is an YES-instance of `MIS3`. Then, there exists an independent set $V'$ for $G$ with the size of at least $K$. We will show that $\mathcal{F}(G) = (S, \mathcal{C}, \Sigma, K')$ is a YES-instance for `MCC-LSHP` for $K' = 2(|E(G)| - K)$. Consider the following procedure:

- Initialize $\Sigma^*$ to be the empty set.
- For each vertex $v \in V'$:
  - Add clade $\{(v, x), (v, y), (v, z)\}$ to $\Sigma^*$ where $x \neq y \neq z \in V(G)$ are the three neighbors of $v$.
- Add every edge in $E(G)$ to $\Sigma^*$.
- Add the full edge set $E(G)$ to $\Sigma^*$.

By construction of $\Sigma^*$, we know that any non-trivial pair of clades satisfies $\sigma_A \cap \sigma_B = \emptyset$ because otherwise $V'$ is not an independent set for $G$. It follows that $\Sigma^*$ is compatible (see Lemma 2.10 and Corollary 2.11 in [11]). Let $T^*$ be the compatibility supertree on $S$ such that $Clade(T) = \Sigma^*$. To conclude our proof, we need to show that $T^*$ satisfies conditions 1–3 for $K' = 2(|E(G)| - K)$. By construction of $\Sigma^*$, $Clade(T^*) = \Sigma^* \subset \Sigma$, so condition 1 holds. By *Observations 1–3*, we can determine the SH-labelings for $(T^*, \mathcal{C})$, which enables us to compute the number of substitutions on any branch $b = u \mapsto v$ in $T^*$. There are four cases to consider:

- **Case 1:** If $u$ is the root and $v$ is an internal (non-root) node, there is 1 substitution on $b$. There are $|V'|$ branches in this category by construction of $\Sigma^*$.
- **Case 2:** If $u$ is an internal (non-root) node and $v$ is a leaf, there is 1 substitution on $b$. There are $3|V'|$ branches in this category by construction of $\Sigma^*$.
- **Case 3:** If both $u$ and $v$ are internal (non-leaf) vertices, there are 0 substitutions on $b$. There are 0 branches in this category by construction of $\Sigma^*$; thus, condition 2 holds.

– **Case 4:** If $u$ is the root and $v$ is a leaf, there are 2 substitutions on $b$. There are $|E(G)| - 3|V'|$ branches in this category by construction of $\Sigma^*$.

Putting this all together, the SHP score is

$$Star(T^*, \mathcal{C}) = |V'| + 3|V'| + 0 + 2(|E(G)| - 3|V'|) = 4|V'| + 2|E(G)| - 6|V'| = 2|E(G)| - 2|V'| \quad (2)$$

Because $|V'| \geq K$, $Star(T^*, \mathcal{C}) \leq 2|E(G)| - 2K$, so condition 3 holds. And $(S, \mathcal{C}, \Sigma, K')$ is a YES-instance of `MCC-LSHP`.

($\Longleftarrow$) Conversely, suppose $\mathcal{F}(G) = (S, \mathcal{C}, \Sigma, K')$ has solution $T^*$ with SHP score at most $K' = 2(|E(G)| - K)$. Consider the tree $\mathcal{T}$ with all of its leaves to a root vertex. By *Observations 1–3*, the SHP score of $\mathcal{T}$ is $2E(G)$. We can reduce the SHP score by adding internal vertices corresponding to non-trivial clades. By construction of $\Sigma$, we added clades of size 2 or 3 but these clades cannot be nested (*Observation 2*). If we add $x$ clades of size 2 to $\mathcal{T}$, its SHP score is reduced by $x$. If we add $y$ clades of size 3 to $\mathcal{T}$, its SHP score is reduced by $2y$. Thus, we could must add at least $y = K$ clades of size 3 or $2K$ clades of size 2 (or some number of clades in between with various sizes) for $\mathcal{T}$ to have SHP score at most $2E(G) - 2K$. Thus, there are at least $K$ non-trivial clades in $Clade(T^*) = \Sigma^* \subset \Sigma$. For each such clade, denoted $\sigma'$, we can add the vertex $v \in V(G)$ that is incident to all cells in $\sigma'$ (which correspond to edges in $E(G)$) to $V'$, which is initially the empty set. By construction, $V'$ has size at least $K$. It remains to be shown that $V'$ is an independent set for $G$. Consider any pair of $\sigma_A, \sigma_B$ of non-trivial clades in $\Sigma^*$. We know that $\sigma_A \cap \sigma_B = \emptyset$ because otherwise clade compatibility is violated (Lemma 2.10 in [11]). The elements of $\sigma_A$ and $\sigma_B$ are cells in $S$ (which correspond to edges in $E(G)$), so the vertex $v_A$ and $v_B$ added to $V'$ for clades $\sigma_A$ and $\sigma_B$, respectively, cannot be incident to the same edge in $G$ (i.e., $v_A$ and $v_B$ are not neighbors). Because this holds for all pairs of non-trivial clades, $V'$ is an independent set.

Since both directions hold, we show that MCC-LSHP is in $P$ if and only if $P = NP$. □

See main text for some notes about similarity to the hardness proof for (unconstrained) Camin-Sokal parsimony by Day, Johnson, and Sankoff [2].

**Theorem 6.** *The DP algorithms for computing Below and Above are correct.*

*Proof.* We have already argued the correctness of recurrences in the main text, so we only need to argue there exists an iteration that correctly fills in the DP matrices. $Below[A]$ depends on $Below[X], Below[Y]$ for all $X, Y \in I[A]$. $|X|$ and $|Y|$ must less than $|A|$. Thus, computing all subproblems for *Below* in *increasing* order of cardinality guarantees all dependency subproblems are computed starting with the base case. Similarly, $Above[X]$ depends on $Above[A]$ and $Below[Y]$ for all $(A, Y) \in J[X]$. Since we have already correctly computed all subproblems for *Below*, we focus on $Above[X]$. $|A|$ must be greater than $|X|$. Thus, computing all subproblems for *Above* in *decreasing* order of cardinality guarantees all dependency subproblems are computed starting with the base case. □

**Theorem 7.** *Counting the number of solutions to CC-LSHP does not increase time or space complexity. Computing clade frequencies $Freq[A]$ for all $A \in \Sigma$ does not increase time complexity but it increases space complexity to $O(|\Sigma|^{1.726})$.*

*Proof of Theorem 7.* To compute clade frequencies, we first need to compute *Above* and *Below*. The DP matrices *Above* and *Below* have the same size as *Star* so they do not increase the space complexity. The calculation of *Below* can be integrated into the DP algorithm for *Star* (Algorithm 4). No additional time or space is required because the subtree bipartitions stored in $I$ do not need to be maintained across subproblems. A consequence is that we can count the number of solutions to CC-LSHP without increasing the time or space complexity (as this value is stored in $Below[S]$ where $S$ is the full cell set). The calculation of *Above* requires $J$ to be saved during the DP algorithm for *Star* (Star-CDP; Algorithm 4), upping the storage requirement from $O(|\Sigma|)$ to $O(|\Sigma|^{1.726})$. To compute *Above*, we iterate over all (reversed) allowed subtree bipartitions stored in $J$. The work per subproblem is constant, so the time complexity is $O(|\Sigma|^{1.726})$. After computing *Below* and *Above*, $Freq[A]$ can be computed for any clade $A \in \Sigma$ in constant time as $Below[A] \cdot Above[A]/Below[S]$. It follows that the calculation of *Freq* does not increase time complexity of Star-CDP, but it does increase storage complexity to $O(|\Sigma|^{1.726})$. □

#### 1.1 Algorithms

**Algorithm 1** `GetStates` has time complexity $O(nm)$ because we can safely assume that $n \gg r$ for CRIPSR/Cas9 lineage tracing

**Input:** Clade $A \subseteq S$ where $S = \{1, 2, \ldots, n\}$ and characters $\mathcal{C} = \{c_j\}_{j=1}^m$ where

- $c_j[i]$ indicates the state of cell $i$ for character $j$
- $r$ is the maximum integer state in $\mathcal{C}$
- each character $c_j$ maps cells to a state in $\mathcal{A} = \{-1, 0, 1, 2, \ldots, r\}$

**Output:** State assignments for vertex in any tree that induces clade $A$

```
function GetStates(A, C)
    states ← an array of length m
    for j ∈ {1, 2, ..., m} do
        found ← an array of zeros of length r + 2
        for x ∈ A do
            found[c_j[x] + 1] ← found[c_j[x] + 1] + 1
        end for
        nstates ← 0; keep ← −1
        for i ∈ {1, 2, ..., r + 1} do
            if found[i] > 0 then
                nstates ← nstates + 1; keep ← i − 1
            end if
        end for
        if nstates ≤ 0 then states[j] ← keep
        else states[j] ← 0
        end if
    end for
    return states
end function
```

**Algorithm 2** `SHPCost` has time complexity $O(nm)$

**Input:** Clades $A, X, Y \subseteq S$ such that $X \cup Y = A$ where $S = \{1, 2, \ldots, n\}$, characters $\mathcal{C} = \{c_j\}_{j=1}^m$, and costs $\mathcal{W} = \{w_j\}_{j=1}^m$ **Output:** SHP cost for outgoing edges of a vertex that induces clade $A$ and has two children that induce clades $X$ and $Y$ respectively

```
function SHPCost(A, X, Y, C, W)
    stA ← GetStates(A, C); stX ← GetStates(X, C); stY ← GetStates(Y, C)
    cost ← 0
    for j ∈ {1, 2, ..., m} do
        if stX[j] ≠ -1 then
            if stA[j] ≠ stX[j] then cost ← cost + w_j[stX[j]]
            end if
        end if
        if stY[j] ≠ -1 then
            if stA[j] ≠ stY[j] then cost ← cost + w_j[stY[j]]
            end if
        end if
    end for
    return cost
end function
```

**Algorithm 3** Preprocessing phase has time complexity $O(n|\Sigma|^{1.726})$ because clades can be stored as bit vectors of size $n$, in which case performing set operations are $O(n)$, checking set membership in $\Sigma$ is an $O(n)$ hash, and using *clade2index* is an $O(n)$ hash

**function** PREPROCESS($\Sigma$)<br>
    Create list *clades* from $\Sigma$ with length $q = |\Sigma|$ as well as list *cladeSizes*<br>
    Do argument sort of *cladeSizes* from least to greatest<br>
    Apply argsort to *cladeSizes*<br>
    Use argsort to create map *clade2index* and list *clades* such that $clade2index[X] = x$ and $clades[x] = X$<br>
    **for** $a \in \{1, 2, \ldots, q\}$ **do**<br>
        $cladeSTBs[a] \leftarrow [\,]$<br>
        **for** $x \in \{1, \ldots, a-1\}$ **do**<br>
            **if** $cladeSizes[x] < cladeSizes[a]$ **then**<br>
                $A \leftarrow clades[a]$; $X \leftarrow clades[x]$<br>
                **if** $X \subset A$ **then**<br>
                    $Y \leftarrow A \setminus X$<br>
                    **if** $Y \in \Sigma$ **then**<br>
                        $y \leftarrow clade2index[Y]$<br>
                        $cladeSTBs[a].append([x, y])$<br>
                    **end if**<br>
            **else**<br>
                break<br>
        **end for**<br>
    **return** $q, clades, cladeSizes, cladeSTBs$<br>
**end function**

**Algorithm 4** Dynamic Programming for CC-LSHP

```
function STAR-CDP(C, W, Σ)
    q, clades, cladeSizes, cladeSTBs ← Preprocess(Σ)
    for a ∈ {1, 2, ..., q} do
        if cladeSizes[a] = 1 then
            Star[a] ← 0
            TraceBack[a] ← NULL
            Below[a] ← 1
        else
            bestStar ← ∞
            for [x, y] ∈ cladeSTBs[a] do
                tmpStar ← Star[x] + Star[y] + SHCost(clades[a], clades[x], clades[y], C, W)
                if bestStar > tmpStar then
                    I ← []; J ← []; bestStar ← tmpStar
                end if
                if bestStar = tmpStar then
                    I[a].append([x, y]); J[x].append([a, y]); J[y].append([a, x])
            end for
            Star[a] ← bestStar
            TraceBack[a] ← first element of I[a]
            Below[a] ← 0
            for [x, y] ∈ I[a] do Below[a] ← Below[a] + Below[x] · Below[y]
    for x ∈ {q, q − 1, ..., 1} do
        if cladeSizes[x] = n then
            Above[x] ← 1
            Freq[a] ← 1
        else
            Above[x] ← 0
            for [a, y] ∈ J[x] do Above[x] ← Above[x] + Above[a] · Below[y]
            Freq[x] ← (Below[x] · Above[x])/Below[q]
    return Star[q], Below[q], clades, Freq, TraceBack
end function
```

#### 2 Supplemental Methods

Scripts used to analyze data are available on Github: `https://github.com/molloy-lab/star-study`.

##### 2.1 Simulated Data Sets

**Startle simulated data sets.** We downloaded the data sets simulated for the Startle study [9, 10] from Github (`https://github.com/raphael-group/startle/blob/main/startle_simulations.zip`). True trees, character matrices, mutation priors were available for each model condition. We focused on the model condition with dropout rate of 0.15 and mutation rate of 0.1 because it had the largest number of replicates (21 in total). Startle-NNI (Python) failed on three replicates with 50 cells and 10 characters (#8, #9, #10), and Startle-ILP failed on one replicate with 50 cells and 20 characters (#1). These four data sets were excluded from analyses to enable a fair comparison across methods. When computing tree accuracy and error metrics, we contracted mutationless branches in both the true and estimated trees because true trees were already non-binary with the vast majority of mutationless branches contracted.

**LAML simulated data sets.** We downloaded the data sets simulated for the LAML study with 250 cells and 30 characters [6, 7] from Github (`https://github.com/raphael-group/laml-experiments/tree/main/sim_tlscl`).

**KP-Tracer Data Sets** We downloaded the KP-Tracer data sets, including Cassiopeia-Hybrid trees, from Zenodo (`https://doi.org/10.5281/zenodo.5847462`).

##### 2.2 Cell Lineage Tree Reconstruction Software Availability and Commands

**Cassiopeia-Greedy.** We downloaded Cassiopeia-Greedy from Github (`https://github.com/YosefLab/Cassiopeia`; commit `9f272fc`). We ran Cassiopeia-Greedy with the following command:

```
python3 run_cassiopeia.py \
    -i [input character matrix] \
    -m [input mutation priors] \
    -o [output] \
    --method 1
```

where setting method option to 1 indicates greedy. In our experiments, we found that Cassiopeia-Greedy typically outputs a **binary tree**.

**Startle-ILP.** We downloaded Startle-ILP from Github (`https://github.com/raphael-group/startle`; commit `0bb0849`) and then built it from source with the following dependencies: gcc v11.2.0, BOOST v1.80, LEMON v1.3.1, IBM ILOG CPLEX Optimizer v12.7. We then ran Startle-ILP using the following command:

```
perl startle_ilp.pl \
    -o [output] \
    -m [input mutation priors] \
    -c [input character matrix] \
    --time-limit 86400
```

In our experiments, we found that Startle-ILP typically outputs a **non-binary tree**. We only ran Startle-ILP on the Startle simulated data sets. We attempted to run Startle-ILP on a few of the LAML simulated data sets (250 cells) and the two larger KP-Tracer data sets (3513_NT_T1_Fam and 3724_NT_All) by adding the option `--threads 16` to increase the number of threads from 1 to 16; however, Startle-ILP still did not complete within 24 hours (86400 seconds) on these data sets.

**Startle-NNI.** We downloaded the C++ version of Startle-NNI from Github (`https://github.com/raphael-group/startle`; commit `b0c02b5`). To run Startle-NNI, we first computed a neighbor-joining (NJ) tree, following the recommended command:

```
python3 nj.py [input character matrix ] --output [output]
```

where nj.py is available in the scripts directory (https://github.com/raphael-group/startle/blob/main/scripts/nj.py). We then ran Startle-NNI (C++) with the following command:

```
startle large \
   [input character matrix] \
   [input mutation priors] \
   [input starting tree computed with NJ] \
   --output [output]
```

In our experiments, we found that Startle-NNI (C++) always returned a **binary tree**. Because we were unable to reproduce results of the Startle study [9] with the newer C++ code, we also downloaded the older Python version of Startle-NNI from commit 0bb0849. We ran Startle-NNI (Python) with the following command:

```
python3 startle.py \
    -m [input mutation priors] \
    --iterations 250 \
    --mode infer \
    --threads 16 \
    --output [output] \
    [input starting tree computed with NJ] \
    [input character matrix]
```

We set the number of threads to 16 to speed up the computation because the Python version is much slower than the C++ version and our goal was to reproduce results from the Startle study, after failing to do so with the Startle-NNI (C++). Unlike Startle-NNI (C++), Startle-NNI (Python) has the user set the number of iterations—we used 250 as reported in the Startle study [9]. In our experiments, Startle-NNI (Python) typically returned a **non-binary tree** (it seemed like mutationless branches under the SH-model were being contracted; see Section 2.4).

**LAML.** We installed LAML [6] with "pip install" following the instructions on Github (https://github.com/raphael-group/LAML/tree/master). We ran LAML version 0.0.4 (commit 3598d14). On simulated data sets, we ran LAML with the following command:

```
run_laml \
    -c [input character matrix] \
    -p [input mutation priors] \
    -t [input starting tree] \
    -o [output prefix] \
    -v \
    --nInitials 1 \
    --topology_search
```

This is the same command used to analyze simulated data in the original LAML study (see Section S2.2.1 in the Supplementary Materials of [6]), except that the code has been renamed from run_problin.py and there is no longer an option --ultrametric in the code base (we assume this option is the default). In the original LAML study [6], the authors computed the starting tree with Startle-NNI (Python). We computed the starting tree with Startle-NNI (C++) because we wanted to use the latest version of the code (see Section 2.9 for details). We later realized we were unable to reproduce results with Startle-NNI (C++) for the Startle simulated data sets; however, the trends using Startle-NNI (C++) on the LAML simulated data sets and biological data sets still seem reasonable. In our experiments, we found that LAML returned a binary tree.

**PAUP*.** PAUP* is a widely used software package for phylogenetics and character parsimony, in particular. We downloaded PAUP* verson 4a168_centos64 from https://paup.phylosolutions.com. To run PAUP* as a heuristic for Star Homoplasy parsimony, we first "binarized" the character matrix (see [9] for description). As an example, the Startle input character matrix

```
,c0,c1
0,1,0
1,0,1
2,2,2
3,2,2
```

```
4,-1,2
5,-1,3
6,-1,3
7,-1,3
8,0,4
```

would be converted to the binarized matrix

```
#NEXUS
Begin data;
        Dimensions ntax=10 nchar=x;
        Format datatype=standard gap=-;
        Matrix
ROOT0
000000
ROOT1
000000
LEAF0
100000
LEAF1
001000
LEAF2
010100
LEAF3
010100
LEAF4
--0100
LEAF5
--0010
LEAF6
--0010
LEAF7
--0010
LEAF8
000001
        ;
End;
```

where we added two dummy leaves ROOT0 and ROOT1 of unedited states to the matrix so that we can root the tree at these leaves. We second convert the mutation priors in integer-valued positive weights by taking the negative log of the probability, multiplying by 100, and then rounding/converting to an integer. As an example, the Startle input mutation priors

```
character,state,probability
c0,1,0.90
c0,2,0.10
c1,1,0.01
c1,2,0.01
c1,3,0.80
c1,4,0.08
```

would be converted into a vector of integer-valued positive weights

```
10 229
461
461
22
252
```

We third execute a heuristic search under weighted Camin-Sokal parsimony using PAUP* with the following command:

```
paup4a168_centos64 [nexus file]
```

where the NEXUS file contains

```
#NEXUS
BEGIN PAUP;
set autoclose=yes warntree=no warnreset=no;
execute [input binarized character matrix];
```

```
outgroup ROOT0;
typeset myctype = irrev.up:1-N;
wtset mywtset vector = [weight vector];
assume typeset=myctype wtset=mywtset;
hsearch start=stepwise addSeq=random swap=None nreps=10 rseed=55555;hsearch start=1 swap=TBR
    nbest=500 rseed=12345;
rootTrees;
savetrees File=[output] root=yes trees=all format=newick;
END;
```

These commands tell PAUP* to

- Build a tree via "random taxon addition" (i.e., cells are put in a random order and then a tree is built by iteratively adding them to the tree so that the parsimony score is optimized).
- Repeated this process ten times and then the take best scoring tree found as the starting tree.
- Perform heuristic search from the starting tree using Tree Bisection and Reconnection (TBR) edit moves.
- Save the 500 best-scoring trees found during the search.

Because PAUP* does not score trees using floating-point weights or arithmetic, we recompute the Star Homoplasy parsimony score for each of the 500 best trees using the command from Section 2.3. The PAUP* tree is the first best scoring tree found in the output file from PAUP*. In our experiments, we found that the trees saved by PAUP* were typically **non-binary**. However, PAUP* is currently closed source and the details of the PAUP* heuristic search and its treatment of binary / non-binary trees are unavailable. Lastly, we computed the strict consensus of the trees that achieve the best score (out of the 500 saved) with the following command:

```
paup4a168_centos64 [nexus file]
```

where the NEXUS file contains

```
#NEXUS
BEGIN PAUP;
set maxtrees=510;
set autoclose=yes warntree=no warnreset=no;
gettrees file=[input best scoring trees];
contree all/strict=yes treefile=[output] format=newick;
END;
```

In our experiments, we found the PAUP* strict consensus (SC) tree was **non-binary**, which makes sense.

**Star-CDP.** Star-CDP is written in C++ with open source code available on Github (`https://github.com/molloy-lab/Star-CDP`). Star-CDP uses ASTRAL-III [13] to build the clade constraint set; we downloaded ASTRAL-III from Github (`https://github.com/smirarab/ASTRAL`; commit ec9844a). To run Star-CDP, we first ran PAUP* using the command above. We then ran Star-CDP (commit `c157857`) with the following command:

```
star-cdp \
    -i [input character matrix] \
    -m [input mutation priors] \
    -x [outgroup] \
    -t [trees for building search space, specifically the 500 best trees saved by PAUP*] \
    -rand [number of random solutions to generate] \
    -consensus \
    --output [output] \
    -nosupp \
    -XOUTG
```

where `-XOUTG` indicates that the outgroup specified with the `-x` option should be removed from the trees. We set the outgroup to be a "dummy" unedited cell to the matrix and replacing ROOT0 and ROOT1 in the PAUP* trees. This Star-CDP command returns files with the prefix `output`:

- [output]_number_of_sol.csv contains number of solutions (i.e., binary trees achieving optimal SH parsimony score) in the constrained search space
- [output]_strict_consensus.tre contains the **strict consensus** of solutions in the constrained search space
- [output]_majority_consensus.tre contains the **majority consensus** of solutions in the constrained search space (not used in our study)
- [output]_greedy_consensus.tre contains a **greedy consensus** of solutions in the constrained search space (not used in our study)
- [output]_one_sol.tre contains a single solution within the constrained search space (this one solution is biased because a clade with the smallest size is always selected when backtracking through the extended dynamic programming matrix)
- if the -rand N option is used, then [output]_random_sol_trees.tre will contain N random solutions generated by randomly selecting a clade at random when backtracking through the extended dynamic programming matrix

#### 2.3 Star Homoplasy (SH) Score

The Star Homoplasy (SH) parsimony scores involves floating-point arithmetic so avoid differences in precision across methods, we recomputed the SH parsimony score by running Startle-NNI (C++) with the following command:

```
startle small \
    [input character matrix] \
    [input mutation priors] \
    [input tree to score] \
    --output [output]
```

#### 2.4 Star Homoplasy (SH) Contraction

In the Startle study [9], tree error metrics are computed **after** contracting mutationless branches under the Star Homoplasy (SH) model. To compute the SH-contraction of $T$, we contract each non-terminal branch $u \mapsto v$ in $T$ for which the SH-labeling of $u$ and $v$ are identical and thus there are no substitutions occurring on the edge. Recall that the SH-labeling assigns states to the internal vertices of $T$ given a character matrix $\mathcal{C}$ and under the assumption that each site in $\mathcal{C}$ evolves under the SH model. The SH-labeling does not depend on the mutation prior; see main text for details. For some data sets, we first compute the SH-contraction of $T^*$ and $\hat{T}$ and then compute error and accuracy metrics according to the equations in Section 2.5.

#### 2.5 Tree Accuracy and Error Metrics

For simulated data sets, we can then compare the estimated tree $\hat{T}$ to the true (i.e., model) tree $T^*$ used to simulated the data (note that both of these trees are **rooted**). Given $T^*$ and $\hat{T}$, we compute the following quantities:

- $TP(T^*, \hat{T})$ := number of **true positive (TP)** branches, i.e., number of clades in both $T^*$ and $\hat{T}$
- $FP(T^*, \hat{T})$ := number of **false positive (FP)** branches, i.e., number of clades in $\hat{T}$ but not $T^*$
- $FN(T^*, \hat{T})$ := number of **false negative (FP)** branches, i.e., number of clades in $T^*$ but not in $\hat{T}$
- $NI(\hat{T})$ := number of clades in the tree $\hat{T}$
- $NI(T^*)$ := number of clades in the tree $T^*$

**Code.** The code for computing error and accuracy metrics is available on Github (`https://github.com/molloy-lab/star-study/blob/main/B_scripts/compare_two_rooted_trees_under_star.py`). We use the following command:

```
python3 compare_two_rooted_trees_under_star.py \
    -t1 [tree 1] \
    -t2 [tree 2] \
    -c1 [0,1] \
    -c2 [0,1] \
    -ex1 [0,1] \
    -ex2 [0,1] \
    -m [input character matrix] \
    -t2p [Path to write the trees after SH-contraction]
```

where options `-c1` and `-c2` indicate whether to compute the SH-contraction for the the trees given in options `-t1` and `-t2`, respectively (1 indicates the SH-contraction should be computed; 0 indicates otherwise). Relatedly, the options `-ex1` and `-ex2` indicate that the trees given in options `-t1` and `-t2`, respectively, should be written to a file after taking their SH-contraction is computed (i.e., it saves the SH-contracted trees for later use). If tree 1 is the true tree $T^*$ and tree 2 is estimated tree $\hat{T}$, then code returns the number of leaves shared in both trees, $NI(T^*)$, $NI(\hat{T})$, $FN(T^*, \hat{T})$, $FP(T^*, \hat{T})$, and $TP(T^*, \hat{T})$.

**Accuracy Metrics.** From the quantities above, we compute accuracy metrics, specifically precision, recall, and f1-score.

$$precision(T^*, \hat{T}) := \frac{TP(T^*, \hat{T})}{NI(\hat{T})} = \frac{TP(T^*, \hat{T})}{TP + FP} \tag{3}$$

$$recall(T^*, \hat{T}) := \frac{TP(T^*, \hat{T})}{NI(T^*)} = \frac{TP(T^*, \hat{T})}{TP + FN} \tag{4}$$

$$f1(T^*, \hat{T}) := \frac{2}{precision(T^*, \hat{T})^{-1} + recall(T^*, \hat{T})^{-1}} \tag{5}$$

**Error Metrics.** From the quantities above, we also compute error metrics. The standard error metric for phylogenetic trees is the Robinson-Foulds (RF) distance [8]:

$$RF(T^*, \hat{T}) := FP(T^*, \hat{T}) + FN(T^*, \hat{T}). \tag{6}$$

where we define the FP and FN in terms of clades (rather than bipartitions) because the trees are rooted [1]. In systematic studies, both $\hat{T}$ and $T^*$ are typically binary, in which case, the normalized RF distance equals the false negative rate (FNR)

$$FNR(T^*, \hat{T}) := \frac{FN(T^*, \hat{T})}{NI(T^*)} \tag{7}$$

and the false positive rate (FPR)

$$FPR(T^*, \hat{T}) := \frac{FP(T^*, \hat{T})}{NI(\hat{T})}. \tag{8}$$

In lineage tracing studies, typically one or both $T^*$ and $\hat{T}$ are non-binary, especially because mutationless branches are often contracted before computing error or accuracy metrics. For non-binary trees, different code bases will perform normalization in different ways. Additionally, some code bases that normalize the RF distance considering the expected distances between a random pair of trees (these types of calculations typically assume both trees are binary). To avoid confusion or other issues, we report the unnormalized RF distance (Eq. 6).

#### 2.6 Handling cells with missing data for KP-Tracer data sets

**Pruning.** We also use the pruning technique technique as introduced in [9]. We first binarize the input matrix $C$ by applying one-hot encoding for all observed states on every site to get matrix $C^{bin}$. Then we define the partial order as the following $i \succcurlyeq i'$ if and only if for all column $j$ in $C^{bin}$, either $C^{bin}_{i,j} = C^{bin}_{i',j}$ or $C^{bin}_{i,j} = C^{bin}_{i',j}$ (i.e., cell $i'$ precedes cell $i$ if we can impute the missing entries of $i'$ to obtain $i$). We then say that a cell $i$ is maximal if there is no other cell $i' \neq i'$ such that $i' \succcurlyeq i$. In the pruning process, we first identify all maximal cells making them the seed of an equivalence class up to missing (`eq-class-mis`). We put all remaining cells in their respective equivalence class based on the seeds; these cells are then removed or pruned the from the character matrix or tree of interest. Code is here: `https://github.com/molloy-lab/star-study/blob/main/B_scripts/KPTracer-Data-Full_deprune_deduplicate/a_prune_all_character_matrix.py`.

**Unpruning.** The unpruning process transforms trees with leaves labeled by seeds of `eq-class-mis` into trees on the complete leaf set. This is achieved by subdividing each edge incident to a cell in `eq-class-mis` with new vertex and then attaching all other cells in the same equivalence class to this new vertex, resulting in a polytomy if the equivalence class has size 3 or greater. Code is here: `https://github.com/molloy-lab/star-study/blob/main/B_scripts/KPTracer-Data-Full_deprune_deduplicate/startle_nni/b1_deprune.py`. We use the two functions, from_newick_get_nx_tree and tree_to_newick_eq_classes for deprune in the utilities python lib used in [9].

**Deduplicating.** We also use the deduplicate process introduced in [6]. We first loop over the cells in the character matrix, saving those with unique sequence that has yet to be observed; these cells are part referred in the set "seed for equivalence class up to exact match" (`eq-class-ex`). We remove all cells that are not seeds in `eq-class-ex` from the character matrix or tree of interest. Code is here: `https://github.com/molloy-lab/star-study/blob/main/B_scripts/KPTracer-Data-Full_deduplicate/a_deduplicate_character_matrix.py`. Note that we modify the utilities python lib in [9] but one line code to check whether two cells have exactly same sequence (see function compute_equivalence_classes_up_to_exact in `https://github.com/molloy-lab/star-study/blob/main/B_scripts/KPTracer-Data-Full_deprune_deduplicate/utilities.py`).

#### 2.7 Cell lineage tree reconstruction pipelines for KP-Tracer data sets

##### 2.7.1 Pipelines based on deduplicated data sets

The deduplicate data analysis pipelines have the following steps:

1. **De-duplicate each character matrix** (Section 2.6).
2. **Run the cell lineage tree reconstruction methods** Startle-NNI (C++), Star-CDP, and PAUP* given the **deduplicated matrix** from step (1) as input.
3. Use LAML to estimate branch lengths and improve the topology of the cell lineage trees from step (2). See pipeline descriptions below.
4. Analyze the cell lineage trees from step (3) in terms of SH parsimony score, inferred migrations (Section 2.11), etc.

**Analysis pipeline #1a (de-duplicate).** We follow the de-duplicate pipeline above and then do not change the topology of the reconstructed cell lineage trees. However, we do estimate branch lengths on the binary cell lineage trees (i.e., the Startle-NNI (C++), Star-CDP-Rand, and Star-CDP-Bias trees) using LAML (Section 2.9). Note that the Cassiopeia-Hybrid trees published for the KP-Tracer data sets [12] have been re-analyzed in both the Startle [9] and LAML [6] studies. Because the Cassiopeia-Hybrid was run on the complete character matrix, rather than the deduplicated matrix, we restricted it the same leaf set as our other reconstructed cell lineage trees by removing all duplicate cells. This same approach was used in the LAML study [6].

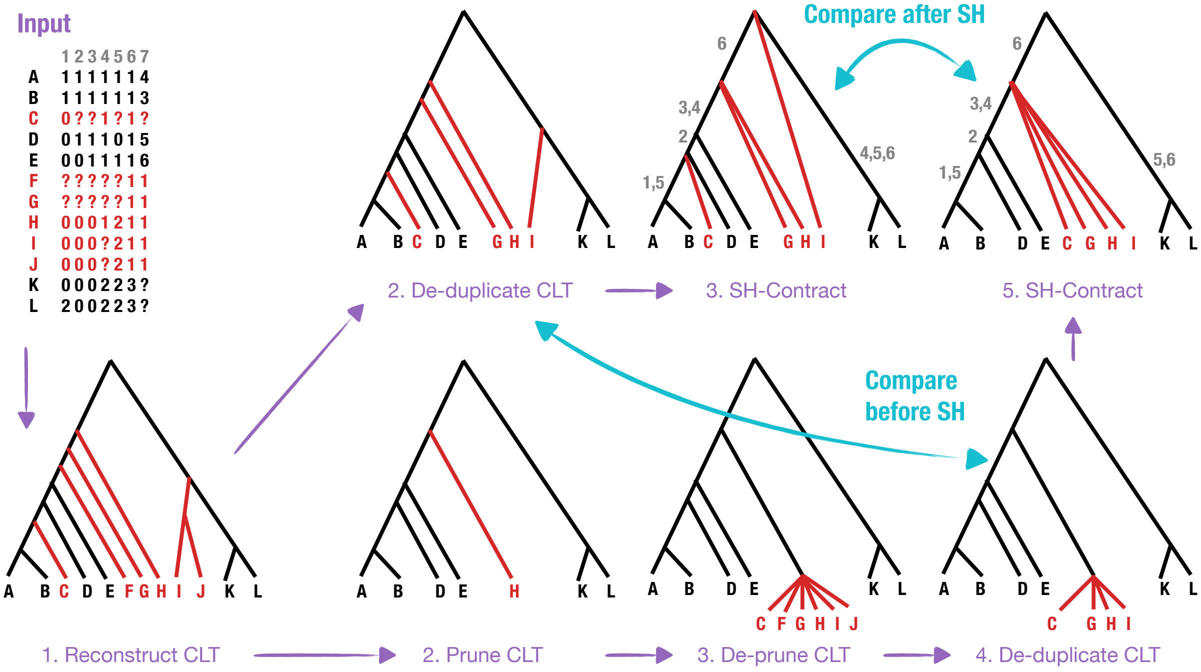

Figure S2: **Data processing pipelines and SH contraction (continued).** This figure shows the two data processing pipelines for the Cassiopeia published tree, which was estimated on the complete data set. Pipeline #1) is the de-duplicate only pipeline (result in step 2), and Pipeline #2) is the prune, deprune, deduplicate (PDD) pipeline (result in step 4). It also shows the SH-contraction of the results, in which mutionless edges under the SH model are contracted. Characters numbers are drawn on the branches in grey where there must be a substitution. Trees can be compared before or after SH contraction.

**Analysis pipeline #1b (de-duplicate + LAML refine).** We follow the de-duplicate above pipeline and then seek to improve the topology of the reconstructed cell lineage trees by using LAML refine. Specifically, for each reconstructed cell lineage tree, we contracted mutationless branches under the SH model (Section 2.4). We then ran LAML to resolve polytomies and estimate branch lengths (Section 2.9).

**Analysis pipeline #1c (de-duplicate + LAML search).** We follow the de-duplicate pipeline above and then seek to improve the topology of the reconstructed cell lineage trees by using LAML search. Specifically, we ran LAML with each of the reconstructed trees (no contractions) as its starting tree (Section 2.9). LAML returns a binary tree with estimated branch lengths.

##### 2.7.2 Analysis pipelines based on pruned data sets.

The pruned data analysis pipelines have the following steps:

1. **Prune each character matrix** (Section 2.6.

2. **Run the cell lineage tree reconstruction methods** Startle-NNI (C++), Star-CDP, and PAUP* given the **pruned matrix** from step (1) as input. (Section 2.2).

3. **De-prune the cell lineage trees** from step (2) (Section 2.6).

4. **De-duplicate the cell lineage trees** from step (3).

5. Use LAML to estimate branch lengths and improve the topology of the cell lineage trees from step (4) given the de-duplicated character matrix (Section 2.7.1). See pipeline descriptions below.

6. Analyze the cell lineage trees from step (5) in terms of SH parsimony score, inferred migrations (Section 2.11), etc.

We refer to the first four steps as **prune, de-prune, and de-duplicate (PDD)**.

**Analysis pipeline #2a (PDD).** We follow the PDD pipeline above and then do not change the topology of the reconstructed cell lineage trees. However, we do estimate branch lengths on the binary cell lineage trees (i.e., the Startle-NNI (C++), Star-CDP-Rand, and Star-CDP-Bias trees) using LAML (Section 2.9). Note that the published trees associated with the KP-Tracer data sets [12] were computed using Cassiopeia-Hybrid on the complete character matrix. These published trees have been re-analyzed in both the Startle [9] and LAML [6] studies. In our study, we restricted the published trees to the same leaf set as our other reconstructed cell lineage trees, by pruning the published trees, de-pruning the published trees, and then de-duplicating the published trees. This PDD analysis of the published trees was not performed in either the Startle [9] or LAML [6] studies.

**Analysis pipeline #2b (PDD + LAML refine).** We follow the PDD above pipeline and then seek to improve the topology of the reconstructed cell lineage trees by using LAML refine. Specifically, for each reconstructed cell lineage tree, we contracted mutationless branches under the SH model (Section 2.4). We then ran LAML to resolve polytomies and estimate branch lengths (Section 2.9).

**Analysis pipeline #2c (PDD + LAML search).** We follow the PDD pipeline above and then seek to improve the topology of the reconstructed cell lineage trees by using LAML search. Specifically, we ran LAML with each of the reconstructed trees (no contractions) as its starting tree (Section 2.9). LAML returns a binary tree with estimated branch lengths.

#### 2.8 Properties of KP-Tracer data sets after pruning or deduplicating

Table S1: **Equicalence classes (ECs) for KP-Tracer data sets after pruning or deduplicating.** Column 1 is the number of cells in the original data sets. Column 2 is the number of cells after creating equivalence classes for deduplicating (i.e., equivalent including missing data) and then removing all but one cell per EC. Column 3 is the number of cells after creating ECs for pruning (i.e., equivalent up to missing data) and then removing all but one cell per EC. Recall that the goal of pruning is to create maximal subsets of leaves such that if they are attached as a polytomy, it does not increase the SHP score more than having one element at that location in the tree (i.e., these cells are equivalent up to missing states). Column 5 is the number of pruning ECs computed for the full data set with more than 2 elements (otherwise depruning just produces a cherry); column 6 is the mean EC size (i.e., number of cells per EC, averaged over these ECs). Column 7 is the number of deduplicating ECs computed for the full data set with more than 2 elements; column 8 is the mean EC size. Column 9 is the number of pruning ECs computed for the deduplicated data set with more than 2 elements; column 10 is mean EC size.

| Tumor Data | # of cells in total | # cells after dedup | # cells after prune | # EC prune | Size EC prune | # EC dedup | Size EC dedup | # EC PDD | Size EC PDD |
|---|---|---|---|---|---|---|---|---|---|
| 3724_NT_All | 21108 | 1461 (93%) | 1207 (17%) | 275 | 72.81 | 324 | 61.09 | 30 | 5.9 |
| 3513_NT_T1_Fam | 1227 | 101 (92%) | 86 (14%) | 14 | 81.79 | 20 | 56.6 | 5 | 4.0 |
| 3515_Lkb1_T1_Fam | 1878 | 1013 (46%) | 878 (13%) | 80 | 12.4875 | 73 | 11.86 | 7 | 5.29 |

#### 2.9 Branch length estimation for KP-Tracer data sets

We used LAML to resolve any polytomies, with the following command:

```
run_laml \
    -c [input character matrix] \
    -p [input mutation priors] \
    -t [input tree] \
    -o [output] \
    -v \
    --nInitials 1 \
    --timescale 6 \
    --resolve_search \
    --parallel
```

where the input tree was computed with different methods (remove `--parallel` and `--resolve_search` if computing branch lengths on fixed topology). We also performed a full tree search with LAML, with the following command:

```
run_laml \
    -c [input character matrix] \
    -p [input mutation priors] \
    -t [input starting tree] \
    -o [output prefix] \
    -v \
    --nInitials 1 \
    --topology_search \
    --maxIters 2500 \
    --parallel \
    --timescale 6
```

This is the same command used to analyze the KP-Tracer data in the original LAML study (see Section S2.4 in the Supplementary Materials of [6]), except that the code has been renamed from `run_problin.py` and there is no longer an option `--ultrametric` in the code base (we assume this option is the default).

#### 2.10 Runtime

Computational experiments were performed on compute nodes outfitted with 32 AMD EPYC-7313 cores and 2TB of RAM. We ran all methods were run with 1 thread, 48 GB of memory, and a maximum wall clock time of 24 hours, except for Startle-NNI (Python), which was given access to 16 threads. When collecting runtime data, requested exclusive access of these compute nodes via slurm scheduler.

**Simulated data sets.** Runtime results for simulated data are shown in Figures S12 and S9. The runtime of Startle-NNI (both Python and C++ versions) includes the time to compute the starting tree with Neighbor Joining (NJ). The runtime of LAML includes the time to compute the Startle-NNI (C++) tree, which in turn includes the time to compute the NJ tree. The runtime of Star-CDP includes the time to run PAUP*.

**KP-Tracer data sets.** For all six analysis pipelines, we had a 48 GB memory limit and 24 hours wall-clock time limit. All methods were run with 1 thread, except LAML, which was given access to 16 threads via the SLURM system. LAML did always complete within 24 hours; however, the LAML log file contains the best tree found for after every 10 NNI iterations. If LAML failed to complete, we took the best tree from the log file. An issue here is that the parameters may not have converged. Notably, LAML failed to converge on 3515_LKb1_T1_Fam on all given start trees.

#### 2.11 Migration analysis for KP-Tracer data sets

We also analyzed the migration history of all estimated trees on all pipelines following the prior studies [9, 3], using the adapted Sankoff algorithm. However, many possible equally optimal labelings for internal nodes might exist in a tree, in order to reduce the number of cross-metastasis seeding events and be comparable to the results in [9, 6], we do the following. We first map the primary tumor and all metastasis to an integer guaranteeing the primary tumor always maps to the least integer in a metastasis family and for different tumor cells in the same metastasis the integers they map to will guarantee that the tumor will small tumor ID is always less than the tumor with a larger ID(See `https://github.com/molloy-lab/star-study/blob/main/B_scripts/KPTracer-Data-Full_deprune_deduplicate/c_parse_site_labeling.py`). Then after getting the parsimonious migrations cost by the Sankoff algorithms, we perform the top-down traversal to get the labeling and for all internal node we always pick up the optimal label mapping to the smallest integer in the set of all integers that optimal labeling mapping to. This guarantees us to acquire the optimal labeling maximizing the number of nodes labeled by the primary tumor with a single top-down traversal as in [9, 6]. We implement these algorithms in the two functions count_migration and top_down (see example in `https://github.com/molloy-lab/star-study/blob/main/B_scripts/KPTracer-Data-Full_deduplicate/paup/f_compute_migration.py`)

### 3 Supplemental Results

#### 3.1 Supplemental results on Startle simulated data sets

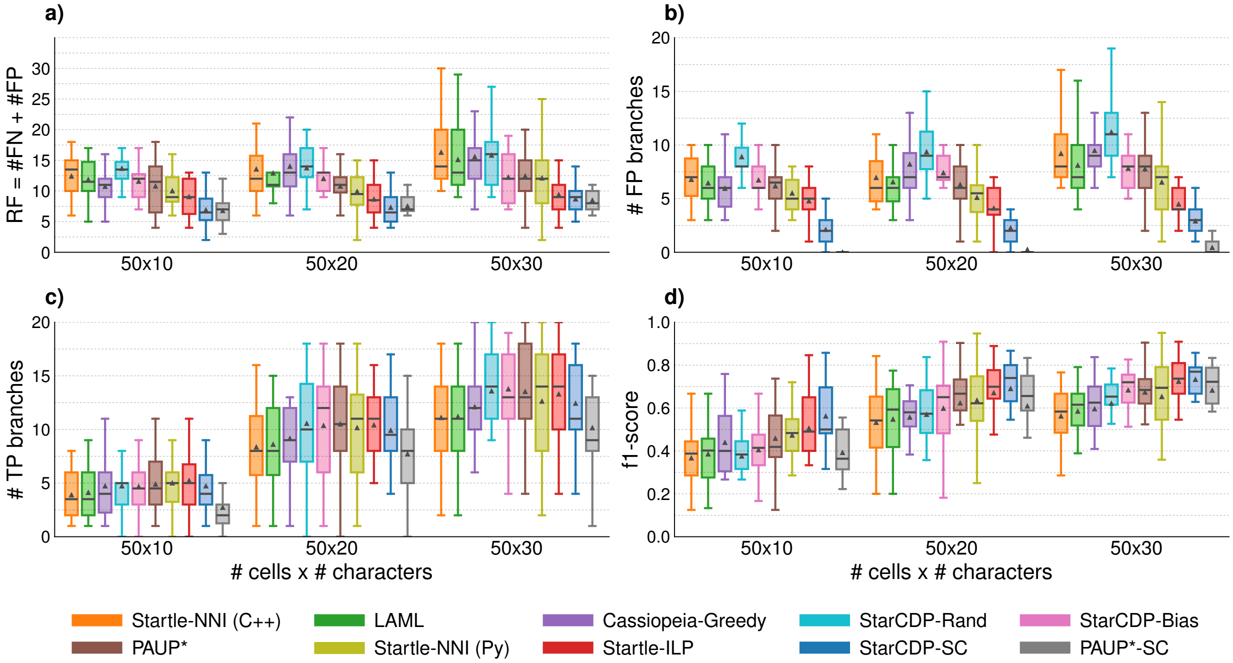

Figure S3: **Tree error and accuracy for Startle simulated data sets with 50 cells** after contracting mutationless branches in both true and estimated trees. Subplots **(a)** and **(b)** show error metrics (lower is better). Subplots **(c)** and **(b)** show accuracy metrics (higher is better). Triangles and bars are means and medians across replicate data sets, respectively.

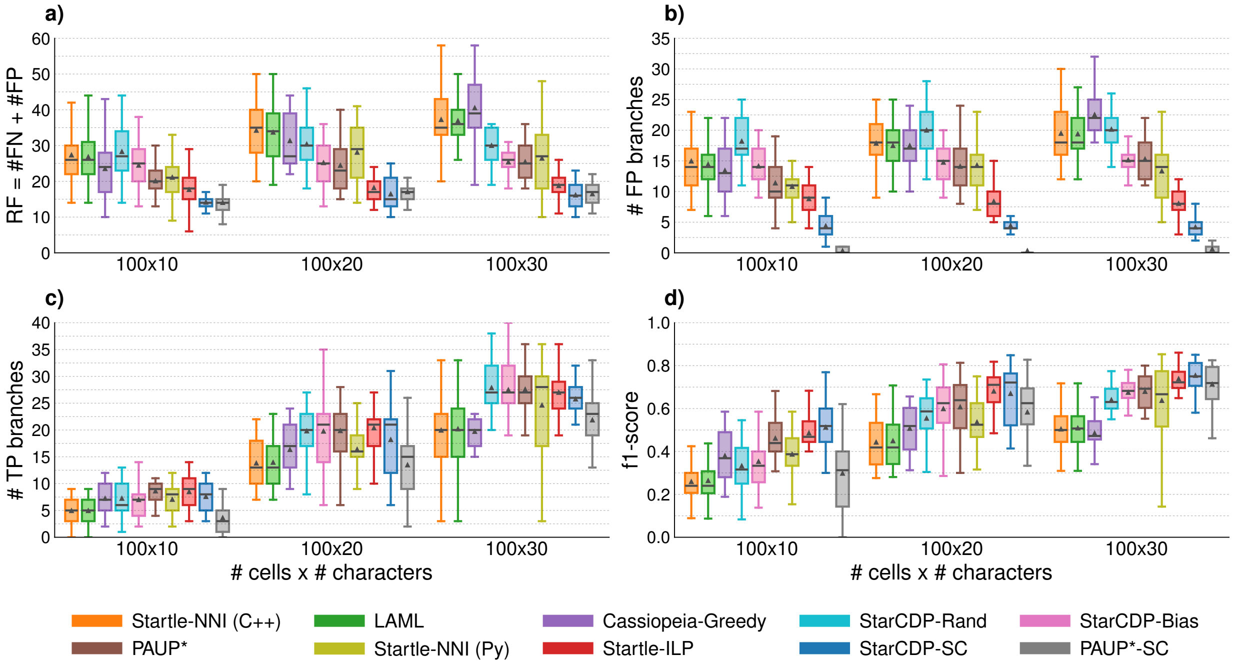

Figure S4: **Tree error and accuracy for Startle simulated data sets with 100 cells** after contracting mutationless branches in both true and estimated trees.

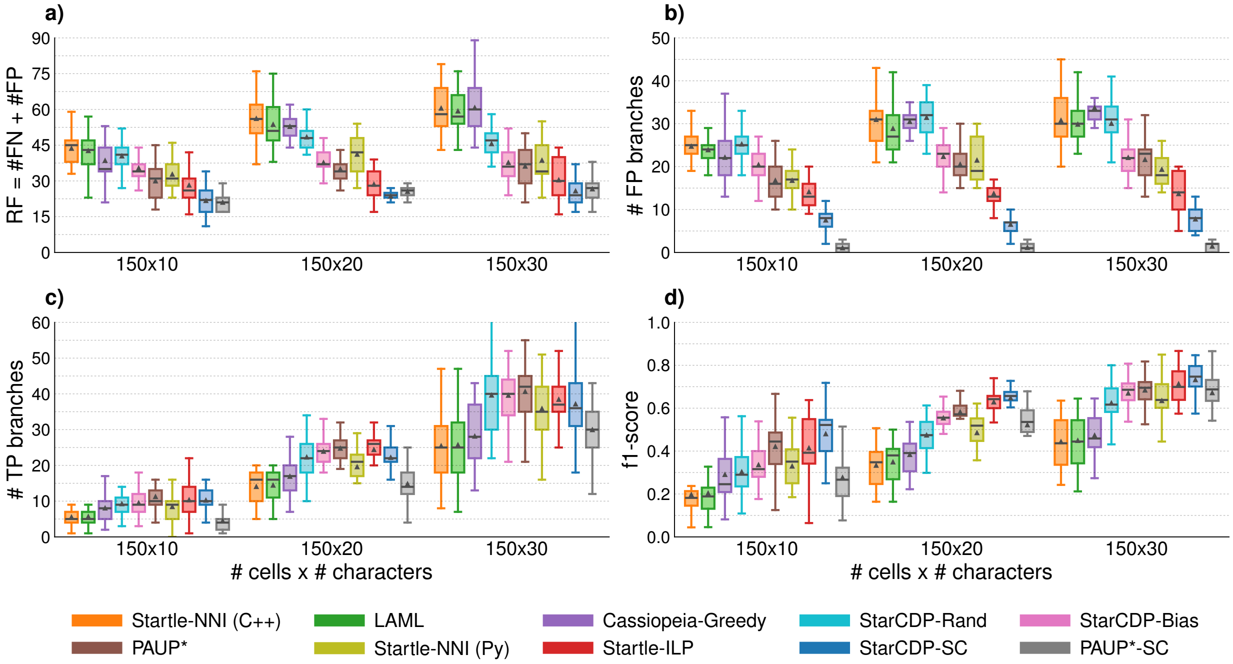

Figure S5: **Tree error and accuracy for Startle simulated data sets with 150 cells** after contracting mutationless branches in both true and estimated trees.

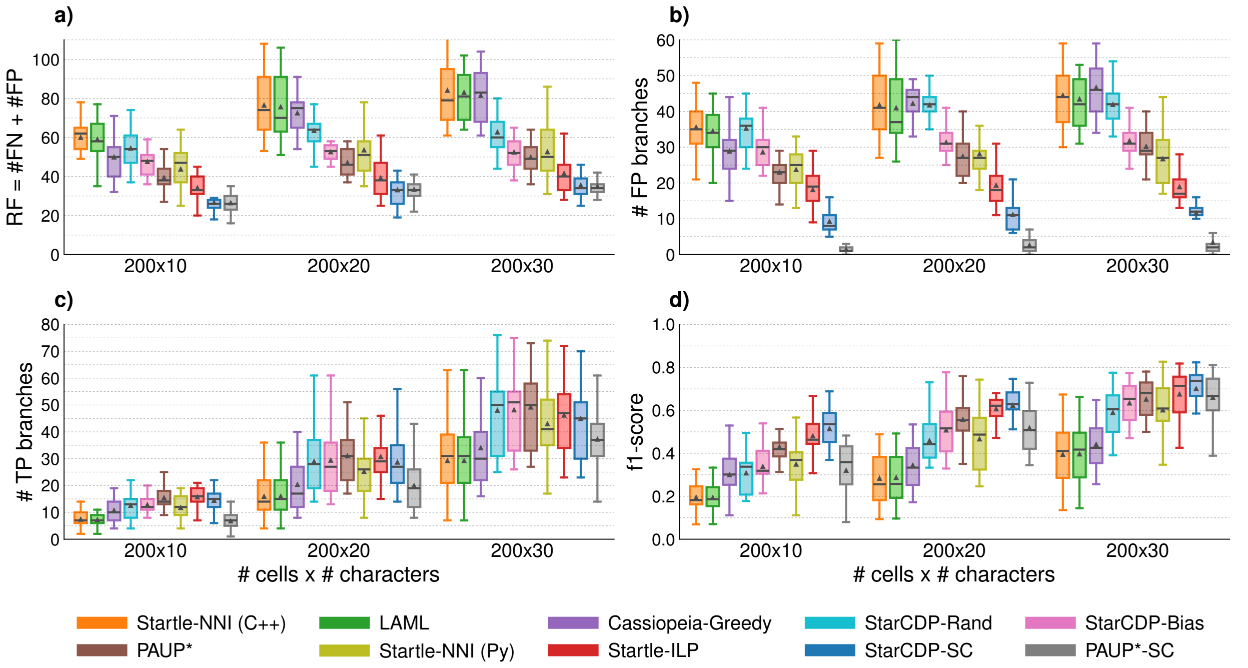

Figure S6: **Tree error and accuracy for Startle simulated data sets with 200 cells** after contracting mutationless branches in both true and estimated trees.

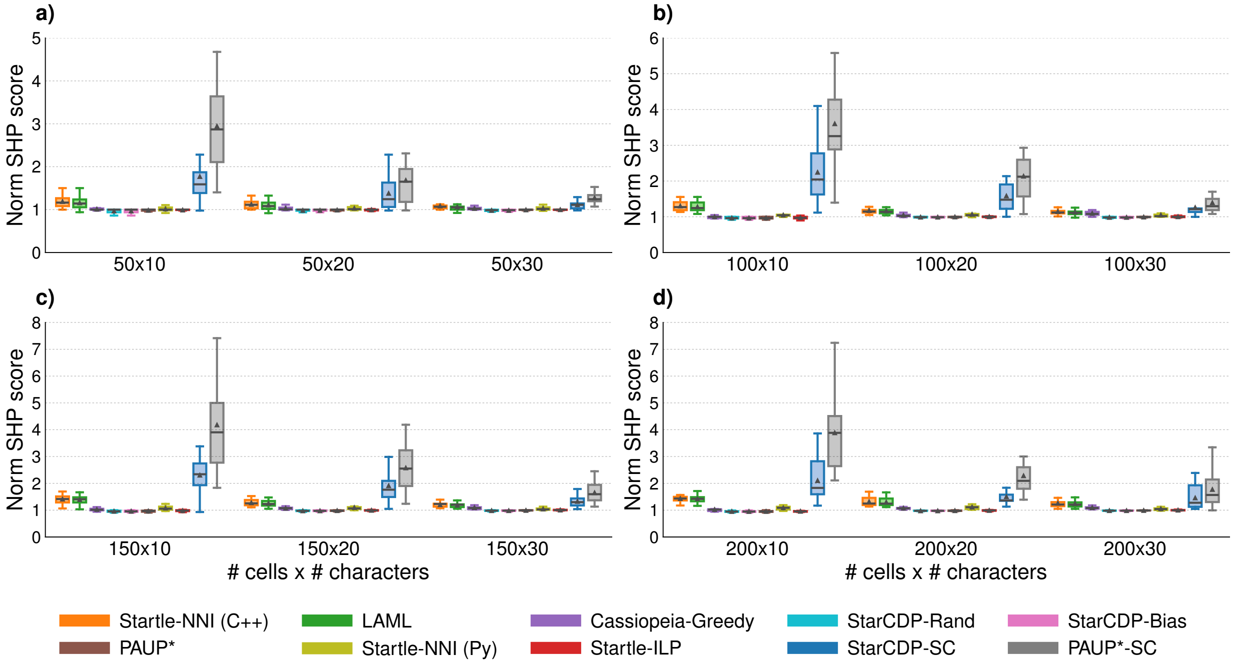

Figure S7: **Normalized SH parsimony score for Startle simulated data sets.** Subplots **(a)**, **(b)**, **(c)**, **(d)** show data sets with 50, 100, 150, and 200 cells, respectively. Triangles and bars are means and medians across replicate data sets, respectively.

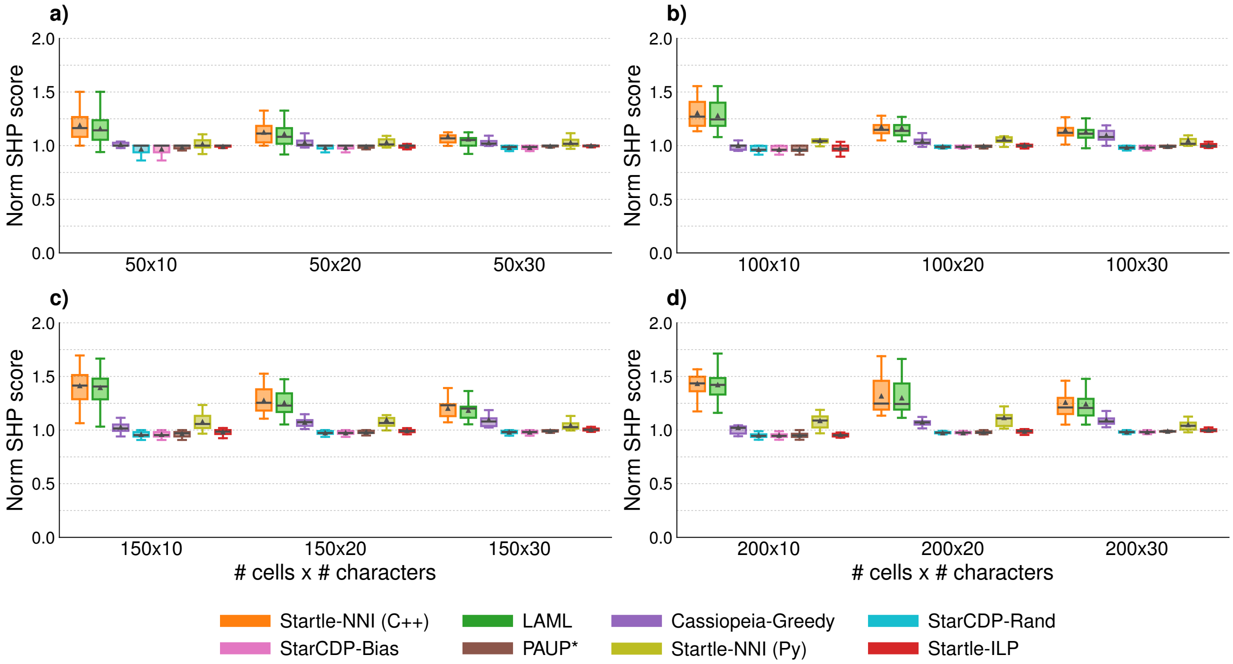

Figure S8: **Normalized SH parsimony score for Startle simulated data sets excluding PAUP* and Star-CDP consensus trees.** Same as Figure S7 but excludes consensus trees to zoom in on $y$-axis.

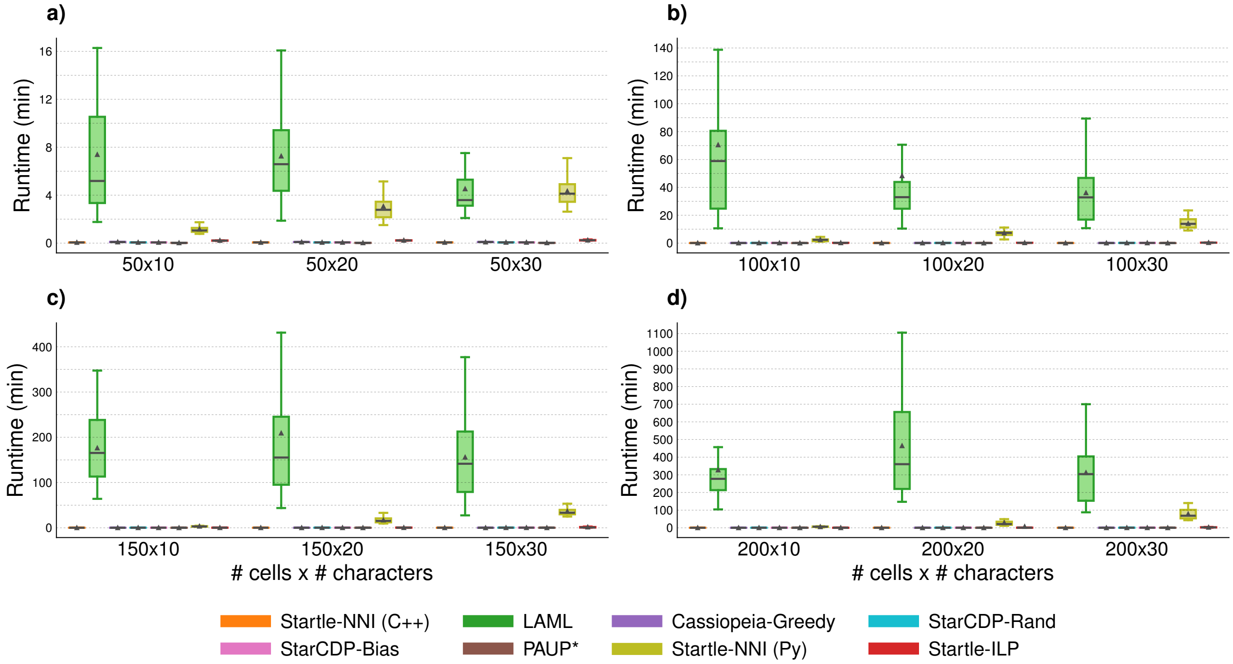

Figure S9: **Runtime (in minutes) for Startle simulated data sets.** Subplots **(a)**, **(b)**, **(c)**, **(d)** show data sets with 50, 100, 150, and 200 cells, respectively. Startle-NNI (Python) was run with 16 threads; all other methods were run with 1 thread. Runtime for Star-CDP includes the time to compute consensus trees (i.e., the Star-CDP-SC tree). Runtime for PAUP*-SC is not shown because it was negligible compared to the time to run the heuristic search.

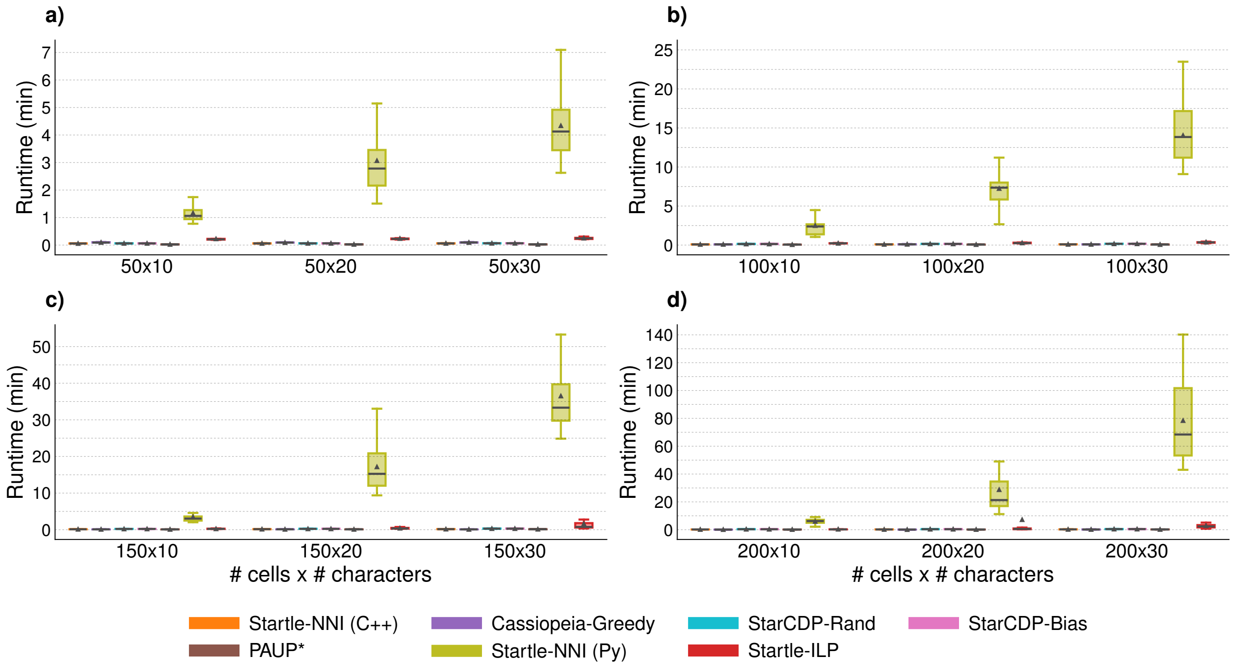

Figure S10: **Runtime (in minutes) for Startle simulated data sets excluding LAML.** Same as Figure S9 but excludes LAML to zoom in on $y$-axis.

Table S2: **Tree error for Startle simulated data sets.** Mean ± standard deviations are across replicates for each method. Tree error merics were computed after SH-contraction of both the true and estimated trees.

| # of cells | # of chars | Startle NNI (C++) | LAML | Cassiopeia Greedy | StarCDP Rand | StarCDP Bias | PAUP* | Startle NNI (Py) | Startle ILP | StarCDP SC | PAUP* SC |
|---|---|---|---|---|---|---|---|---|---|---|---|
| *RF distance = FN + FP* | | | | | | | | | | | |
| 50 | 10 | 12.4 ± 3.4 | 11.8 ± 3.4 | 10.7 ± 3.1 | 13.7 ± 2.7 | 11.6 ± 2.8 | 10.8 ± 4.2 | 10.0 ± 2.8 | 9.1 ± 3.2 | 6.9 ± 3.0 | **6.8 ± 2.2** |
| 50 | 20 | 13.6 ± 5.6 | 12.9 ± 4.9 | 14.0 ± 4.8 | 13.8 ± 3.7 | 12.0 ± 3.2 | 10.8 ± 3.0 | 9.9 ± 4.2 | 8.6 ± 3.1 | **7.3 ± 2.7** | 7.5 ± 1.7 |
| 50 | 30 | 16.3 ± 5.3 | 15.1 ± 5.4 | 15.6 ± 5.5 | 15.8 ± 4.8 | 12.2 ± 3.7 | 12.4 ± 3.6 | 12.1 ± 5.7 | 9.4 ± 3.4 | 8.7 ± 2.5 | **8.5 ± 2.2** |
| 100 | 10 | 27.3 ± 8.1 | 26.8 ± 8.6 | 23.5 ± 7.9 | 28.3 ± 7.3 | 24.5 ± 6.6 | 20.1 ± 6.5 | 21.1 ± 5.5 | 17.7 ± 5.4 | 14.1 ± 4.4 | **14.0 ± 3.7** |
| 100 | 20 | 34.2 ± 8.0 | 33.7 ± 8.0 | 31.3 ± 7.4 | 30.4 ± 6.3 | 25.2 ± 6.3 | 24.4 ± 6.8 | 28.1 ± 7.9 | 18.2 ± 4.6 | **16.4 ± 4.6** | 17.0 ± 3.4 |
| 100 | 30 | 37.2 ± 8.6 | 36.9 ± 8.7 | 40.5 ± 8.4 | 30.0 ± 5.2 | 25.4 ± 3.8 | 25.4 ± 5.9 | 26.4 ± 10.5 | 18.8 ± 4.0 | **16.1 ± 3.5** | 16.5 ± 3.3 |
| 150 | 10 | 43.6 ± 8.5 | 42.7 ± 8.0 | 38.5 ± 9.0 | 40.4 ± 7.3 | 35.4 ± 5.9 | 30.0 ± 7.6 | 32.8 ± 6.8 | 28.2 ± 7.6 | 21.7 ± 5.8 | **21.0 ± 3.6** |
| 150 | 20 | 56.1 ± 10.4 | 53.7 ± 9.8 | 52.8 ± 5.3 | 48.4 ± 6.6 | 37.7 ± 5.9 | 35.0 ± 5.2 | 41.1 ± 8.4 | 28.6 ± 6.2 | **23.6 ± 3.1** | 25.6 ± 3.8 |
| 150 | 30 | 60.5 ± 10.1 | 59.3 ± 8.5 | 60.7 ± 11.5 | 45.6 ± 8.3 | 37.7 ± 7.6 | 36.1 ± 8.1 | 38.6 ± 8.8 | 30.4 ± 9.2 | **25.8 ± 5.8** | 26.6 ± 5.4 |
| 200 | 10 | 60.0 ± 9.9 | 59.0 ± 9.8 | 49.9 ± 11.9 | 54.4 ± 9.0 | 47.6 ± 7.4 | 39.3 ± 7.5 | 43.7 ± 11.1 | 34.1 ± 9.0 | 26.6 ± 5.6 | **26.4 ± 4.6** |
| 200 | 20 | 76.5 ± 15.8 | 75.6 ± 16.7 | 72.5 ± 9.1 | 63.5 ± 7.8 | 52.6 ± 7.4 | 47.0 ± 9.1 | 53.5 ± 13.3 | 39.2 ± 10.6 | **33.1 ± 8.5** | 33.3 ± 5.1 |
| 200 | 30 | 84.0 ± 16.2 | 83.0 ± 15.2 | 81.5 ± 13.1 | 62.7 ± 10.4 | 52.4 ± 8.1 | 49.8 ± 8.5 | 52.6 ± 14.0 | 41.5 ± 10.0 | 35.5 ± 5.9 | **35.0 ± 5.5** |
| *Number of false negative (FN) branches/clades* | | | | | | | | | | | |
| 50 | 10 | 5.6 ± 1.8 | 5.4 ± 1.6 | 4.8 ± 1.6 | 4.8 ± 1.4 | 4.8 ± 1.5 | 4.6 ± 2.0 | 4.5 ± 1.7 | **4.3 ± 1.8** | 4.8 ± 2.2 | 6.8 ± 2.2 |
| 50 | 20 | 6.6 ± 2.9 | 6.4 ± 2.7 | 5.8 ± 2.3 | **4.4 ± 1.5** | 4.6 ± 1.6 | **4.4 ± 1.2** | 4.8 ± 2.2 | 4.6 ± 1.5 | 5.0 ± 1.3 | 7.2 ± 1.7 |
| 50 | 30 | 7.1 ± 3.2 | 7.0 ± 3.1 | 6.1 ± 2.6 | 4.6 ± 2.1 | **4.4 ± 1.9** | 4.7 ± 1.7 | 5.6 ± 2.9 | 4.9 ± 2.1 | 5.8 ± 2.0 | 8.0 ± 2.3 |
| 100 | 10 | 12.4 ± 3.6 | 12.4 ± 3.6 | 10.1 ± 3.7 | 10.1 ± 3.8 | 10.3 ± 3.8 | **8.7 ± 3.4** | 10.3 ± 3.0 | 8.9 ± 3.0 | 9.8 ± 3.1 | 13.7 ± 3.6 |
| 100 | 20 | 16.4 ± 4.3 | 16.2 ± 4.2 | 13.9 ± 4.0 | 10.4 ± 2.8 | 10.4 ± 3.3 | 10.3 ± 3.1 | 13.8 ± 4.2 | **9.8 ± 2.3** | 12.0 ± 3.9 | 16.7 ± 3.5 |
| 100 | 30 | 17.8 ± 4.4 | 17.5 ± 4.4 | 18.0 ± 4.3 | **9.9 ± 3.0** | 10.2 ± 2.5 | 10.1 ± 3.1 | 13.1 ± 5.4 | 10.7 ± 2.2 | 12.0 ± 3.0 | 15.9 ± 3.0 |
| 150 | 10 | 18.9 ± 4.0 | 18.8 ± 3.9 | 16.4 ± 3.8 | 15.2 ± 3.0 | 15.0 ± 2.8 | **13.2 ± 3.7** | 16.0 ± 3.7 | 14.1 ± 4.0 | 14.2 ± 3.4 | 20.0 ± 3.7 |
| 150 | 20 | 25.2 ± 4.5 | 24.8 ± 4.3 | 22.3 ± 3.4 | 17.0 ± 3.5 | 15.3 ± 3.0 | **14.5 ± 2.3** | 19.7 ± 3.7 | 14.9 ± 3.4 | 17.0 ± 2.3 | 24.4 ± 4.0 |
| 150 | 30 | 29.7 ± 4.3 | 29.4 ± 4.5 | 27.0 ± 6.4 | 15.5 ± 4.6 | 15.6 ± 4.0 | **14.5 ± 3.7** | 19.3 ± 5.2 | 16.7 ± 4.7 | 18.0 ± 4.0 | 25.2 ± 5.1 |
| 200 | 10 | 24.4 ± 4.6 | 24.5 ± 4.5 | 21.0 ± 4.8 | 19.2 ± 4.2 | 19.0 ± 3.7 | 16.3 ± 3.4 | 20.0 ± 5.6 | **16.0 ± 4.5** | 17.3 ± 4.0 | 25.0 ± 4.5 |
| 200 | 20 | 34.7 ± 7.1 | 34.7 ± 7.2 | 30.3 ± 5.5 | 21.7 ± 4.2 | 21.2 ± 4.3 | **19.6 ± 4.7** | 25.4 ± 6.4 | 19.9 ± 5.2 | 22.0 ± 5.0 | 30.7 ± 4.4 |
| 200 | 30 | 39.6 ± 6.9 | 39.6 ± 6.7 | 34.8 ± 5.9 | 20.8 ± 5.6 | 20.7 ± 5.1 | **19.6 ± 4.6** | 25.9 ± 7.4 | 22.6 ± 5.8 | 23.9 ± 5.2 | 31.6 ± 6.4 |
| *Number of false positive (FP) branches/clades* | | | | | | | | | | | |
| 50 | 10 | 6.8 ± 2.1 | 6.4 ± 2.2 | 5.9 ± 1.8 | 8.9 ± 1.7 | 6.7 ± 1.6 | 6.2 ± 2.5 | 5.5 ± 1.5 | 4.8 ± 1.8 | 2.1 ± 1.4 | **0.0 ± 0.0** |
| 50 | 20 | 7.0 ± 3.0 | 6.6 ± 2.6 | 8.2 ± 3.1 | 9.4 ± 2.6 | 7.4 ± 1.9 | 6.3 ± 2.2 | 5.1 ± 2.3 | 4.1 ± 2.0 | 2.2 ± 2.0 | **0.2 ± 0.6** |
| 50 | 30 | 9.2 ± 2.8 | 8.1 ± 3.0 | 9.5 ± 3.1 | 11.2 ± 2.9 | 7.8 ± 2.0 | 7.8 ± 2.3 | 6.5 ± 3.1 | 4.5 ± 1.5 | 2.9 ± 1.5 | **0.4 ± 0.6** |
| 100 | 10 | 15.0 ± 5.2 | 14.4 ± 5.5 | 13.4 ± 4.9 | 18.2 ± 3.9 | 14.1 ± 3.2 | 11.4 ± 3.6 | 10.8 ± 2.7 | 8.8 ± 2.6 | 4.4 ± 1.9 | **0.3 ± 0.5** |
| 100 | 20 | 17.9 ± 4.0 | 17.5 ± 4.0 | 17.5 ± 4.0 | 20.0 ± 3.9 | 14.8 ± 3.4 | 14.1 ± 4.1 | 14.3 ± 4.1 | 8.4 ± 2.5 | 4.4 ± 1.4 | **0.3 ± 0.6** |
| 100 | 30 | 19.5 ± 4.8 | 19.4 ± 4.8 | 22.5 ± 4.6 | 20.1 ± 2.8 | 15.1 ± 2.5 | 15.3 ± 3.4 | 13.3 ± 5.3 | 8.0 ± 2.2 | 4.2 ± 1.8 | **0.6 ± 0.9** |
| 150 | 10 | 24.7 ± 4.8 | 23.9 ± 4.6 | 22.1 ± 5.6 | 25.2 ± 4.8 | 20.4 ± 4.0 | 16.8 ± 4.6 | 16.8 ± 3.5 | 14.1 ± 4.0 | 7.5 ± 3.2 | **1.0 ± 1.4** |
| 150 | 20 | 31.0 ± 6.7 | 28.9 ± 6.1 | 30.5 ± 2.5 | 31.4 ± 4.1 | 22.3 ± 3.7 | 20.5 ± 3.7 | 21.5 ± 5.1 | 13.7 ± 3.3 | 6.5 ± 1.9 | **1.1 ± 1.0** |
| 150 | 30 | 30.8 ± 6.3 | 29.9 ± 4.5 | 33.7 ± 5.9 | 30.0 ± 4.9 | 22.1 ± 4.3 | 21.6 ± 5.0 | 19.3 ± 3.9 | 13.7 ± 4.8 | 7.8 ± 2.9 | **1.4 ± 1.5** |
| 200 | 10 | 35.6 ± 6.6 | 34.5 ± 6.5 | 28.9 ± 7.9 | 35.2 ± 5.6 | 28.6 ± 4.5 | 23.0 ± 4.9 | 23.7 ± 5.9 | 18.1 ± 4.8 | 9.2 ± 2.9 | **1.4 ± 1.0** |
| 200 | 20 | 41.8 ± 9.1 | 41.0 ± 10.1 | 42.2 ± 4.5 | 41.8 ± 4.3 | 31.3 ± 3.8 | 27.4 ± 5.4 | 28.1 ± 7.2 | 19.3 ± 5.6 | 11.1 ± 4.2 | **2.6 ± 2.5** |
| 200 | 30 | 44.5 ± 10.1 | 43.4 ± 9.0 | 46.7 ± 8.1 | 41.9 ± 5.6 | 31.8 ± 4.5 | 30.2 ± 5.3 | 26.7 ± 7.1 | 18.9 ± 4.5 | 11.6 ± 2.7 | **3.4 ± 5.3** |
| *Number of true positive (TP) branches/clades* | | | | | | | | | | | |
| 50 | 10 | 3.9 ± 2.1 | 4.1 ± 2.4 | 4.7 ± 2.7 | 4.7 ± 2.7 | 4.7 ± 2.9 | 4.9 ± 2.7 | 5.0 ± 2.5 | **5.2 ± 2.8** | 4.7 ± 2.4 | 2.7 ± 2.2 |
| 50 | 20 | 8.4 ± 3.7 | 8.6 ± 3.8 | 9.2 ± 3.0 | **10.6 ± 4.7** | 10.4 ± 4.5 | 10.5 ± 4.3 | 10.2 ± 4.7 | 10.4 ± 4.0 | 9.9 ± 4.0 | 7.7 ± 3.9 |
| 50 | 30 | 11.1 ± 4.4 | 11.2 ± 4.2 | 12.1 ± 4.2 | 13.6 ± 4.2 | **13.8 ± 3.8** | 13.5 ± 4.1 | 12.6 ± 5.2 | 13.3 ± 4.2 | 12.4 ± 3.4 | 10.1 ± 3.6 |
| 100 | 10 | 5.0 ± 2.3 | 5.0 ± 2.3 | 7.2 ± 3.0 | 7.2 ± 3.2 | 7.0 ± 3.4 | **8.6 ± 3.0** | 7.0 ± 3.1 | 8.5 ± 3.0 | 7.6 ± 2.8 | 3.6 ± 2.5 |
| 100 | 20 | 13.8 ± 4.4 | 14.0 ± 4.5 | 16.3 ± 4.5 | 19.8 ± 6.3 | 19.8 ± 6.2 | 19.9 ± 6.4 | 16.4 ± 4.3 | **20.4 ± 5.6** | 18.2 ± 6.4 | 13.5 ± 5.8 |
| 100 | 30 | 20.0 ± 7.1 | 20.2 ± 7.2 | 19.7 ± 6.2 | **27.9 ± 6.6** | 27.5 ± 6.1 | 27.6 ± 6.0 | 24.6 ± 7.9 | 27.0 ± 5.2 | 25.8 ± 5.8 | 21.9 ± 5.8 |
| 150 | 10 | 5.5 ± 3.2 | 5.7 ± 3.3 | 8.0 ± 3.7 | 9.2 ± 4.6 | 9.5 ± 4.4 | **11.2 ± 4.6** | 8.4 ± 3.9 | 10.3 ± 4.5 | 10.2 ± 3.4 | 4.5 ± 3.0 |
| 150 | 20 | 14.0 ± 4.2 | 14.4 ± 4.1 | 16.9 ± 5.2 | 22.3 ± 5.7 | 23.9 ± 5.3 | **24.8 ± 4.7** | 19.6 ± 4.9 | 24.4 ± 4.4 | 22.2 ± 4.4 | 14.8 ± 5.1 |
| 150 | 30 | 25.4 ± 9.6 | 25.7 ± 9.9 | 28.1 ± 9.0 | 39.6 ± 11.2 | 39.6 ± 9.9 | **40.6 ± 10.1** | 35.9 ± 11.6 | 38.4 ± 8.8 | 37.1 ± 10.0 | 30.0 ± 10.8 |
| 200 | 10 | 7.4 ± 3.1 | 7.2 ± 3.0 | 10.7 ± 4.3 | 12.6 ± 5.1 | 12.8 ± 5.4 | 15.4 ± 5.3 | 11.8 ± 4.4 | **15.7 ± 3.7** | 14.4 ± 4.0 | 6.8 ± 3.3 |
| 200 | 20 | 15.9 ± 8.4 | 16.0 ± 8.4 | 20.3 ± 9.4 | 28.9 ± 12.1 | 29.4 ± 12.2 | **31.0 ± 11.0** | 25.2 ± 11.7 | 30.7 ± 9.0 | 28.7 ± 10.3 | 19.9 ± 9.7 |
| 200 | 30 | 29.2 ± 13.8 | 29.2 ± 13.8 | 34.0 ± 13.5 | 48.0 ± 15.9 | 48.1 ± 14.5 | **49.2 ± 14.2** | 43.0 ± 16.2 | 46.2 ± 14.4 | 44.9 ± 14.4 | 37.2 ± 13.5 |

Table S3: **Tree accuracy and runtime (in seconds) for Startle simulated data sets.** Mean ± standard deviations are across replicates for each method. Tree accuracy metrics were computed after SH-contraction of both true and estimated trees.

| # of cells | # of chars | Startle NNI (C++) | LAML | Cassiopeia Greedy | StarCDP Rand | StarCDP Bias | PAUP* | Startle NNI (Py) | Startle ILP | StarCDP SC | PAUP* SC |
|---|---|---|---|---|---|---|---|---|---|---|---|
| *Precision = TP / (TP + FP)* | | | | | | | | | | | |
| 50 | 10 | 0.35 ± 0.15 | 0.37 ± 0.16 | 0.42 ± 0.14 | 0.32 ± 0.14 | 0.37 ± 0.17 | 0.43 ± 0.16 | 0.45 ± 0.16 | 0.50 ± 0.19 | 0.70 ± 0.16 | **0.94 ± 0.23** |
| 50 | 20 | 0.53 ± 0.18 | 0.55 ± 0.18 | 0.53 ± 0.16 | 0.51 ± 0.18 | 0.55 ± 0.17 | 0.60 ± 0.20 | 0.63 ± 0.18 | 0.69 ± 0.20 | 0.79 ± 0.22 | **0.93 ± 0.22** |
| 50 | 30 | 0.53 ± 0.14 | 0.57 ± 0.16 | 0.55 ± 0.15 | 0.54 ± 0.14 | 0.63 ± 0.11 | 0.63 ± 0.11 | 0.64 ± 0.20 | 0.73 ± 0.12 | 0.81 ± 0.09 | **0.96 ± 0.05** |
| 100 | 10 | 0.25 ± 0.12 | 0.26 ± 0.13 | 0.36 ± 0.12 | 0.28 ± 0.11 | 0.32 ± 0.12 | 0.43 ± 0.08 | 0.38 ± 0.13 | 0.48 ± 0.12 | 0.63 ± 0.11 | **0.84 ± 0.29** |
| 100 | 20 | 0.43 ± 0.12 | 0.44 ± 0.11 | 0.48 ± 0.12 | 0.49 ± 0.12 | 0.56 ± 0.13 | 0.58 ± 0.13 | 0.53 ± 0.12 | 0.70 ± 0.10 | 0.79 ± 0.10 | **0.98 ± 0.05** |
| 100 | 30 | 0.50 ± 0.13 | 0.50 ± 0.13 | 0.46 ± 0.12 | 0.57 ± 0.10 | 0.64 ± 0.08 | 0.64 ± 0.08 | 0.64 ± 0.18 | 0.77 ± 0.06 | 0.86 ± 0.04 | **0.98 ± 0.04** |
| 150 | 10 | 0.18 ± 0.11 | 0.19 ± 0.11 | 0.27 ± 0.12 | 0.26 ± 0.10 | 0.31 ± 0.10 | 0.40 ± 0.12 | 0.32 ± 0.13 | 0.42 ± 0.15 | 0.58 ± 0.16 | **0.81 ± 0.24** |
| 150 | 20 | 0.31 ± 0.10 | 0.34 ± 0.09 | 0.35 ± 0.08 | 0.41 ± 0.08 | 0.51 ± 0.09 | 0.55 ± 0.06 | 0.48 ± 0.11 | 0.64 ± 0.09 | 0.77 ± 0.08 | **0.93 ± 0.06** |
| 150 | 30 | 0.44 ± 0.12 | 0.45 ± 0.12 | 0.45 ± 0.10 | 0.56 ± 0.10 | 0.64 ± 0.08 | 0.65 ± 0.10 | 0.63 ± 0.11 | 0.74 ± 0.09 | 0.82 ± 0.07 | **0.95 ± 0.05** |
| 200 | 10 | 0.17 ± 0.07 | 0.17 ± 0.07 | 0.28 ± 0.11 | 0.26 ± 0.08 | 0.30 ± 0.10 | 0.39 ± 0.09 | 0.33 ± 0.11 | 0.47 ± 0.08 | 0.60 ± 0.11 | **0.83 ± 0.13** |
| 200 | 20 | 0.27 ± 0.11 | 0.27 ± 0.11 | 0.31 ± 0.10 | 0.39 ± 0.10 | 0.47 ± 0.11 | 0.52 ± 0.07 | 0.46 ± 0.14 | 0.61 ± 0.08 | 0.71 ± 0.11 | **0.89 ± 0.11** |
| 200 | 30 | 0.38 ± 0.15 | 0.39 ± 0.14 | 0.41 ± 0.12 | 0.52 ± 0.11 | 0.59 ± 0.10 | 0.61 ± 0.10 | 0.60 ± 0.14 | 0.69 ± 0.11 | 0.78 ± 0.07 | **0.92 ± 0.09** |
| *Recall = TP / (TP + FN)* | | | | | | | | | | | |
| 50 | 10 | 0.39 ± 0.14 | 0.40 ± 0.15 | 0.47 ± 0.16 | 0.46 ± 0.18 | 0.44 ± 0.20 | 0.50 ± 0.20 | 0.50 ± 0.18 | **0.52 ± 0.20** | 0.49 ± 0.18 | 0.26 ± 0.15 |
| 50 | 20 | 0.54 ± 0.16 | 0.56 ± 0.16 | 0.60 ± 0.11 | **0.66 ± 0.20** | **0.66 ± 0.16** | **0.66 ± 0.19** | 0.64 ± 0.17 | **0.66 ± 0.18** | 0.62 ± 0.18 | 0.47 ± 0.18 |
| 50 | 30 | 0.60 ± 0.18 | 0.60 ± 0.17 | 0.65 ± 0.16 | 0.73 ± 0.17 | **0.75 ± 0.12** | 0.73 ± 0.12 | 0.67 ± 0.20 | 0.72 ± 0.13 | 0.68 ± 0.11 | 0.55 ± 0.14 |
| 100 | 10 | 0.28 ± 0.11 | 0.28 ± 0.11 | 0.42 ± 0.12 | 0.41 ± 0.15 | 0.40 ± 0.16 | **0.50 ± 0.12** | 0.40 ± 0.13 | 0.49 ± 0.12 | 0.44 ± 0.12 | 0.19 ± 0.12 |
| 100 | 20 | 0.46 ± 0.12 | 0.46 ± 0.12 | 0.54 ± 0.12 | 0.64 ± 0.13 | 0.64 ± 0.14 | 0.64 ± 0.14 | 0.54 ± 0.12 | **0.66 ± 0.11** | 0.59 ± 0.15 | 0.43 ± 0.15 |
| 100 | 30 | 0.52 ± 0.14 | 0.52 ± 0.14 | 0.51 ± 0.12 | **0.73 ± 0.11** | 0.72 ± 0.08 | **0.73 ± 0.08** | 0.64 ± 0.18 | 0.71 ± 0.06 | 0.68 ± 0.09 | 0.57 ± 0.10 |
| 150 | 10 | 0.22 ± 0.11 | 0.22 ± 0.11 | 0.32 ± 0.13 | 0.36 ± 0.14 | 0.37 ± 0.12 | **0.45 ± 0.15** | 0.34 ± 0.13 | 0.42 ± 0.14 | 0.42 ± 0.12 | 0.17 ± 0.10 |
| 150 | 20 | 0.36 ± 0.10 | 0.37 ± 0.10 | 0.42 ± 0.10 | 0.56 ± 0.10 | 0.60 ± 0.10 | **0.63 ± 0.07** | 0.50 ± 0.10 | 0.62 ± 0.08 | 0.56 ± 0.07 | 0.37 ± 0.11 |
| 150 | 30 | 0.45 ± 0.12 | 0.45 ± 0.12 | 0.50 ± 0.12 | 0.71 ± 0.11 | 0.71 ± 0.09 | **0.73 ± 0.09** | 0.64 ± 0.13 | 0.69 ± 0.08 | 0.66 ± 0.10 | 0.53 ± 0.13 |
| 200 | 10 | 0.23 ± 0.08 | 0.22 ± 0.08 | 0.33 ± 0.11 | 0.39 ± 0.11 | 0.39 ± 0.12 | 0.48 ± 0.10 | 0.37 ± 0.13 | **0.50 ± 0.10** | 0.45 ± 0.09 | 0.21 ± 0.09 |
| 200 | 20 | 0.30 ± 0.12 | 0.30 ± 0.12 | 0.39 ± 0.12 | 0.55 ± 0.11 | 0.56 ± 0.12 | **0.60 ± 0.10** | 0.48 ± 0.14 | **0.60 ± 0.08** | 0.56 ± 0.10 | 0.38 ± 0.11 |
| 200 | 30 | 0.41 ± 0.14 | 0.41 ± 0.14 | 0.48 ± 0.11 | 0.68 ± 0.12 | 0.69 ± 0.10 | **0.70 ± 0.09** | 0.61 ± 0.14 | 0.66 ± 0.11 | 0.64 ± 0.11 | 0.53 ± 0.12 |
| *f1-score* | | | | | | | | | | | |
| 50 | 10 | 0.37 ± 0.14 | 0.38 ± 0.16 | 0.44 ± 0.15 | 0.38 ± 0.15 | 0.40 ± 0.18 | 0.46 ± 0.17 | 0.47 ± 0.16 | 0.50 ± 0.19 | **0.56 ± 0.15** | 0.39 ± 0.18 |
| 50 | 20 | 0.53 ± 0.17 | 0.55 ± 0.17 | 0.56 ± 0.14 | 0.57 ± 0.19 | 0.60 ± 0.17 | 0.62 ± 0.19 | 0.64 ± 0.18 | 0.67 ± 0.19 | **0.69 ± 0.18** | 0.61 ± 0.20 |
| 50 | 30 | 0.56 ± 0.16 | 0.58 ± 0.16 | 0.60 ± 0.16 | 0.62 ± 0.15 | 0.68 ± 0.12 | 0.67 ± 0.11 | 0.65 ± 0.20 | 0.72 ± 0.12 | **0.73 ± 0.09** | 0.68 ± 0.14 |
| 100 | 10 | 0.26 ± 0.11 | 0.26 ± 0.11 | 0.38 ± 0.12 | 0.33 ± 0.12 | 0.35 ± 0.14 | 0.46 ± 0.10 | 0.39 ± 0.13 | 0.49 ± 0.12 | **0.51 ± 0.12** | 0.30 ± 0.17 |
| 100 | 20 | 0.44 ± 0.12 | 0.45 ± 0.12 | 0.51 ± 0.11 | 0.55 ± 0.13 | 0.60 ± 0.13 | 0.61 ± 0.13 | 0.54 ± 0.12 | **0.68 ± 0.10** | 0.67 ± 0.13 | 0.58 ± 0.16 |
| 100 | 30 | 0.50 ± 0.14 | 0.51 ± 0.14 | 0.48 ± 0.12 | 0.64 ± 0.10 | 0.68 ± 0.08 | 0.68 ± 0.08 | 0.64 ± 0.18 | 0.74 ± 0.06 | **0.75 ± 0.07** | 0.71 ± 0.09 |
| 150 | 10 | 0.20 ± 0.11 | 0.20 ± 0.11 | 0.29 ± 0.12 | 0.30 ± 0.12 | 0.34 ± 0.11 | 0.42 ± 0.13 | 0.33 ± 0.13 | 0.41 ± 0.15 | **0.48 ± 0.13** | 0.28 ± 0.13 |
| 150 | 20 | 0.33 ± 0.10 | 0.35 ± 0.10 | 0.38 ± 0.09 | 0.47 ± 0.08 | 0.55 ± 0.09 | 0.58 ± 0.06 | 0.48 ± 0.11 | 0.63 ± 0.09 | **0.65 ± 0.07** | 0.52 ± 0.12 |
| 150 | 30 | 0.44 ± 0.12 | 0.45 ± 0.12 | 0.47 ± 0.11 | 0.62 ± 0.10 | 0.67 ± 0.08 | 0.68 ± 0.09 | 0.64 ± 0.12 | 0.71 ± 0.08 | **0.73 ± 0.08** | 0.67 ± 0.12 |
| 200 | 10 | 0.19 ± 0.07 | 0.19 ± 0.07 | 0.30 ± 0.11 | 0.31 ± 0.09 | 0.34 ± 0.11 | 0.43 ± 0.09 | 0.35 ± 0.12 | 0.48 ± 0.09 | **0.51 ± 0.09** | 0.32 ± 0.12 |
| 200 | 20 | 0.28 ± 0.11 | 0.29 ± 0.11 | 0.34 ± 0.10 | 0.46 ± 0.11 | 0.51 ± 0.12 | 0.56 ± 0.08 | 0.47 ± 0.14 | 0.61 ± 0.08 | **0.62 ± 0.10** | 0.52 ± 0.11 |
| 200 | 30 | 0.40 ± 0.14 | 0.40 ± 0.14 | 0.44 ± 0.12 | 0.59 ± 0.11 | 0.63 ± 0.10 | 0.65 ± 0.09 | 0.60 ± 0.14 | 0.68 ± 0.11 | **0.70 ± 0.09** | 0.66 ± 0.11 |
| *Runtime (in seconds)* | | | | | | | | | | | |
| 50 | 10 | 3.5 ± 0.2 | 443.3 ± 337.7 | 5.8 ± 0.8 | 3.7 ± 0.3 | 3.8 ± 0.3 | **1.5 ± 0.2** | 70.1 ± 22.2 | 13.6 ± 2.8 | NA | NA |
| 50 | 20 | 3.6 ± 0.3 | 435.5 ± 286.1 | 5.5 ± 0.3 | 3.8 ± 0.3 | 3.8 ± 0.3 | **1.6 ± 0.2** | 184.1 ± 79.4 | 14.3 ± 2.4 | NA | NA |
| 50 | 30 | 3.7 ± 0.4 | 271.6 ± 127.4 | 5.7 ± 0.5 | 4.0 ± 0.3 | 4.0 ± 0.3 | **1.6 ± 0.2** | 260.3 ± 76.6 | 15.2 ± 2.4 | NA | NA |
| 100 | 10 | 5.2 ± 0.5 | 4228.6 ± 4032.4 | 5.6 ± 0.4 | 8.4 ± 1.2 | 8.5 ± 1.2 | **3.5 ± 0.5** | 149.8 ± 80.9 | 13.9 ± 1.3 | NA | NA |
| 100 | 20 | 5.5 ± 0.6 | 2897.2 ± 2696.1 | 6.4 ± 1.2 | 9.2 ± 0.9 | 9.2 ± 0.9 | **4.0 ± 0.7** | 434.9 ± 124.2 | 16.6 ± 4.2 | NA | NA |
| 100 | 30 | 5.8 ± 1.1 | 2165.4 ± 1385.1 | 5.7 ± 0.6 | 10.0 ± 0.9 | 10.0 ± 0.9 | **4.3 ± 0.6** | 843.1 ± 226.6 | 23.1 ± 10.2 | NA | NA |
| 150 | 10 | 7.6 ± 0.7 | 10578.2 ± 4238.3 | 6.2 ± 0.6 | 13.0 ± 1.6 | 13.3 ± 2.3 | **5.6 ± 1.0** | 210.3 ± 92.6 | 15.2 ± 3.4 | NA | NA |
| 150 | 20 | 9.2 ± 1.4 | 12551.5 ± 9824.4 | 6.8 ± 0.5 | 14.8 ± 1.6 | 14.7 ± 1.4 | **6.4 ± 0.9** | 1030.3 ± 396.8 | 24.9 ± 9.2 | NA | NA |
| 150 | 30 | 10.1 ± 1.5 | 9358.1 ± 5405.3 | **5.9 ± 1.1** | 18.3 ± 2.4 | 18.4 ± 2.2 | 8.6 ± 1.7 | 2191.0 ± 600.6 | 99.3 ± 148.8 | NA | NA |
| 200 | 10 | 11.4 ± 1.1 | 19640.8 ± 13419.4 | **6.7 ± 1.2** | 22.3 ± 5.9 | 22.0 ± 5.5 | 9.5 ± 1.7 | 358.0 ± 116.7 | 22.3 ± 29.4 | NA | NA |
| 200 | 20 | 13.1 ± 2.0 | 27874.5 ± 17153.7 | **7.1 ± 1.3** | 23.2 ± 4.0 | 23.3 ± 4.2 | 10.1 ± 2.5 | 1727.1 ± 1150.1 | 434.8 ± 1200.0 | NA | NA |
| 200 | 30 | 15.3 ± 2.7 | 18745.0 ± 10537.1 | **11.8 ± 4.7** | 28.4 ± 3.6 | 28.3 ± 3.5 | 12.8 ± 2.2 | 4705.4 ± 1711.9 | 172.0 ± 114.8 | NA | NA |

#### 3.2 Supplemental results on LAML simulated data sets

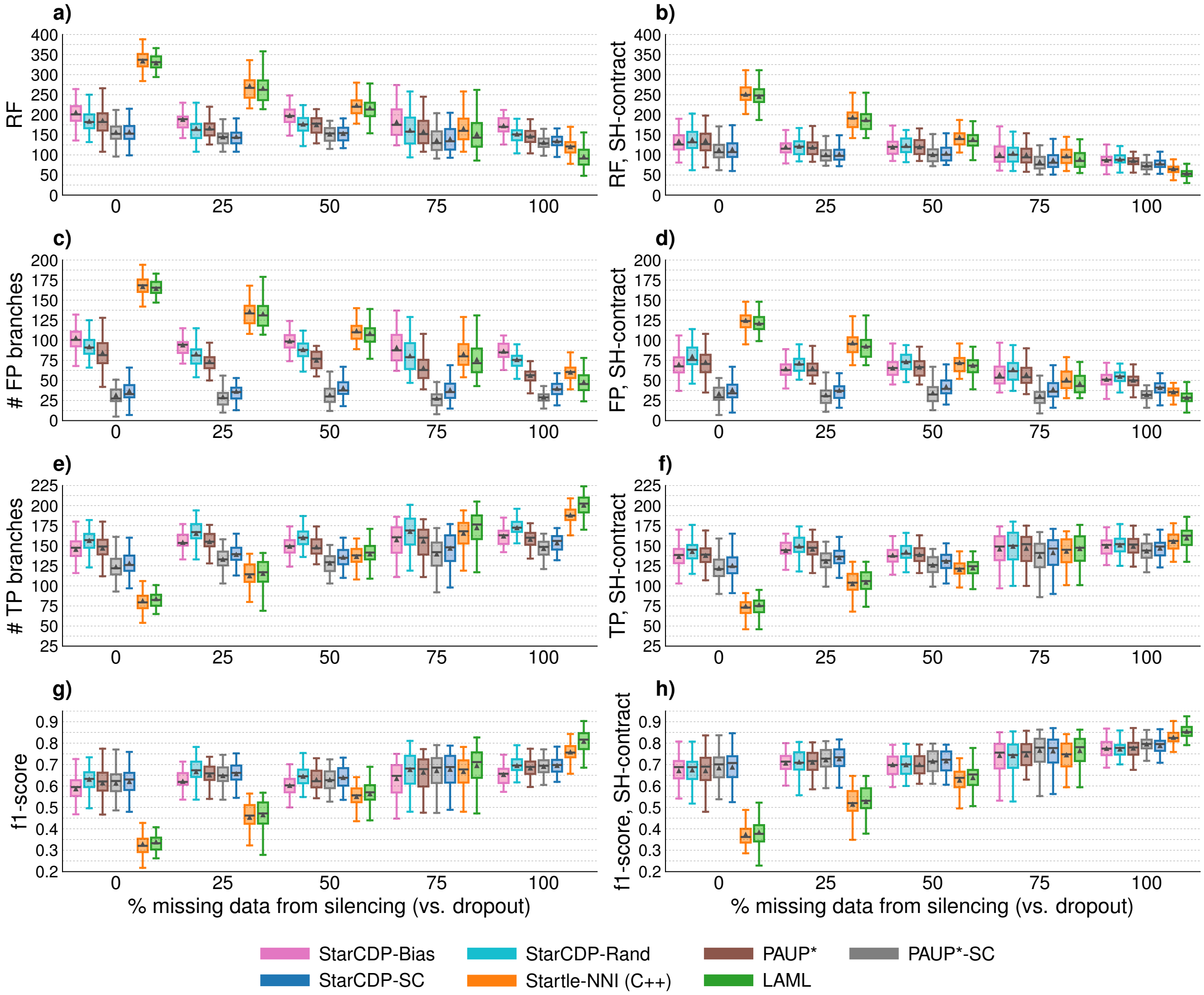

Figure S11: **Tree error and accuracy for LAML simulated data sets with 250 cells.** Subplots **(a)**–**(d)** show error metrics (lower is better). Subplots **(e)**–**(h)** show accuracy metrics (higher is better). For subplots in the right column, SH contraction is performed on both true and estimated trees before computing error and accuracy metrics, unlike subplots in the left column (where the true tree is binary and the estimated trees are binary except PAUP*, PAUP*-SC, and Star-CDP-SC). Triangles and bars are means and medians across replicate data sets, respectively

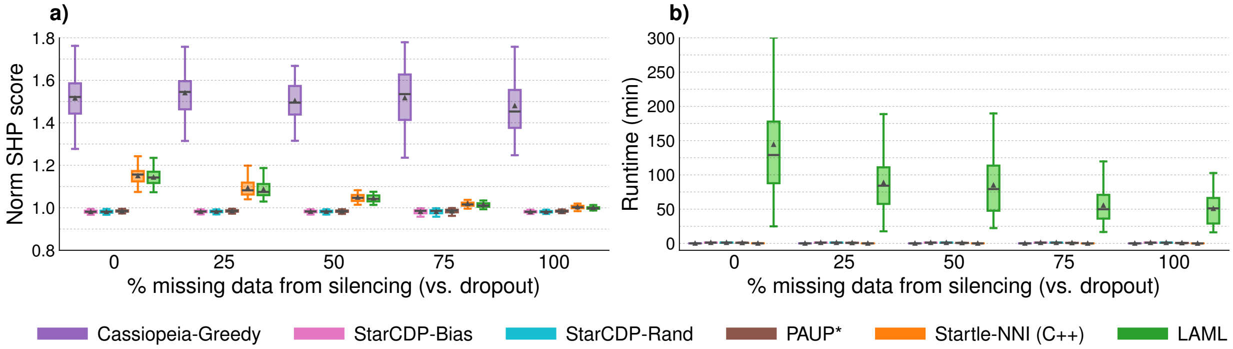

Figure S12: **Normalized SH parsimony score and runtime (in minutes) for LAML simulated data sets with 250 cells.** Scores are normalized by dividing the score of the score of the true tree. Runtime for Star-CDP includes the time to compute consensus trees (e.g., Star-CDP-SC). Runtime for PAUP*-SC is not shown because it was negligible compared to the time to run the heuristic search.

Table S4: **Tree error (no contraction) for LAML simulated data sets with 250 cells.** Mean ± standard deviations are across 50 replicates for each method.

| Proportion silencing | Cassiopeia Greedy | StarCDP Bias | StarCDP Rand | PAUP* | PAUP* SC | StarCDP SC | Startle NNI (C++) | LAML |
|---|---|---|---|---|---|---|---|---|
| *RF distance = # FN + # FP* | | | | | | | | |
| 0 | $406.0 \pm 21.2$ | $205.5 \pm 30.6$ | $183.4 \pm 37.4$ | $185.3 \pm 34.6$ | $\mathbf{156.1 \pm 27.7}$ | $156.9 \pm 30.0$ | $333.5 \pm 25.6$ | $328.4 \pm 28.5$ |
| 25 | $394.3 \pm 19.8$ | $188.0 \pm 26.3$ | $165.7 \pm 29.7$ | $166.5 \pm 26.3$ | $\mathbf{144.5 \pm 21.9}$ | $145.3 \pm 23.4$ | $271.4 \pm 33.4$ | $265.4 \pm 33.2$ |
| 50 | $375.5 \pm 20.5$ | $197.7 \pm 21.1$ | $176.4 \pm 23.0$ | $174.8 \pm 20.8$ | $\mathbf{151.5 \pm 18.7}$ | $153.0 \pm 18.8$ | $223.3 \pm 28.0$ | $216.9 \pm 28.8$ |
| 75 | $368.4 \pm 22.8$ | $181.6 \pm 34.4$ | $160.8 \pm 37.4$ | $157.6 \pm 32.1$ | $\mathbf{136.1 \pm 27.2}$ | $139.4 \pm 28.7$ | $165.4 \pm 33.8$ | $151.6 \pm 39.2$ |
| 100 | $341.5 \pm 24.1$ | $172.3 \pm 22.7$ | $151.6 \pm 21.1$ | $146.3 \pm 18.9$ | $130.8 \pm 15.2$ | $134.4 \pm 16.0$ | $120.1 \pm 21.4$ | $\mathbf{95.4 \pm 27.4}$ |
| *Number of false negative (FN) branches* | | | | | | | | |
| 0 | $203.3 \pm 10.7$ | $102.8 \pm 15.3$ | $\mathbf{91.7 \pm 18.7}$ | $101.2 \pm 16.6$ | $124.5 \pm 15.9$ | $120.1 \pm 16.1$ | $166.7 \pm 12.8$ | $164.2 \pm 14.3$ |
| 25 | $197.4 \pm 9.9$ | $94.0 \pm 13.2$ | $\mathbf{82.9 \pm 14.9}$ | $93.1 \pm 13.4$ | $115.1 \pm 13.3$ | $109.1 \pm 13.3$ | $135.7 \pm 16.7$ | $132.7 \pm 16.6$ |
| 50 | $188.4 \pm 10.2$ | $98.9 \pm 10.5$ | $\mathbf{88.2 \pm 11.5}$ | $99.8 \pm 10.6$ | $119.8 \pm 11.1$ | $112.6 \pm 11.2$ | $111.6 \pm 14.0$ | $108.5 \pm 14.4$ |
| 75 | $184.7 \pm 11.6$ | $90.8 \pm 17.2$ | $80.4 \pm 18.7$ | $92.4 \pm 17.3$ | $108.5 \pm 17.6$ | $101.4 \pm 18.0$ | $82.7 \pm 16.9$ | $\mathbf{75.8 \pm 19.6}$ |
| 100 | $171.1 \pm 12.0$ | $86.2 \pm 11.4$ | $75.8 \pm 10.5$ | $90.2 \pm 10.6$ | $101.7 \pm 11.8$ | $94.7 \pm 11.0$ | $60.1 \pm 10.7$ | $\mathbf{47.7 \pm 13.7}$ |
| *Number of false positive (FP) branches* | | | | | | | | |
| 0 | $202.7 \pm 10.6$ | $102.8 \pm 15.3$ | $91.7 \pm 18.7$ | $84.1 \pm 18.3$ | $\mathbf{31.5 \pm 14.7}$ | $36.8 \pm 16.2$ | $166.7 \pm 12.8$ | $164.2 \pm 14.3$ |
| 25 | $196.9 \pm 9.8$ | $94.0 \pm 13.2$ | $82.9 \pm 14.9$ | $73.4 \pm 13.5$ | $\mathbf{29.3 \pm 11.0}$ | $36.3 \pm 12.5$ | $135.7 \pm 16.7$ | $132.7 \pm 16.6$ |
| 50 | $187.1 \pm 10.3$ | $98.9 \pm 10.5$ | $88.2 \pm 11.5$ | $75.0 \pm 11.0$ | $\mathbf{31.7 \pm 11.1}$ | $40.4 \pm 11.2$ | $111.6 \pm 14.0$ | $108.5 \pm 14.4$ |
| 75 | $183.6 \pm 11.2$ | $90.8 \pm 17.2$ | $80.4 \pm 18.7$ | $65.3 \pm 15.6$ | $\mathbf{27.6 \pm 12.1}$ | $38.1 \pm 13.3$ | $82.7 \pm 16.9$ | $75.8 \pm 19.6$ |
| 100 | $170.5 \pm 12.1$ | $86.2 \pm 11.4$ | $75.8 \pm 10.5$ | $56.1 \pm 9.6$ | $\mathbf{29.2 \pm 8.0}$ | $39.8 \pm 9.5$ | $60.1 \pm 10.7$ | $47.7 \pm 13.7$ |
| *Number of true positive (TP) branches* | | | | | | | | |
| 0 | $44.7 \pm 10.7$ | $145.2 \pm 15.3$ | $\mathbf{156.3 \pm 18.7}$ | $146.8 \pm 16.6$ | $123.5 \pm 15.9$ | $127.9 \pm 16.1$ | $81.3 \pm 12.8$ | $83.8 \pm 14.3$ |
| 25 | $50.6 \pm 9.9$ | $154.0 \pm 13.2$ | $\mathbf{165.1 \pm 14.9}$ | $154.9 \pm 13.4$ | $132.9 \pm 13.3$ | $138.9 \pm 13.3$ | $112.3 \pm 16.7$ | $115.3 \pm 16.6$ |
| 50 | $59.6 \pm 10.2$ | $149.1 \pm 10.5$ | $\mathbf{159.8 \pm 11.5}$ | $148.2 \pm 10.6$ | $128.2 \pm 11.1$ | $135.4 \pm 11.2$ | $136.4 \pm 14.0$ | $139.5 \pm 14.4$ |
| 75 | $63.3 \pm 11.6$ | $157.2 \pm 17.2$ | $167.6 \pm 18.7$ | $155.6 \pm 17.3$ | $139.5 \pm 17.6$ | $146.6 \pm 18.0$ | $165.3 \pm 16.9$ | $\mathbf{172.2 \pm 19.6}$ |
| 100 | $76.9 \pm 12.0$ | $161.8 \pm 11.4$ | $172.2 \pm 10.5$ | $157.8 \pm 10.6$ | $146.3 \pm 11.8$ | $153.3 \pm 11.0$ | $187.9 \pm 10.7$ | $\mathbf{200.3 \pm 13.7}$ |

Table S5: **Tree error (SH contraction) for LAML simulated data sets with 250 cells.** Mean ± standard deviations are across 50 replicates for each method.

| Proportion silencing | Cassiopeia Greedy | StarCDP Bias | StarCDP Rand | PAUP* | PAUP* SC | StarCDP SC | Startle NNI (C++) | LAML |
|---|---|---|---|---|---|---|---|---|
| *RF distance = # FN + # FP* | | | | | | | | |
| 0 | $340.1 \pm 18.6$ | $134.0 \pm 28.8$ | $137.8 \pm 32.3$ | $135.4 \pm 31.3$ | $\mathbf{112.3 \pm 26.4}$ | $113.7 \pm 28.2$ | $250.6 \pm 22.6$ | $245.2 \pm 24.8$ |
| 25 | $329.7 \pm 19.4$ | $119.7 \pm 21.5$ | $121.1 \pm 23.1$ | $118.8 \pm 23.5$ | $\mathbf{100.5 \pm 19.6}$ | $101.7 \pm 20.8$ | $193.4 \pm 30.2$ | $187.4 \pm 28.9$ |
| 50 | $303.3 \pm 22.0$ | $119.1 \pm 20.2$ | $122.9 \pm 19.6$ | $119.6 \pm 20.3$ | $\mathbf{100.5 \pm 19.0}$ | $103.5 \pm 19.5$ | $143.6 \pm 23.4$ | $138.4 \pm 22.9$ |
| 75 | $293.5 \pm 23.4$ | $100.6 \pm 24.5$ | $103.5 \pm 26.2$ | $99.8 \pm 24.1$ | $\mathbf{82.8 \pm 21.8}$ | $86.7 \pm 22.7$ | $97.1 \pm 21.6$ | $88.9 \pm 21.9$ |
| 100 | $264.0 \pm 21.8$ | $86.7 \pm 15.9$ | $89.0 \pm 14.5$ | $85.9 \pm 15.3$ | $73.8 \pm 12.8$ | $79.1 \pm 13.8$ | $65.2 \pm 13.5$ | $\mathbf{54.0 \pm 13.3}$ |
| *Number of false negative (FN) branches* | | | | | | | | |
| 0 | $162.1 \pm 8.9$ | $64.2 \pm 13.8$ | $\mathbf{58.0 \pm 15.5}$ | $62.9 \pm 14.7$ | $78.9 \pm 14.2$ | $75.6 \pm 14.3$ | $126.1 \pm 10.9$ | $124.3 \pm 11.7$ |
| 25 | $156.3 \pm 9.5$ | $55.4 \pm 10.4$ | $\mathbf{49.9 \pm 11.1}$ | $53.7 \pm 11.2$ | $68.8 \pm 10.8$ | $64.2 \pm 10.8$ | $96.9 \pm 15.2$ | $95.0 \pm 14.6$ |
| 50 | $142.4 \pm 11.4$ | $53.9 \pm 9.4$ | $\mathbf{49.9 \pm 8.6}$ | $53.2 \pm 9.5$ | $66.2 \pm 9.4$ | $61.4 \pm 9.8$ | $71.6 \pm 11.5$ | $69.8 \pm 11.3$ |
| 75 | $136.9 \pm 10.9$ | $43.5 \pm 10.2$ | $\mathbf{40.1 \pm 11.1}$ | $42.7 \pm 9.8$ | $52.4 \pm 10.8$ | $47.9 \pm 10.6$ | $46.3 \pm 10.1$ | $43.0 \pm 10.9$ |
| 100 | $123.2 \pm 10.8$ | $35.4 \pm 7.0$ | $34.1 \pm 6.3$ | $35.7 \pm 6.5$ | $41.5 \pm 6.7$ | $38.0 \pm 6.5$ | $29.6 \pm 6.4$ | $\mathbf{25.4 \pm 6.4}$ |
| *Number of false positive (FP) branches* | | | | | | | | |
| 0 | $178.0 \pm 10.9$ | $69.8 \pm 15.9$ | $79.8 \pm 17.3$ | $72.5 \pm 17.4$ | $\mathbf{33.3 \pm 14.9}$ | $38.1 \pm 16.0$ | $124.6 \pm 12.7$ | $120.9 \pm 14.0$ |
| 25 | $173.4 \pm 10.6$ | $64.3 \pm 11.9$ | $71.2 \pm 12.5$ | $65.1 \pm 13.0$ | $\mathbf{31.7 \pm 11.3}$ | $37.5 \pm 12.4$ | $96.5 \pm 15.5$ | $92.4 \pm 14.8$ |
| 50 | $160.9 \pm 11.4$ | $65.2 \pm 11.6$ | $73.0 \pm 11.5$ | $66.4 \pm 11.7$ | $\mathbf{34.2 \pm 11.8}$ | $42.1 \pm 11.8$ | $71.9 \pm 12.7$ | $68.6 \pm 12.6$ |
| 75 | $156.6 \pm 13.6$ | $57.1 \pm 15.0$ | $63.4 \pm 15.5$ | $57.1 \pm 14.9$ | $\mathbf{30.3 \pm 13.0}$ | $38.8 \pm 13.9$ | $50.8 \pm 12.4$ | $45.9 \pm 11.8$ |
| 100 | $140.8 \pm 12.4$ | $51.3 \pm 9.9$ | $54.9 \pm 9.2$ | $50.2 \pm 10.0$ | $32.3 \pm 8.4$ | $41.0 \pm 9.3$ | $35.6 \pm 8.0$ | $\mathbf{28.6 \pm 7.6}$ |
| *Number of true positive (TP) branches* | | | | | | | | |
| 0 | $38.5 \pm 10.4$ | $136.4 \pm 15.4$ | $\mathbf{142.6 \pm 17.3}$ | $137.7 \pm 16.4$ | $121.7 \pm 15.7$ | $125.1 \pm 15.8$ | $74.6 \pm 12.4$ | $76.3 \pm 13.3$ |
| 25 | $43.0 \pm 9.6$ | $143.9 \pm 12.8$ | $\mathbf{149.3 \pm 13.7}$ | $145.5 \pm 13.4$ | $130.5 \pm 13.1$ | $135.1 \pm 13.3$ | $102.4 \pm 16.1$ | $104.3 \pm 15.6$ |
| 50 | $49.5 \pm 9.0$ | $138.0 \pm 10.7$ | $\mathbf{142.0 \pm 10.0}$ | $138.7 \pm 10.6$ | $125.7 \pm 10.8$ | $130.5 \pm 10.9$ | $120.3 \pm 12.8$ | $122.1 \pm 12.7$ |
| 75 | $52.3 \pm 11.1$ | $145.7 \pm 18.0$ | $\mathbf{149.1 \pm 19.1}$ | $146.5 \pm 17.7$ | $136.8 \pm 18.4$ | $141.2 \pm 18.5$ | $142.9 \pm 17.5$ | $146.2 \pm 18.5$ |
| 100 | $61.5 \pm 10.5$ | $149.3 \pm 11.6$ | $150.6 \pm 11.6$ | $149.1 \pm 11.5$ | $143.2 \pm 11.8$ | $146.7 \pm 11.5$ | $155.2 \pm 11.1$ | $\mathbf{159.3 \pm 12.8}$ |

Table S6: **Tree accuracy (no contraction) and runtime (in seconds) for LAML simulated data sets with 250 cells.** Mean ± standard deviations are across replicates for each method.

| Proportion silencing | Cassiopeia Greedy | StarCDP Bias | StarCDP Rand | PAUP* | PAUP* SC | StarCDP SC | Startle NNI (C++) | LAML |
|---|---|---|---|---|---|---|---|---|
| *Precision = TP / (TP + FP)* | | | | | | | | |
| 0 | $0.18 \pm 0.04$ | $0.59 \pm 0.06$ | $0.63 \pm 0.08$ | $0.64 \pm 0.08$ | $\mathbf{0.80 \pm 0.09}$ | $0.78 \pm 0.09$ | $0.33 \pm 0.05$ | $0.34 \pm 0.06$ |
| 25 | $0.20 \pm 0.04$ | $0.62 \pm 0.05$ | $0.67 \pm 0.06$ | $0.68 \pm 0.06$ | $\mathbf{0.82 \pm 0.07}$ | $0.79 \pm 0.07$ | $0.45 \pm 0.07$ | $0.46 \pm 0.07$ |
| 50 | $0.24 \pm 0.04$ | $0.60 \pm 0.04$ | $0.64 \pm 0.05$ | $0.66 \pm 0.05$ | $\mathbf{0.80 \pm 0.06}$ | $0.77 \pm 0.06$ | $0.55 \pm 0.06$ | $0.56 \pm 0.06$ |
| 75 | $0.26 \pm 0.05$ | $0.63 \pm 0.07$ | $0.68 \pm 0.08$ | $0.70 \pm 0.07$ | $\mathbf{0.83 \pm 0.07}$ | $0.79 \pm 0.07$ | $0.67 \pm 0.07$ | $0.69 \pm 0.08$ |
| 100 | $0.31 \pm 0.05$ | $0.65 \pm 0.05$ | $0.69 \pm 0.04$ | $0.74 \pm 0.04$ | $\mathbf{0.83 \pm 0.04}$ | $0.80 \pm 0.04$ | $0.76 \pm 0.04$ | $0.81 \pm 0.06$ |
| *Recall = TP / (TP + FN)* | | | | | | | | |
| 0 | $0.18 \pm 0.04$ | $0.59 \pm 0.06$ | $\mathbf{0.63 \pm 0.08}$ | $0.59 \pm 0.07$ | $0.50 \pm 0.06$ | $0.52 \pm 0.06$ | $0.33 \pm 0.05$ | $0.34 \pm 0.06$ |
| 25 | $0.20 \pm 0.04$ | $0.62 \pm 0.05$ | $\mathbf{0.67 \pm 0.06}$ | $0.62 \pm 0.05$ | $0.54 \pm 0.05$ | $0.56 \pm 0.05$ | $0.45 \pm 0.07$ | $0.46 \pm 0.07$ |
| 50 | $0.24 \pm 0.04$ | $0.60 \pm 0.04$ | $\mathbf{0.64 \pm 0.05}$ | $0.60 \pm 0.04$ | $0.52 \pm 0.04$ | $0.55 \pm 0.04$ | $0.55 \pm 0.06$ | $0.56 \pm 0.06$ |
| 75 | $0.26 \pm 0.05$ | $0.63 \pm 0.07$ | $0.68 \pm 0.08$ | $0.63 \pm 0.07$ | $0.56 \pm 0.07$ | $0.59 \pm 0.07$ | $0.67 \pm 0.07$ | $\mathbf{0.69 \pm 0.08}$ |
| 100 | $0.31 \pm 0.05$ | $0.65 \pm 0.05$ | $0.69 \pm 0.04$ | $0.64 \pm 0.04$ | $0.59 \pm 0.05$ | $0.62 \pm 0.04$ | $0.76 \pm 0.04$ | $\mathbf{0.81 \pm 0.06}$ |
| *f1-score* | | | | | | | | |
| 0 | $0.18 \pm 0.04$ | $0.59 \pm 0.06$ | $\mathbf{0.63 \pm 0.08}$ | $0.61 \pm 0.07$ | $0.61 \pm 0.07$ | $0.62 \pm 0.07$ | $0.33 \pm 0.05$ | $0.34 \pm 0.06$ |
| 25 | $0.20 \pm 0.04$ | $0.62 \pm 0.05$ | $\mathbf{0.67 \pm 0.06}$ | $0.65 \pm 0.06$ | $0.65 \pm 0.06$ | $0.66 \pm 0.06$ | $0.45 \pm 0.07$ | $0.46 \pm 0.07$ |
| 50 | $0.24 \pm 0.04$ | $0.60 \pm 0.04$ | $\mathbf{0.64 \pm 0.05}$ | $0.63 \pm 0.04$ | $0.63 \pm 0.05$ | $\mathbf{0.64 \pm 0.05}$ | $0.55 \pm 0.06$ | $0.56 \pm 0.06$ |
| 75 | $0.26 \pm 0.05$ | $0.63 \pm 0.07$ | $0.68 \pm 0.08$ | $0.66 \pm 0.07$ | $0.67 \pm 0.07$ | $0.68 \pm 0.07$ | $0.67 \pm 0.07$ | $\mathbf{0.69 \pm 0.08}$ |
| 100 | $0.31 \pm 0.05$ | $0.65 \pm 0.05$ | $0.69 \pm 0.04$ | $0.68 \pm 0.04$ | $0.69 \pm 0.04$ | $0.70 \pm 0.04$ | $0.76 \pm 0.04$ | $\mathbf{0.81 \pm 0.06}$ |
| *Runtime (in seconds)* | | | | | | | | |
| 0 | $\mathbf{7.3 \pm 3.0}$ | $70.5 \pm 8.2$ | $70.8 \pm 8.3$ | $55.8 \pm 8.1$ | NA | NA | $8.8 \pm 4.3$ | $8653.5 \pm 4592.1$ |
| 25 | $8.1 \pm 4.9$ | $68.0 \pm 8.2$ | $68.0 \pm 8.2$ | $51.8 \pm 7.8$ | NA | NA | $\mathbf{6.6 \pm 3.3}$ | $5312.6 \pm 2485.7$ |
| 50 | $7.5 \pm 4.1$ | $67.7 \pm 8.2$ | $67.3 \pm 7.7$ | $48.8 \pm 8.2$ | NA | NA | $\mathbf{5.9 \pm 2.5}$ | $5097.6 \pm 2743.6$ |
| 75 | $7.2 \pm 2.9$ | $66.4 \pm 7.1$ | $66.6 \pm 7.2$ | $47.2 \pm 6.7$ | NA | NA | $\mathbf{4.1 \pm 1.5}$ | $3326.6 \pm 1549.0$ |
| 100 | $6.2 \pm 0.8$ | $64.6 \pm 5.6$ | $64.6 \pm 5.3$ | $41.4 \pm 5.4$ | NA | NA | $\mathbf{3.5 \pm 0.9}$ | $3045.6 \pm 1588.5$ |

Table S7: **Tree accuracy (SH contraction) and runtime (in seconds) for LAML simulated data sets with 250 cells.** Mean ± standard deviations are across replicates for each method.

| Proportion silencing | Cassiopeia Greedy | StarCDP Bias | StarCDP Rand | PAUP* | PAUP* SC | StarCDP SC | Startle NNI (C++) | LAML |
|---|---|---|---|---|---|---|---|---|
| *Precision = TP / (TP + FP)* | | | | | | | | |
| 0 | $0.18 \pm 0.05$ | $0.66 \pm 0.07$ | $0.64 \pm 0.08$ | $0.66 \pm 0.08$ | $\mathbf{0.79 \pm 0.09}$ | $0.77 \pm 0.09$ | $0.37 \pm 0.06$ | $0.39 \pm 0.06$ |
| 25 | $0.20 \pm 0.04$ | $0.69 \pm 0.06$ | $0.68 \pm 0.06$ | $0.69 \pm 0.06$ | $\mathbf{0.80 \pm 0.07}$ | $0.78 \pm 0.07$ | $0.51 \pm 0.08$ | $0.53 \pm 0.08$ |
| 50 | $0.24 \pm 0.04$ | $0.68 \pm 0.05$ | $0.66 \pm 0.05$ | $0.68 \pm 0.05$ | $\mathbf{0.79 \pm 0.06}$ | $0.76 \pm 0.06$ | $0.63 \pm 0.06$ | $0.64 \pm 0.06$ |
| 75 | $0.25 \pm 0.05$ | $0.72 \pm 0.08$ | $0.70 \pm 0.08$ | $0.72 \pm 0.08$ | $\mathbf{0.82 \pm 0.08}$ | $0.78 \pm 0.08$ | $0.74 \pm 0.07$ | $0.76 \pm 0.07$ |
| 100 | $0.30 \pm 0.05$ | $0.74 \pm 0.05$ | $0.73 \pm 0.04$ | $0.75 \pm 0.05$ | $0.82 \pm 0.04$ | $0.78 \pm 0.04$ | $0.81 \pm 0.04$ | $\mathbf{0.85 \pm 0.04}$ |
| *Recall = TP / (TP + FN)* | | | | | | | | |
| 0 | $0.19 \pm 0.05$ | $0.68 \pm 0.07$ | $\mathbf{0.71 \pm 0.08}$ | $0.69 \pm 0.07$ | $0.61 \pm 0.07$ | $0.62 \pm 0.07$ | $0.37 \pm 0.06$ | $0.38 \pm 0.06$ |
| 25 | $0.22 \pm 0.05$ | $0.72 \pm 0.05$ | $\mathbf{0.75 \pm 0.06}$ | $0.73 \pm 0.06$ | $0.65 \pm 0.06$ | $0.68 \pm 0.06$ | $0.51 \pm 0.08$ | $0.52 \pm 0.08$ |
| 50 | $0.26 \pm 0.05$ | $0.72 \pm 0.05$ | $\mathbf{0.74 \pm 0.04}$ | $0.72 \pm 0.05$ | $0.66 \pm 0.05$ | $0.68 \pm 0.05$ | $0.63 \pm 0.06$ | $0.64 \pm 0.06$ |
| 75 | $0.28 \pm 0.05$ | $0.77 \pm 0.06$ | $\mathbf{0.78 \pm 0.07}$ | $0.77 \pm 0.06$ | $0.72 \pm 0.07$ | $0.74 \pm 0.07$ | $0.75 \pm 0.06$ | $0.77 \pm 0.07$ |
| 100 | $0.33 \pm 0.05$ | $0.81 \pm 0.04$ | $0.82 \pm 0.03$ | $0.81 \pm 0.04$ | $0.78 \pm 0.04$ | $0.79 \pm 0.04$ | $0.84 \pm 0.03$ | $\mathbf{0.86 \pm 0.04}$ |
| *f1-score* | | | | | | | | |
| 0 | $0.18 \pm 0.05$ | $0.67 \pm 0.07$ | $0.67 \pm 0.08$ | $0.67 \pm 0.08$ | $0.68 \pm 0.07$ | $\mathbf{0.69 \pm 0.08}$ | $0.37 \pm 0.06$ | $0.38 \pm 0.06$ |
| 25 | $0.21 \pm 0.04$ | $0.71 \pm 0.06$ | $0.71 \pm 0.06$ | $0.71 \pm 0.06$ | $0.72 \pm 0.06$ | $\mathbf{0.73 \pm 0.06}$ | $0.51 \pm 0.08$ | $0.53 \pm 0.08$ |
| 50 | $0.25 \pm 0.05$ | $0.70 \pm 0.05$ | $0.70 \pm 0.05$ | $0.70 \pm 0.05$ | $0.71 \pm 0.05$ | $\mathbf{0.72 \pm 0.05}$ | $0.63 \pm 0.06$ | $0.64 \pm 0.06$ |
| 75 | $0.26 \pm 0.05$ | $0.74 \pm 0.07$ | $0.74 \pm 0.07$ | $0.74 \pm 0.07$ | $\mathbf{0.76 \pm 0.07}$ | $\mathbf{0.76 \pm 0.07}$ | $0.74 \pm 0.06$ | $\mathbf{0.76 \pm 0.07}$ |
| 100 | $0.32 \pm 0.05$ | $0.78 \pm 0.04$ | $0.77 \pm 0.04$ | $0.78 \pm 0.04$ | $0.80 \pm 0.04$ | $0.79 \pm 0.04$ | $0.83 \pm 0.04$ | $\mathbf{0.85 \pm 0.04}$ |
| *Runtime (in seconds)* | | | | | | | | |
| 0 | $\mathbf{7.3 \pm 3.0}$ | $70.5 \pm 8.2$ | $70.8 \pm 8.3$ | $55.8 \pm 8.1$ | NA | NA | $8.8 \pm 4.3$ | $8653.5 \pm 4592.1$ |
| 25 | $8.1 \pm 4.9$ | $68.0 \pm 8.2$ | $68.0 \pm 8.2$ | $51.8 \pm 7.8$ | NA | NA | $\mathbf{6.6 \pm 3.3}$ | $5312.6 \pm 2485.7$ |
| 50 | $7.5 \pm 4.1$ | $67.7 \pm 8.2$ | $67.3 \pm 7.7$ | $48.8 \pm 8.2$ | NA | NA | $\mathbf{5.9 \pm 2.5}$ | $5097.6 \pm 2743.6$ |
| 75 | $7.2 \pm 2.9$ | $66.4 \pm 7.1$ | $66.6 \pm 7.2$ | $47.2 \pm 6.7$ | NA | NA | $\mathbf{4.1 \pm 1.5}$ | $3326.6 \pm 1549.0$ |
| 100 | $6.2 \pm 0.8$ | $64.6 \pm 5.6$ | $64.6 \pm 5.3$ | $41.4 \pm 5.4$ | NA | NA | $\mathbf{3.5 \pm 0.9}$ | $3045.6 \pm 1588.5$ |

#### 3.3 Supplemental results on KP-Tracer data sets

Table S8: **Runtime (in seconds) for KP-Tracer tumors.** Runtimes given for pipelines 1b,2b,1c,2c are the time to run LAML. The runtime is set to (x) 86400.0 when LAML did not complete. The runtime for StarCDP does not include the time to run PAUP* to compute the constrained search space. The runtime to compute a strict consensus tree of the best scoring trees found by PAUP* is not given (as it is very fast). The colors correspond to the pipeline/method that perform best in terms of the number of migrations and re-seeding events.

| Tumor Data | Analysis Pipeline | Startle NNI (C++) | StarCDP Bias | StarCDP Rand | StarCDP SC | PAUP* | PAUP* SC |
|---|---|---|---|---|---|---|---|
| 3724_NT_All | 1a | 527.0 | 651.4 | 651.4 | 651.4 | 3606.0 | – |
| | 2a | 299.1 | 406.8 | 406.8 | 406.8 | 14302.0 | – |
| | 1b | 868.9 | 1499.6 | 2937.2 | 29305.0 | **1406.9** | 28659.0 |
| | 2b | 28589.0 | 2857.3 | 1187.2 | 29283.0 | 28916.0 | 34526.0 |
| | 1c | (x) 86400.0 | (x) 86400.0 | 44779.0 | (x) 86400.0 | **3152.6** | (x) **86400.0** |
| | 2c | (x) 86400.0 | 1731.3 | 1677.3 | (x) 86400.0 | 5587.0 | (x) 86400.0 |
| 3513_NT_T1_Fam | 1a | 28.7 | 5.5 | 5.5 | 5.5 | 4.3 | – |
| | 2a | 4.9 | 2.9 | 2.9 | 2.9 | 34.2 | – |
| | 1b | 194.8 | 896.8 | 4517.0 | 1202.0 | **880.6** | 923.1 |
| | 2b | 1179.9 | 335.6 | 336.5 | 1236.8 | 635.7 | 1997.1 |
| | 1c | 786.4 | 1974.9 | 2630.9 | **2076.7** | **1056.4** | 1068.6 |
| | 2c | 438.7 | 693.2 | 205.7 | 852.8 | 705.1 | 1394.8 |
| 3515_LKB1_T1_FAM | 1a | 261.4 | **462.9** | 462.9 | 462.9 | 581.3 | – |
| | 2a | 200.4 | 351.7 | **351.7** | 351.7 | 597.5 | – |
| | 1b | 12779.0 | (x) 86400.0 | 69204.0 | (x) 86400.0 | (x) 86400.0 | (x) 86400.0 |
| | 2b | (x) 86400.0 | (x) 86400.0 | (x) 86400.0 | (x) 86400.0 | (x) 86400.0 | (x) 86400.0 |
| | 1c | (x) 86400.0 | (x) 86400.0 | (x) 86400.0 | (x) 86400.0 | (x) 86400.0 | (x) 86400.0 |
| | 2c | 77923.0 | (x) 86400.0 | (x) 86400.0 | (x) 86400.0 | 82301.0 | (x) 86400.0 |

Table S9: **SH parsimony scores for KP-Tracer tumors.**

| Tumor Data | Analysis Pipeline | Cassiopeia Hybrid | Startle NNI (C++) | StarCDP Bias | StarCDP Rand | StarCDP SC | PAUP* | PAUP* SC |
|---|---|---|---|---|---|---|---|---|
| 3724_NT_All | 1a | 4824.7 | 5493.1 | 4494.0 | 4494.0 | 4549.4 | 4494.0 | 4618.8 |
| | 2a | 4759.1 | 5104.5 | **4462.8** | **4462.8** | 4779.4 | **4462.8** | 5020.7 |
| | 1b | NA | 5493.1 | 4494.0 | 4494.0 | 4516.5 | **4494.0** | 4522.5 |
| | 2b | NA | 5103.2 | **4462.8** | **4462.8** | 4603.6 | **4462.8** | 4738.3 |
| | 1c | NA | 5493.1 | 4500.9 | 4494.0 | 4513.5 | **4494.0** | **4530.3** |
| | 2c | NA | 5110.3 | **4462.8** | **4462.8** | 4542.5 | **4462.8** | 4580.2 |
| 3513_NT_T1_Fam | 1a | 317.6 | 317.5 | 312.6 | 312.6 | 637.4 | 312.6 | 1079.5 |
| | 2a | 317.2 | 316.2 | **306.9** | **306.9** | 381.6 | **306.9** | 815.4 |
| | 1b | NA | 317.5 | 312.6 | 312.6 | 340.5 | **312.6** | 313.8 |
| | 2b | NA | 316.2 | 306.9 | 306.9 | 310.6 | 306.9 | 318.3 |
| | 1c | NA | 317.6 | 313.1 | 313.1 | **318.3** | **312.8** | 317.1 |
| | 2c | NA | 316.3 | 309.6 | 308.4 | 318.4 | 308.0 | 313.1 |
| 3515_LKB1_T1_Fam | 1a | 6042.5 | 5989.7 | **5365.2** | 5365.2 | 5409.1 | 5365.2 | 5422.2 |
| | 2a | 5989.1 | 5833.3 | **5352.1** | **5352.1** | 5362.4 | **5352.1** | 5373.3 |
| | 1b | NA | 5978.7 | 5365.2 | 5365.2 | 5401.0 | 5365.2 | 5404.4 |
| | 2b | NA | 5832.9 | **5352.1** | **5352.1** | 5360.5 | **5352.1** | 5363.1 |
| | 1c | NA | 6015.6 | 5365.2 | 5418.1 | 5407.3 | 5365.2 | 5415.7 |
| | 2c | NA | 5837.9 | **5352.1** | **5352.1** | 5360.5 | **5352.1** | 5376.1 |

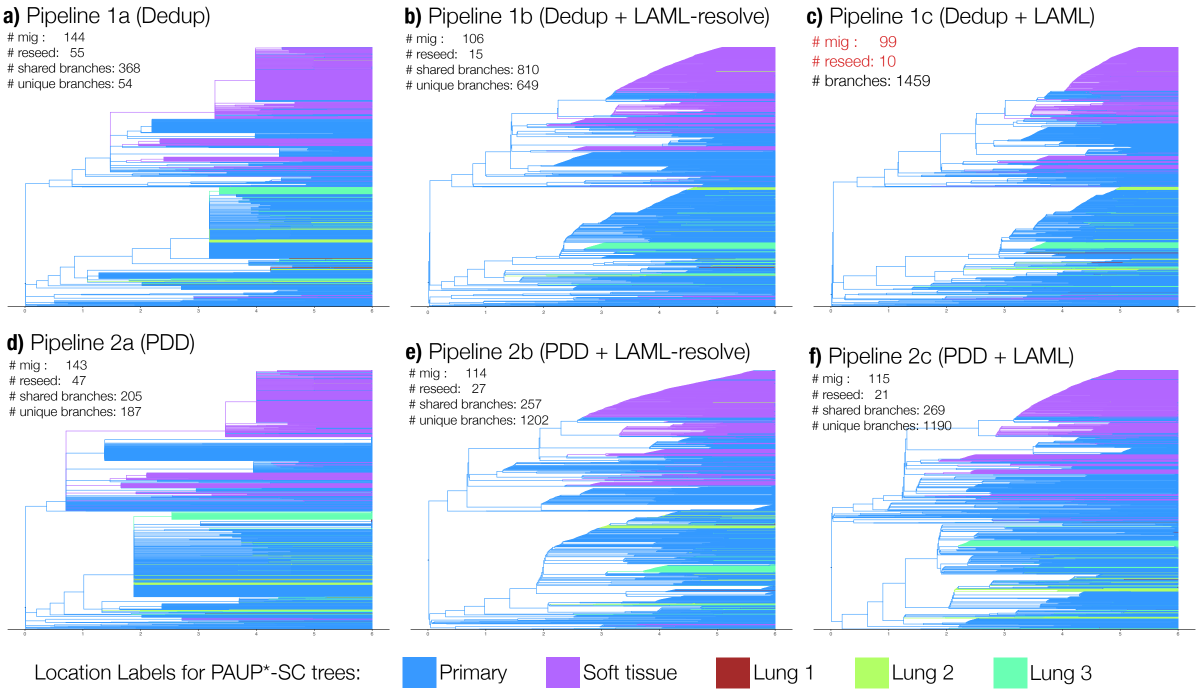

Figure S13: **Results of applying PAUP*-SC on KP-Tracer data within six pipelines.** PAUP*-SC run within pipeline 1c achieves the lowest number of migrations (99) and reseedings (10) of the six approaches. The result is a binary tree with 1459 internal branches. Only 55% of these internal branches are shared by the next best approach: pipeline 1b. Note that branch lengths were estimated with LAML, although this might not work very well for pipelines 1a and 2a as the trees are non-binary.

Table S10: **Number of migrations inferred for KP-Tracer tumor 3724_NT_All.** This metastasis family has a primary tumor, with three lung mets as well as one soft tissue met; see Figure 7 in [12]. It was previously analyzed by Mai *et al.* for the LAML [6] study. Mai *et al.* reported 119 migrations (56 reseedings) for the Startle-NNI (Python) tree, 99 migrations (42 reseedings) for the LAML tree, which starts its search from the Startle-NNI (Python) tree, 136 migrations (68 reseedings) for the Cassiopeia-Hybrid tree. Our analysis also found 136 migrations (68 migrations) for pipeline 1a, although simply pruning and unpruning cells with missing data (as part of pipeline 2a) drops the number of migrations and reseedings down to 114 and 32, respectively, lowering the number of migrations and reseedings below those reported for the Startle-NNI (Python) tree.

| Migration Type | Analysis Pipeline | Cassiopeia Hybrid | Startle NNI (C++) | StarCDP Bias | StarCDP Rand | StarCDP SC | PAUP* | PAUP* SC |
|---|---|---|---|---|---|---|---|---|
| migration | 1a | 136 | **113** | 123 | 127 | 133 | 143 | 144 |
| | 2a | 114 | 114 | **112** | 115 | 120 | 134 | 143 |
| | 1b | NA | 114 | 125 | 130 | 132 | **102** | 106 |
| | 2b | NA | 116 | 112 | 114 | 110 | 110 | 114 |
| | 1c | NA | 113 | 128 | 127 | 125 | **102** | **99** |
| | 2c | NA | 118 | 111 | 113 | 114 | 105 | 115 |
| reseeding | 1a | 68 | **32** | 53 | 55 | 57 | 56 | 55 |
| | 2a | 53 | 38 | **32** | 37 | 42 | 43 | 47 |
| | 1b | NA | 36 | 53 | 53 | 56 | **9** | 15 |
| | 2b | NA | 31 | 28 | 31 | 34 | 24 | 27 |
| | 1c | NA | 32 | 51 | 55 | 55 | **9** | **10** |
| | 2c | NA | 33 | 26 | 32 | 34 | 20 | 21 |
| Primary1↦Lung3 | 1a | 7 | 22 | 19 | 19 | 21 | 19 | 21 |
| | 2a | 7 | 14 | 21 | 21 | 21 | 21 | 22 |
| | 1b | NA | 22 | 19 | 19 | 18 | 21 | 21 |
| | 2b | NA | 16 | 22 | 22 | 22 | 23 | 22 |
| | 1c | NA | 22 | 21 | 19 | 19 | 21 | 20 |
| | 2c | NA | 16 | 22 | 22 | 22 | 23 | 27 |
| Lung2↦Lung1 | 1a | 0 | 0 | 0 | 0 | 0 | 0 | 0 |
| | 2a | 0 | 0 | 0 | 0 | 0 | 0 | 0 |
| | 1b | NA | 0 | 0 | 0 | 0 | 0 | 0 |
| | 2b | NA | 0 | 0 | 0 | 0 | 0 | 0 |
| | 1c | NA | 0 | 0 | 0 | 0 | 0 | 1 |
| | 2c | NA | 0 | 0 | 0 | 0 | 0 | 0 |
| Primary1↦Soft1 | 1a | 15 | 22 | 25 | 26 | 25 | 25 | 25 |
| | 2a | 9 | 26 | 32 | 30 | 27 | 26 | 30 |
| | 1b | NA | 21 | 25 | 28 | 28 | 48 | 46 |
| | 2b | NA | 30 | 35 | 33 | 28 | 35 | 36 |
| | 1c | NA | 22 | 25 | 26 | 24 | 48 | 45 |
| | 2c | NA | 27 | 36 | 32 | 30 | 36 | 37 |
| Lung1↦Lung2 | 1a | 0 | 0 | 1 | 1 | 1 | 1 | 1 |
| | 2a | 0 | 0 | 0 | 0 | 0 | 0 | 0 |
| | 1b | NA | 0 | 1 | 1 | 1 | 1 | 1 |
| | 2b | NA | 0 | 0 | 0 | 0 | 0 | 0 |
| | 1c | NA | 0 | 1 | 1 | 0 | 1 | 0 |
| | 2c | NA | 0 | 0 | 0 | 0 | 0 | 0 |
| Soft1↦Primary1 | 1a | 57 | 27 | 46 | 47 | 51 | 51 | 51 |
| | 2a | 45 | 27 | 26 | 30 | 36 | 38 | 42 |
| | 1b | NA | 30 | 46 | 45 | 47 | **9** | 14 |
| | 2b | NA | 25 | 23 | 26 | 29 | 20 | 21 |
| | 1c | NA | 27 | 46 | 47 | 46 | **9** | **10** |
| | 2c | NA | 26 | 22 | 27 | 29 | 17 | 18 |
| Lung3↦Primary1 | 1a | 11 | 2 | 4 | 5 | 4 | 5 | 4 |
| | 2a | 8 | 7 | 5 | 5 | 5 | 5 | 5 |
| | 1b | NA | 2 | 4 | 5 | 4 | 0 | 1 |
| | 2b | NA | 4 | 3 | 3 | 3 | 1 | 3 |
| | 1c | NA | 2 | 3 | 5 | 4 | 0 | 0 |
| | 2c | NA | 6 | 3 | 3 | 4 | 0 | 0 |
| Lung2↦Lung3 | 1a | 0 | 0 | 0 | 0 | 0 | 0 | 0 |
| | 2a | 0 | 0 | 0 | 0 | 0 | 0 | 0 |
| | 1b | NA | 0 | 0 | 0 | 0 | 0 | 0 |
| | 2b | NA | 0 | 0 | 0 | 0 | 1 | 1 |
| | 1c | NA | 0 | 0 | 0 | 0 | 0 | 0 |
| | 2c | NA | 0 | 0 | 0 | 0 | 1 | 1 |
| Soft1↦Lung2 | 1a | 1 | 1 | 1 | 1 | 1 | 1 | 1 |
| | 2a | 1 | 1 | 1 | 1 | 1 | 1 | 1 |
| | 1b | NA | 1 | 1 | 1 | 1 | 1 | 1 |
| | 2b | NA | 1 | 1 | 1 | 1 | 1 | 1 |
| | 1c | NA | 1 | 1 | 1 | 1 | 1 | 1 |
| | 2c | NA | 1 | 1 | 1 | 1 | 1 | 1 |
| Lung2↦Primary1 | 1a | 0 | 3 | 3 | 3 | 2 | 0 | 0 |
| | 2a | 0 | 4 | 1 | 2 | 1 | 0 | 0 |
| | 1b | NA | 4 | 3 | 3 | 5 | 0 | 0 |
| | 2b | NA | 2 | 2 | 2 | 2 | 3 | 3 |
| | 1c | NA | 3 | 2 | 3 | 5 | 0 | 0 |
| | 2c | NA | 1 | 1 | 2 | 1 | 3 | 3 |
| Primary1↦Lung1 | 1a | 9 | 8 | 8 | 8 | 8 | 8 | 8 |
| | 2a | 9 | 9 | 9 | 9 | 9 | 9 | 9 |
| | 1b | NA | 8 | 8 | 8 | 8 | 8 | 8 |
| | 2b | NA | 9 | 9 | 9 | 9 | 9 | 9 |
| | 1c | NA | 8 | 8 | 8 | 8 | 8 | 7 |
| | 2c | NA | 8 | 9 | 9 | 8 | 9 | 9 |
| Primary1↦Lung2 | 1a | 36 | 28 | 16 | 17 | 20 | 33 | 33 |
| | 2a | 35 | 26 | 17 | 17 | 20 | 34 | 34 |
| | 1b | NA | 26 | 18 | 20 | 20 | 14 | 14 |
| | 2b | NA | 29 | 17 | 18 | 16 | 17 | 18 |
| | 1c | NA | 28 | 21 | 17 | 18 | 14 | 15 |
| | 2c | NA | 33 | 17 | 17 | 19 | 15 | 19 |

Table S11: **Number of migrations inferred for KP-Tracer tumor 3513_NT_T1_Fam.** This data set was analyzed in the Startle study but it is somewhat not detailed which leaf set the trees are on, so we don't make a comparison.

| Migration Type | Analysis Pipeline | Cassiopeia Hybrid | Startle NNI (C++) | StarCDP Bias | StarCDP Rand | StarCDP SC | PAUP* | PAUP* SC |
|---|---|---|---|---|---|---|---|---|
| migration | 1a | 40 | **28** | 33 | 34 | 40 | 36 | 40 |
| | 2a | 36 | 31 | 27 | **25** | 34 | 36 | 36 |
| | 1b | NA | 30 | 34 | 33 | 20 | **20** | 21 |
| | 2b | NA | 27 | 25 | 28 | 30 | 30 | 29 |
| | 1c | NA | 28 | 35 | 35 | **23** | **19** | 24 |
| | 2c | NA | 31 | 29 | 25 | 26 | 28 | 33 |
| reseeding | 1a | 0 | **7** | 7 | 3 | 0 | 0 | 0 |
| | 2a | 1 | 5 | 3 | **2** | 1 | 1 | 1 |
| | 1b | NA | 1 | 3 | 0 | 6 | **2** | 1 |
| | 2b | NA | 2 | 2 | 3 | 2 | 6 | 5 |
| | 1c | NA | 1 | 7 | 1 | **0** | **3** | 4 |
| | 2c | NA | 5 | 2 | 2 | 5 | 2 | 4 |
| Primary1↦Kidney1 | 1a | 2 | 1 | 2 | 1 | 2 | 2 | 2 |
| | 2a | 2 | 2 | 2 | 1 | 2 | 2 | 2 |
| | 1b | NA | 1 | 2 | 1 | 1 | 2 | 1 |
| | 2b | NA | 1 | 2 | 2 | 1 | 1 | 1 |
| | 1c | NA | 1 | 1 | 1 | 2 | 1 | 1 |
| | 2c | NA | 2 | 2 | 2 | 2 | 1 | 2 |
| Kidney1↦Lymph-tissue1 | 1a | 0 | 1 | 0 | 0 | 0 | 0 | 0 |
| | 2a | 0 | 0 | 0 | 0 | 0 | 0 | 0 |
| | 1b | NA | 0 | 0 | 0 | 1 | 0 | 0 |
| | 2b | NA | 0 | 0 | 0 | 0 | 0 | 0 |
| | 1c | NA | 0 | 0 | 0 | 0 | 0 | 0 |
| | 2c | NA | 0 | 0 | 0 | 0 | 0 | 0 |
| Lymph-tissue1↦Primary1 | 1a | 0 | 7 | 7 | 3 | 0 | 0 | 0 |
| | 2a | 1 | 5 | 3 | 2 | 1 | 1 | 1 |
| | 1b | NA | 1 | 3 | 0 | 6 | 2 | 1 |
| | 2b | NA | 2 | 2 | 3 | 2 | 6 | 5 |
| | 1c | NA | 1 | 7 | 1 | 0 | 3 | 4 |
| | 2c | NA | 5 | 2 | 2 | 5 | 2 | 4 |
| Primary1↦Lymph1 | 1a | 38 | 19 | 24 | 30 | 38 | 34 | 38 |
| | 2a | 33 | 24 | 22 | 22 | 31 | 33 | 33 |
| | 1b | NA | 28 | 29 | 32 | 12 | 16 | 19 |
| | 2b | NA | 24 | 21 | 23 | 27 | 23 | 23 |
| | 1c | NA | 26 | 27 | 33 | 21 | 15 | 19 |
| | 2c | NA | 24 | 25 | 21 | 19 | 25 | 27 |

Table S12: **Number of migrations inferred for KP-Tracer tumor 3515_LKB1_T1_FAM.** This metastasis family has a primary tumor, with two lung mets and three kidney mets, as well as one lymph met; see Figure 7 in [12]

| Migration Type | Analysis Pipeline | Cassiopeia Hybrid | Startle NNI (C++) | StarCDP Bias | StarCDP Rand | StarCDP SC | PAUP* | PAUP* SC |
|---|---|---|---|---|---|---|---|---|
| migration | 1a | 171 | 170 | **145** | 147 | 162 | 154 | 158 |
| | 2a | 175 | 161 | 154 | 156 | 165 | 158 | 161 |
| | 1b | NA | 168 | 147 | **146** | 149 | 150 | 149 |
| | 2b | NA | 159 | 152 | 153 | 150 | 145 | 151 |
| | 1c | NA | 167 | 147 | 145 | 146 | 146 | 148 |
| | 2c | NA | 153 | 150 | 153 | 151 | 153 | 153 |
| reseeding | 1a | 4 | 12 | **3** | 3 | 7 | 5 | 7 |
| | 2a | 4 | 5 | 4 | 4 | 3 | 3 | 3 |
| | 1b | NA | 9 | 2 | **2** | 4 | 8 | 7 |
| | 2b | NA | 8 | 3 | 3 | 4 | 4 | 4 |
| | 1c | NA | 11 | 6 | 3 | 4 | 8 | 7 |
| | 2c | NA | 5 | 4 | 5 | 4 | 4 | 4 |
| Lung1↦Kidney3 | 1a | 1 | 0 | 1 | 1 | 1 | 1 | 1 |
| | 2a | 2 | 0 | 1 | 2 | 1 | 1 | 1 |
| | 1b | NA | 0 | 1 | 1 | 1 | 1 | 1 |
| | 2b | NA | 0 | 1 | 2 | 1 | 1 | 1 |
| | 1c | NA | 0 | 1 | 1 | 1 | 1 | 1 |
| | 2c | NA | 0 | 1 | 2 | 1 | 1 | 1 |
| Primary1↦Lung2 | 1a | 1 | 1 | 1 | 1 | 1 | 1 | 1 |
| | 2a | 1 | 1 | 1 | 1 | 1 | 1 | 1 |
| | 1b | NA | 1 | 1 | 1 | 1 | 1 | 1 |
| | 2b | NA | 1 | 1 | 1 | 1 | 1 | 1 |
| | 1c | NA | 1 | 1 | 1 | 1 | 1 | 1 |
| | 2c | NA | 1 | 1 | 1 | 1 | 1 | 1 |
| Kidney3↦Kidney2 | 1a | 19 | 49 | 29 | 34 | 46 | 35 | 32 |
| | 2a | 23 | 46 | 20 | 22 | 46 | 31 | 31 |
| | 1b | NA | 42 | 43 | 36 | 36 | 25 | 20 |
| | 2b | NA | 42 | 18 | 16 | 17 | 6 | 15 |
| | 1c | NA | 51 | 24 | 18 | 26 | 36 | 28 |
| | 2c | NA | 30 | 16 | 15 | 18 | 15 | 17 |
| Lung1↦Kidney2 | 1a | 1 | 2 | 0 | 0 | 0 | 0 | 0 |
| | 2a | 2 | 0 | 0 | 0 | 0 | 0 | 0 |
| | 1b | NA | 1 | 0 | 0 | 0 | 0 | 0 |
| | 2b | NA | 0 | 0 | 0 | 0 | 0 | 0 |
| | 1c | NA | 1 | 0 | 0 | 0 | 0 | 0 |
| | 2c | NA | 2 | 0 | 0 | 0 | 0 | 0 |
| Primary1↦Kidney1 | 1a | 2 | 2 | 2 | 1 | 1 | 2 | 1 |
| | 2a | 2 | 4 | 2 | 2 | 2 | 2 | 2 |
| | 1b | NA | 2 | 2 | 2 | 1 | 2 | 3 |
| | 2b | NA | 4 | 2 | 2 | 2 | 2 | 2 |
| | 1c | NA | 2 | 2 | 1 | 1 | 2 | 2 |
| | 2c | NA | 4 | 2 | 2 | 2 | 2 | 2 |
| Kidney1↦Kidney3 | 1a | 2 | 0 | 1 | 2 | 0 | 0 | 0 |
| | 2a | 2 | 0 | 1 | 2 | 1 | 1 | 1 |
| | 1b | NA | 1 | 1 | 1 | 3 | 3 | 2 |
| | 2b | NA | 0 | 2 | 2 | 3 | 2 | 2 |
| | 1c | NA | 1 | 1 | 3 | 2 | 4 | 4 |
| | 2c | NA | 1 | 2 | 2 | 2 | 1 | 1 |
| Lung1↦Primary1 | 1a | 0 | 2 | 0 | 0 | 0 | 0 | 0 |
| | 2a | 2 | 0 | 0 | 0 | 0 | 0 | 0 |
| | 1b | NA | 2 | 0 | 0 | 0 | 0 | 0 |
| | 2b | NA | 0 | 0 | 0 | 0 | 0 | 0 |
| | 1c | NA | 3 | 0 | 0 | 0 | 0 | 0 |
| | 2c | NA | 0 | 0 | 0 | 0 | 0 | 0 |
| Primary1↦Lung1 | 1a | 3 | 3 | 2 | 2 | 2 | 2 | 2 |
| | 2a | 2 | 5 | 2 | 2 | 2 | 2 | 2 |
| | 1b | NA | 4 | 2 | 2 | 2 | 2 | 2 |
| | 2b | NA | 4 | 2 | 2 | 2 | 2 | 2 |
| | 1c | NA | 2 | 2 | 2 | 2 | 2 | 2 |
| | 2c | NA | 3 | 2 | 2 | 2 | 2 | 2 |
| Primary1↦Lymph1 | 1a | 20 | 16 | 17 | 17 | 18 | 18 | 18 |
| | 2a | 20 | 17 | 18 | 17 | 20 | 20 | 20 |
| | 1b | NA | 20 | 20 | 18 | 17 | 11 | 10 |
| | 2b | NA | 18 | 20 | 19 | 18 | 17 | 17 |
| | 1c | NA | 19 | 15 | 16 | 17 | 10 | 10 |
| | 2c | NA | 20 | 18 | 17 | 18 | 18 | 18 |
| Lymph1↦Primary1 | 1a | 0 | 3 | 1 | 2 | 2 | 2 | 2 |
| | 2a | 1 | 2 | 2 | 2 | 1 | 1 | 1 |
| | 1b | NA | 0 | 0 | 1 | 2 | 5 | 5 |
| | 2b | NA | 1 | 1 | 1 | 2 | 2 | 2 |
| | 1c | NA | 1 | 4 | 2 | 2 | 5 | 5 |
| | 2c | NA | 0 | 2 | 2 | 2 | 2 | 2 |

Table S13: **Number of migrations inferred for KP-Tracer tumor 3515_LKB1_T1_FAM (continued).**

| Migration Type | Analysis Pipeline | Cassiopeia Hybrid | Startle NNI (C++) | StarCDP Bias | StarCDP Rand | StarCDP SC | PAUP* | PAUP* SC |
|---|---|---|---|---|---|---|---|---|
| Primary1↦Kidney2 | 1a | 10 | 7 | 5 | 7 | 5 | 6 | 5 |
| | 2a | 8 | 8 | 8 | 7 | 8 | 7 | 8 |
| | 1b | NA | 8 | 5 | 7 | 7 | 6 | 7 |
| | 2b | NA | 5 | 8 | 7 | 8 | 7 | 8 |
| | 1c | NA | 9 | 5 | 7 | 7 | 5 | 7 |
| | 2c | NA | 6 | 8 | 6 | 8 | 7 | 8 |
| Kidney3↦Lung1 | 1a | 2 | 1 | 0 | 0 | 0 | 0 | 0 |
| | 2a | 1 | 1 | 0 | 0 | 1 | 0 | 1 |
| | 1b | NA | 0 | 0 | 0 | 0 | 0 | 0 |
| | 2b | NA | 2 | 0 | 0 | 0 | 0 | 0 |
| | 1c | NA | 1 | 0 | 0 | 0 | 0 | 0 |
| | 2c | NA | 3 | 0 | 0 | 0 | 0 | 0 |
| Kidney2↦Kidney3 | 1a | 88 | 56 | 65 | 57 | 63 | 64 | 73 |
| | 2a | 88 | 52 | 78 | 78 | 61 | 72 | 72 |
| | 1b | NA | 58 | 51 | 54 | 55 | 71 | 75 |
| | 2b | NA | 58 | 76 | 80 | 75 | 85 | 80 |
| | 1c | NA | 48 | 71 | 76 | 65 | 57 | 64 |
| | 2c | NA | 61 | 76 | 83 | 75 | 84 | 80 |
| Kidney2↦Primary-1 | 1a | 0 | 0 | 1 | 1 | 1 | 0 | 1 |
| | 2a | 0 | 2 | 0 | 0 | 0 | 0 | 0 |
| | 1b | NA | 1 | 1 | 0 | 1 | 0 | 1 |
| | 2b | NA | 2 | 0 | 0 | 0 | 0 | 0 |
| | 1c | NA | 0 | 1 | 1 | 1 | 1 | 1 |
| | 2c | NA | 3 | 0 | 1 | 0 | 0 | 0 |
| Kidney1↦Kidney2 | 1a | 0 | 1 | 0 | 0 | 0 | 0 | 0 |
| | 2a | 0 | 0 | 0 | 0 | 0 | 0 | 0 |
| | 1b | NA | 2 | 0 | 0 | 2 | 0 | 0 |
| | 2b | NA | 0 | 0 | 0 | 0 | 0 | 0 |
| | 1c | NA | 1 | 0 | 0 | 0 | 0 | 0 |
| | 2c | NA | 0 | 1 | 0 | 1 | 0 | 0 |
| Lung2↦Kidney1 | 1a | 3 | 5 | 2 | 3 | 3 | 3 | 3 |
| | 2a | 3 | 3 | 3 | 3 | 3 | 3 | 3 |
| | 1b | NA | 5 | 2 | 3 | 3 | 3 | 3 |
| | 2b | NA | 3 | 3 | 3 | 3 | 3 | 3 |
| | 1c | NA | 5 | 2 | 3 | 3 | 3 | 3 |
| | 2c | NA | 3 | 3 | 3 | 3 | 3 | 3 |
| Primary1↦Kidney3 | 1a | 6 | 4 | 8 | 9 | 5 | 8 | 5 |
| | 2a | 9 | 9 | 8 | 8 | 8 | 7 | 8 |
| | 1b | NA | 4 | 8 | 10 | 7 | 8 | 10 |
| | 2b | NA | 4 | 8 | 8 | 8 | 7 | 8 |
| | 1c | NA | 4 | 8 | 4 | 7 | 8 | 10 |
| | 2c | NA | 4 | 8 | 7 | 8 | 7 | 8 |
| Kidney2↦Kidney1 | 1a | 7 | 5 | 6 | 7 | 8 | 7 | 8 |
| | 2a | 7 | 5 | 7 | 7 | 6 | 6 | 6 |
| | 1b | NA | 6 | 5 | 6 | 7 | 7 | 7 |
| | 2b | NA | 6 | 7 | 7 | 8 | 8 | 6 |
| | 1c | NA | 6 | 6 | 7 | 8 | 7 | 7 |
| | 2c | NA | 7 | 6 | 8 | 7 | 7 | 6 |
| Kidney3↦Kidney1 | 1a | 2 | 6 | 3 | 3 | 2 | 2 | 2 |
| | 2a | 1 | 5 | 1 | 1 | 2 | 2 | 2 |
| | 1b | NA | 5 | 4 | 3 | 3 | 2 | 1 |
| | 2b | NA | 4 | 1 | 1 | 0 | 0 | 2 |
| | 1c | NA | 5 | 3 | 3 | 2 | 2 | 2 |
| | 2c | NA | 3 | 2 | 0 | 1 | 1 | 2 |
| Kidney3↦Primary1 | 1a | 4 | 7 | 1 | 0 | 4 | 3 | 4 |
| | 2a | 1 | 1 | 2 | 2 | 2 | 2 | 2 |
| | 1b | NA | 6 | 1 | 1 | 1 | 3 | 1 |
| | 2b | NA | 5 | 2 | 2 | 2 | 2 | 2 |
| | 1c | NA | 7 | 1 | 0 | 1 | 2 | 1 |
| | 2c | NA | 2 | 2 | 2 | 2 | 2 | 2 |

Table S14: **Distance between CLTs for KP-Tracer tumor 3724_NT_All.** All trees are compared to the CLT estimated with **PAUP*-SC as part of pipeline 1c** because it achieved the second lowest number of reseedings (10 instead of 9) and the lowest number of migrations (99).

| Tree Comparison | Analysis Pipeline | Cassiopeia Hybrid | Startle NNI (C++) | StarCDP Bias | StarCDP Rand | StarCDP SC | PAUP* | **PAUP* SC** |
|---|---|---|---|---|---|---|---|---|
| TP (No contraction) | 1a | 149 | 70 | 424 | 440 | 374 | 379 | 368 |
| | 2a | 177 | 116 | 238 | 241 | 209 | 212 | 205 |
| | 1b | NA | 62 | 424 | 435 | 451 | 843 | 810 |
| | 2b | NA | 122 | 240 | 235 | 251 | 256 | 257 |
| | **1c** | NA | 70 | 454 | 441 | 490 | 843 | **1459** |
| | 2c | NA | 130 | 240 | 243 | 262 | 257 | 269 |
| FP (No contraction) | 1a | 268 | 1389 | 1035 | 1019 | 480 | 109 | 54 |
| | 2a | 249 | 1222 | 1100 | 1097 | 622 | 250 | 187 |
| | 1b | NA | 1397 | 1035 | 1024 | 1008 | 616 | 649 |
| | 2b | NA | 1337 | 1219 | 1224 | 1208 | 1203 | 1202 |
| | **1c** | NA | 1389 | 1005 | 1018 | 969 | 616 | **0** |
| | 2c | NA | 1329 | 1219 | 1216 | 1197 | 1202 | 1190 |
| FN (No contraction) | 1a | 1310 | 1389 | 1035 | 1019 | 1085 | 1080 | 1091 |
| | 2a | 1282 | 1343 | 1221 | 1218 | 1250 | 1247 | 1254 |
| | 1b | NA | 1397 | 1035 | 1024 | 1008 | 616 | 649 |
| | 2b | NA | 1337 | 1219 | 1224 | 1208 | 1203 | 1202 |
| | **1c** | NA | 1389 | 1005 | 1018 | 969 | 616 | **0** |
| | 2c | NA | 1329 | 1219 | 1216 | 1197 | 1202 | 1190 |
| RF (No contraction) | 1a | 1578 | 2778 | 2070 | 2038 | 1565 | 1189 | 1145 |
| | 2a | 1531 | 2565 | 2321 | 2315 | 1872 | 1497 | 1441 |
| | 1b | NA | 2794 | 2070 | 2048 | 2016 | 1232 | 1298 |
| | 2b | NA | 2674 | 2438 | 2448 | 2416 | 2406 | 2404 |
| | **1c** | NA | 2778 | 2010 | 2036 | 1938 | 1232 | **0** |
| | 2c | NA | 2658 | 2438 | 2432 | 2394 | 2404 | 2380 |
| TP (SH contraction) | 1a | 136 | 41 | 356 | 362 | 360 | 362 | 357 |
| | 2a | 161 | 78 | 194 | 192 | 192 | 194 | 189 |
| | 1b | NA | 41 | 356 | 362 | 359 | 362 | 366 |
| | 2b | NA | 79 | 194 | 192 | 192 | 194 | 190 |
| | **1c** | NA | 41 | 348 | 362 | 351 | 362 | **415** |
| | 2c | NA | 78 | 194 | 192 | 193 | 194 | 192 |
| FP (SH contraction) | 1a | 230 | 254 | 76 | 76 | 51 | 74 | 38 |
| | 2a | 203 | 259 | 198 | 208 | 168 | 201 | 161 |
| | 1b | NA | 256 | 76 | 76 | 76 | 75 | 71 |
| | 2b | NA | 261 | 198 | 208 | 216 | 201 | 256 |
| | **1c** | NA | 254 | 73 | 75 | 56 | 75 | **0** |
| | 2c | NA | 248 | 198 | 208 | 203 | 201 | 226 |
| FN (SH contraction) | 1a | 279 | 374 | 59 | 53 | 55 | 53 | 58 |
| | 2a | 254 | 337 | 221 | 223 | 223 | 221 | 226 |
| | 1b | NA | 374 | 59 | 53 | 56 | 53 | 49 |
| | 2b | NA | 336 | 221 | 223 | 223 | 221 | 225 |
| | **1c** | NA | 374 | 67 | 53 | 64 | 53 | **0** |
| | 2c | NA | 337 | 221 | 223 | 222 | 221 | 223 |
| RF (SH contraction) | 1a | 509 | 628 | 135 | 129 | 106 | 127 | 96 |
| | 2a | 457 | 596 | 419 | 431 | 391 | 422 | 387 |
| | 1b | NA | 630 | 135 | 129 | 132 | 128 | 120 |
| | 2b | NA | 597 | 419 | 431 | 439 | 422 | 481 |
| | **1c** | NA | 628 | 140 | 128 | 120 | 128 | **0** |
| | 2c | NA | 585 | 419 | 431 | 425 | 422 | 449 |

Table S15: **Distance between CLTs for KP-Tracer tumor 3513_NT_T1_Fam.** All trees are compared to the CLT estimated with Star-CDP as part of pipeline 1c because it achieved the lowest number of reseedings (0) and a reasonable number of migrations (23 instead of 19 or 20).

| Tree Comparison | Analysis Pipeline | Cassiopeia Hybrid | Startle NNI (C++) | StarCDP Bias | StarCDP Rand | **StarCDP SC** | PAUP* | PAUP* SC |
|---|---|---|---|---|---|---|---|---|
| TP (No contraction) | 1a | 4 | 6 | 9 | 7 | 6 | 8 | 3 |
| | 2a | 5 | 6 | 7 | 7 | 6 | 7 | 6 |
| | 1b | NA | 5 | 9 | 7 | 14 | 10 | 10 |
| | 2b | NA | 5 | 9 | 8 | 8 | 9 | 10 |
| | **1c** | NA | 5 | 8 | 8 | **99** | 9 | 8 |
| | 2c | NA | 6 | 7 | 11 | 9 | 8 | 8 |
| FP (No contraction) | 1a | 17 | 93 | 90 | 92 | 0 | 21 | 0 |
| | 2a | 15 | 83 | 82 | 82 | 33 | 24 | 8 |
| | 1b | NA | 94 | 90 | 92 | 85 | 89 | 89 |
| | 2b | NA | 94 | 90 | 91 | 91 | 90 | 89 |
| | **1c** | NA | 94 | 91 | 91 | **0** | 90 | 91 |
| | 2c | NA | 93 | 92 | 88 | 90 | 91 | 91 |
| FN (No contraction) | 1a | 95 | 93 | 90 | 92 | 93 | 91 | 96 |
| | 2a | 94 | 93 | 92 | 92 | 93 | 92 | 93 |
| | 1b | NA | 94 | 90 | 92 | 85 | 89 | 89 |
| | 2b | NA | 94 | 90 | 91 | 91 | 90 | 89 |
| | **1c** | NA | 94 | 91 | 91 | **0** | 90 | 91 |
| | 2c | NA | 93 | 92 | 88 | 90 | 91 | 91 |
| RF (No contraction) | 1a | 112 | 186 | 180 | 184 | 93 | 112 | 96 |
| | 2a | 109 | 176 | 174 | 174 | 126 | 116 | 101 |
| | 1b | NA | 188 | 180 | 184 | 170 | 178 | 178 |
| | 2b | NA | 188 | 180 | 182 | 182 | 180 | 178 |
| | **1c** | NA | 188 | 182 | 182 | **0** | 180 | 182 |
| | 2c | NA | 186 | 184 | 176 | 180 | 182 | 182 |
| TP (SH contraction) | 1a | 4 | 5 | 7 | 6 | 6 | 7 | 3 |
| | 2a | 5 | 5 | 7 | 7 | 6 | 7 | 6 |
| | 1b | NA | 5 | 7 | 6 | 8 | 7 | 8 |
| | 2b | NA | 4 | 7 | 6 | 8 | 7 | 8 |
| | **1c** | NA | 5 | 8 | 7 | **28** | 7 | 5 |
| | 2c | NA | 6 | 7 | 8 | 7 | 7 | 8 |
| FP (SH contraction) | 1a | 12 | 12 | 14 | 17 | 0 | 17 | 0 |
| | 2a | 10 | 9 | 19 | 21 | 10 | 20 | 8 |
| | 1b | NA | 12 | 15 | 17 | 35 | 17 | 15 |
| | 2b | NA | 11 | 19 | 22 | 16 | 20 | 23 |
| | **1c** | NA | 11 | 13 | 12 | **0** | 15 | 15 |
| | 2c | NA | 9 | 16 | 17 | 19 | 18 | 20 |
| FN (SH contraction) | 1a | 24 | 23 | 21 | 22 | 22 | 21 | 25 |
| | 2a | 23 | 23 | 21 | 21 | 22 | 21 | 22 |
| | 1b | NA | 23 | 21 | 22 | 20 | 21 | 20 |
| | 2b | NA | 24 | 21 | 22 | 20 | 21 | 20 |
| | **1c** | NA | 23 | 20 | 21 | **0** | 21 | 23 |
| | 2c | NA | 22 | 21 | 20 | 21 | 21 | 20 |
| RF (SH contraction) | 1a | 36 | 35 | 35 | 39 | 22 | 38 | 25 |
| | 2a | 33 | 32 | 40 | 42 | 32 | 41 | 30 |
| | 1b | NA | 35 | 36 | 39 | 55 | 38 | 35 |
| | 2b | NA | 35 | 40 | 44 | 36 | 41 | 43 |
| | **1c** | NA | 34 | 33 | 33 | **0** | 36 | 38 |
| | 2c | NA | 31 | 37 | 37 | 40 | 39 | 40 |

Table S16: **Distance between CLTs for KP-Tracer tumor 3515_LKB1_T1_FAM.** All trees are compared to the CLT estimated using StarCDP-Bias as part of pipeline 1b because it achieved the lowest number of reseedings (2) and second lowest number of migrations (147 instead of 145/146).

| Tree Comparison | Analysis Pipeline | Cassiopeia Hybrid | Startle NNI (C++) | **StarCDP Bias** | StarCDP Rand | StarCDP SC | PAUP* | PAUP* SC |
|---|---|---|---|---|---|---|---|---|
| | 1a | 195 | 147 | 864 | 637 | 576 | 603 | 570 |
| | 2a | 207 | 185 | 354 | 334 | 304 | 309 | 294 |
| TP (No contraction) | **1b** | NA | 154 | **1011** | 592 | 680 | 628 | 595 |
| | 2b | NA | 186 | 356 | 333 | 341 | 326 | 316 |
| | 1c | NA | 150 | 848 | 434 | 696 | 641 | 609 |
| | 2c | NA | 193 | 358 | 335 | 344 | 326 | 317 |
| | 1a | 320 | 864 | 147 | 374 | 33 | 94 | 61 |
| | 2a | 322 | 769 | 600 | 620 | 262 | 373 | 330 |
| FP (No contraction) | **1b** | NA | 857 | **0** | 419 | 331 | 383 | 416 |
| | 2b | NA | 825 | 655 | 678 | 670 | 685 | 695 |
| | 1c | NA | 861 | 163 | 577 | 315 | 370 | 402 |
| | 2c | NA | 818 | 653 | 676 | 667 | 685 | 694 |
| | 1a | 816 | 864 | 147 | 374 | 435 | 408 | 441 |
| | 2a | 804 | 826 | 657 | 677 | 707 | 702 | 717 |
| FN (No contraction) | **1b** | NA | 857 | **0** | 419 | 331 | 383 | 416 |
| | 2b | NA | 825 | 655 | 678 | 670 | 685 | 695 |
| | 1c | NA | 861 | 163 | 577 | 315 | 370 | 402 |
| | 2c | NA | 818 | 653 | 676 | 667 | 685 | 694 |
| | 1a | 1136 | 1728 | 294 | 748 | 468 | 502 | 502 |
| | 2a | 1126 | 1595 | 1257 | 1297 | 969 | 1075 | 1047 |
| RF (No contraction) | **1b** | NA | 1714 | **0** | 838 | 662 | 766 | 832 |
| | 2b | NA | 1650 | 1310 | 1356 | 1340 | 1370 | 1390 |
| | 1c | NA | 1722 | 326 | 1154 | 630 | 740 | 804 |
| | 2c | NA | 1636 | 1306 | 1352 | 1334 | 1370 | 1388 |
| | 1a | 179 | 132 | 516 | 499 | 492 | 504 | 481 |
| | 2a | 194 | 170 | 292 | 292 | 290 | 290 | 284 |
| TP (SH contraction) | **1b** | NA | 135 | **521** | 498 | 492 | 504 | 486 |
| | 2b | NA | 170 | 292 | 292 | 294 | 290 | 288 |
| | 1c | NA | 131 | 511 | 379 | 490 | 501 | 484 |
| | 2c | NA | 167 | 292 | 292 | 294 | 290 | 289 |
| | 1a | 294 | 353 | 4 | 27 | 6 | 18 | 5 |
| | 2a | 278 | 324 | 202 | 208 | 190 | 205 | 186 |
| FP (SH contraction) | **1b** | NA | 354 | **0** | 29 | 21 | 18 | 20 |
| | 2b | NA | 326 | 202 | 208 | 194 | 207 | 196 |
| | 1c | NA | 329 | 6 | 91 | 17 | 19 | 17 |
| | 2c | NA | 296 | 202 | 207 | 194 | 203 | 191 |
| | 1a | 342 | 389 | 5 | 22 | 29 | 17 | 40 |
| | 2a | 327 | 351 | 229 | 229 | 231 | 231 | 237 |
| FN (SH contraction) | **1b** | NA | 386 | **0** | 23 | 29 | 17 | 35 |
| | 2b | NA | 351 | 229 | 229 | 227 | 231 | 233 |
| | 1c | NA | 390 | 10 | 142 | 31 | 20 | 37 |
| | 2c | NA | 354 | 229 | 229 | 227 | 231 | 232 |
| | 1a | 636 | 742 | 9 | 49 | 35 | 35 | 45 |
| | 2a | 605 | 675 | 431 | 437 | 421 | 436 | 423 |
| RF (SH contraction) | **1b** | NA | 740 | **0** | 52 | 50 | 35 | 55 |
| | 2b | NA | 677 | 431 | 437 | 421 | 438 | 429 |
| | 1c | NA | 719 | 16 | 233 | 48 | 39 | 54 |
| | 2c | NA | 650 | 431 | 436 | 421 | 434 | 423 |

[5] Daniel Kane and Terence Tao. A bound on partitioning clusters. *The Electronic Journal of Combinatorics*, 2017.

[6] Uyen Mai, Gillian Chu, and Benjamin J. Raphael. Maximum likelihood inference of time-scaled cell lineage trees with mixed-type missing data. *bioRxiv*, 2024.

[7] Uyen Mai, Gillian Chu, and Benjamin J. Raphael. Maximum likelihood inference of time-scaled cell lineage trees with mixed-type missing data. In Jian Ma, editor, *Research in Computational Molecular Biology*, page 360, Cham, 2024. Springer Nature Switzerland.

[8] D.F. Robinson and L.R. Foulds. Comparison of phylogenetic trees. *Mathematical Biosciences*, 53(1):131–147, 1981.

[9] Palash Sashittal, Henri Schmidt, Michelle Chan, and Benjamin J. Raphael. Startle: A star homoplasy approach for CRISPR-Cas9 lineage tracing. *Cell Systems*, 14:1113–1121, 2023.

[10] Palash Sashittal, Henri Schmidt, Michelle Chan, and Benjamin J. Raphael. Startle: a star homoplasy approach for crispr-cas9 lineage tracing. In Haixu Tang, editor, *Research in Computational Molecular Biology*, page 229, Cham, 2023. Springer Nature Switzerland.

[11] Tandy Warnow. *Computational Phylogenetics: An Introduction to Designing Methods for Phylogeny Estimation.* Cambridge University Press, Cambridge, United Kingdom, 2017.

[12] Dian Yang, Matthew G Jones, Santiago Naranjo, William M Rideout, 3rd, Kyung Hoi Joseph Min, Raymond Ho, Wei Wu, Joseph M Replogle, Jennifer L Page, Jeffrey J Quinn, Felix Horns, Xiaojie Qiu, Michael Z Chen, William A Freed-Pastor, Christopher S McGinnis, David M Patterson, Zev J Gartner, Eric D Chow, Trever G Bivona, Michelle M Chan, Nir Yosef, Tyler Jacks, and Jonathan S Weissman. Lineage tracing reveals the phylodynamics, plasticity, and paths of tumor evolution. *Cell*, 185(11):1905–1923.e25, 2022.

[13] Chao Zhang, Maryam Rabiee, Erfan Sayyari, and Siavash Mirarab. ASTRAL-III: Polynomial time species tree reconstruction from partially resolved gene trees. *BMC Bioinformatics*, 19(6):153, 2018.
